# Metabolic pathway analysis in the presence of biological constraints

**DOI:** 10.1101/2020.06.27.175455

**Authors:** Philippe Dague

## Abstract

Metabolic pathway analysis is a key method to study a metabolism in its steady state and the concept of elementary fluxes (EFs) plays a major role in the analysis of a network in terms of non-decomposable pathways. The supports of the EFs contain in particular those of the elementary flux modes (EFMs), which are the support-minimal pathways, and EFs coincide with EFMs when the only flux constraints are given by the irreversibility of certain reactions. Practical use of both EFMs and EFs has been hampered by the combinatorial explosion of their number in large, genomescale, systems. The EFs give the possible pathways in a steady state but the real pathways are limited by biological constraints, such as thermodynamic or, more generally, kinetic constraints and regulatory constraints from the genetic network. We provide results on the mathematical structure and geometrical characterization of the solution space in the presence of such biological constraints and revisit the concept of EFMs and EFs in this framework. We show that most of the results depend only on very general properties of compatibility of constraints with the sign function: either sign-invariance for regulatory constraints or sign-monotonicity (a stronger property) for thermodynamic and kinetic constraints. We show in particular that EFs for sign-monotone constraints are just those of the original EFs that satisfy the constraint and we show how to integrate their computation efficiently in the double description method, the most widely used method in the tools dedicated to EFMs computation.

## 1 Introduction

### 1.1 Metabolic networks

In order to ensure this paper is self-contained and has no prerequisite to be read, we summarize in this introduction the state of the art related to the subject and fix the notations adopted throughout the paper. The results quoted being known, are thus given without proof and the reader is invited to refer to [42, 46, 41, 25, 48, 30, 22], in addition to the references in the text, for the proofs or historical surveys.

A metabolic network is made up of a set of *r* biochemical enzymatic reactions. Each reaction consumes certain metabolites (called substrates of the reaction) and produces other metabolites (called products of the reaction). Each metabolite is assigned a coefficient in the reaction, its stoichiometric coefficient (counted negatively for substrates and positively for products). We distinguish internal from external metabolites w.r.t. the system under study (e.g., a bacterium, an eukaryotic cell), the reactions involving both internal and external metabolites being transfer reactions. If *m* is the number of internal metabolites (*r* > *m*), the network is thus given by its stoichiometric matrix 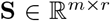, where coefficient **S**_*ji*_ is the stoichiometric coefficient of internal metabolite *j* in reaction *i* (positive if *j* is a product of reaction *i* and negative if it is a substrate). A state of the network at a given time *t* is given by the net rates (or fluxes) in each of its reactions at *t*, i.e., by a flux vector (or rate vector, or flux distribution) 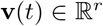. Denoting by 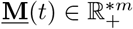 the vector of the concentrations of internal metabolites at *t*, the time evolution of the network is thus given by:

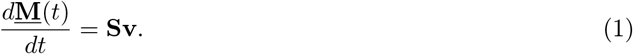

### 1.2 Steady-state behavior and flux subspace

We are interested in the steady-state behavior of the network. The steady-state assumption means that the concentrations of internal metabolites remain constant along time (no accumulation or reduction of internal metabolites inside the system, an approximation which is valid for short time periods, e.g., a few minutes) and leads thus to the fundamental equation:

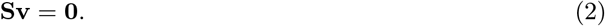

With only this assumption, the solution space *Sol*_**s**_, i.e., the space of all admissible flux vectors **v**, is thus the linear subspace *FS* of 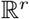 given by the kernel, or nullspace [38], of **S**:

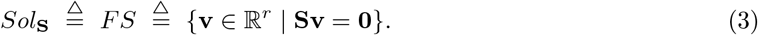

The dimension of the flux subspace is given by *dim*(*FS*) = *r* – *rank*(**S**) ≥ *r* – *m*. Often, possibly after a pre-processing to eliminate its linearly dependent rows, **S** is assumed to be of full rank and then *dim*(*FS*) = *r* – *m*.

The support of a vector 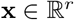 is defined by:

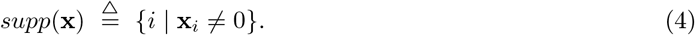

The support of a flux vector **v** has thus an important biological signification as it represents the reactions involved in the subnetwork (that we shall call pathway) defined by **v** (i.e., those through which the flux given by **v** is not null).

### 1.3 Irreversible reactions, flux cones and polyhedral cones

In addition to the homogeneous equality constraints provided by the steady-state assumption, the flux vectors have also in general to satisfy homogeneous inequality constraints corresponding to reactions known as irreversible, whose set will be noted **Irr**:

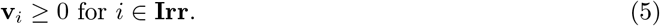

This means that fluxes in irreversible reactions are constrained to be nonnegative (in reversible reactions, fluxes may be either positive, or negative or null and the direction fixed as positive is arbitrary, the role between substrates and products being able to switch). If *r_I_* = |**Irr**|, with 0 ≤ *r_I_* ≤ *r*, is the number of irreversible reactions, the solution space *Sol*_**S,Irr**_ is the intersection of the linear subspace *FS* with *r_I_* nonnegative half-spaces, it is thus a particular case of a convex polyhedral cone, called s-cone (subspace cone or special cone) or flux cone, noted *FC*:

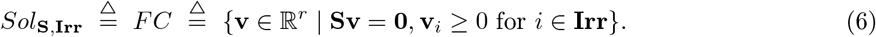

A (general) convex polyhedral cone is defined implicitly (or by intension) by finitely many homogeneous linear inequalities:

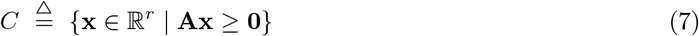

with a suitable matrix 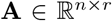 (called a representation matrix of *C*) and is thus the intersection of n half-spaces whose frontiers contain the origin. The dimension of *C*, noted *dim*(*C*), is defined as the dimension of its affine span. A flux cone corresponds thus to a particular matrix 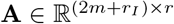 given by:

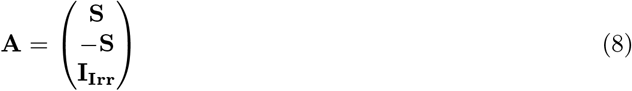

where 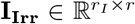 is the extension of the (*r_I_* × *r_I_*) identity matrix by columns of zeros corresponding to reversible reactions. This means that, for a flux cone, the homogeneous linear inequalities are of a special type: part of these (*m*) are actually equalities defining a lower-dimensional subspace given by the nullspace of the stoichiometric matrix **S**, the others (*r_I_*) being nonnegativity constraints regarding some single coordinate variables (given by the irreversible reactions) corresponding thus to particular half-spaces defined by such positive coordinate axes.

Conversely, to any convex polyhedral cone *C* in 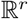 defined by a representation matrix 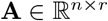, we can associate a flux cone *FC_C_* of the same dimension in 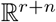 defined by:

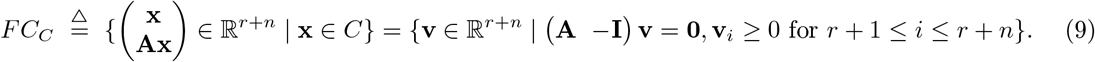

Using this correspondence (which defines a bijection of *C* onto *FC_C_*), several properties, proven for flux cones, can actually be lifted to general convex polyhedral cones.

From the definition of a cone, for every nonzero element **x** of *C*, the whole half-line {*α***x** | *α* ≥ 0} is contained in *C*. This is called a ray of *C*. Thus a flux vector is defined up to a positive scalar multiplication. The lineality space of *C* is the union of all lines of *C*, i.e. {**x** ∈ *C* | –**x** ∈ *C*}. If *C* is defined by a representation matrix **A**, its lineality space thus equals the nullspace of **A**, i.e. 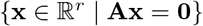. For a flux cone *FC* given by (6), its lineality space is thus constituted by the flux vectors **v** such that **v**_*i*_ = 0 for *i* ∈ **Irr**, i.e., flux vectors involving only reversible reactions (and thus the global flux can go in either one direction or the other). The cone *C* is called pointed if it does not contain a line, i.e., if its lineality space is reduced to {**0**}. For example, if *C* is contained in a closed orthant, it is pointed (where the 3^r^ closed orthants are defined by 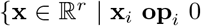 for *i* = 1,…, *r*} for an operator vector **op** ∈ {≤, =, ≥}^*r*^). In particular a flux cone *FC* with only irreversible reactions (*r_I_* = *r*) is necessarily pointed as it is included in the positive *r*-orthant (i.e., of dimension *r*). Actually, reversible flux vectors very rarely occur in metabolic networks, which therefore often give rise to pointed flux cones.

### 1.4 Extreme vectors and generating sets

We are interested in finding an explicit (by extension) representation of *C* in the form of a (minimal) set of generators. A nonzero vector **x** ∈ *C* is called extreme (or extreme pathway [37, 17] if **x** is a flux vector), if:

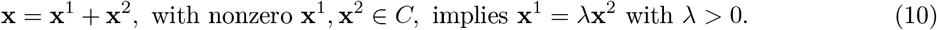

If **x** ∈ *C* is extreme, then {*α***x** | *α* ≥ 0} is called an extreme ray of *C* as all its nonzero elements are extreme (and thus for simplifying notations, we will not distinguish extreme vectors and extreme rays when it does not create confusion). In fact, *C* has an extreme ray if and only if *C* is pointed and, in this case, the extreme rays are the edges (faces of dimension one) of *C* and, according to Minkowski’s theorem, constitute the unique minimal (finite) set of generators of *C* for conical (i.e., nonnegative linear) combination:

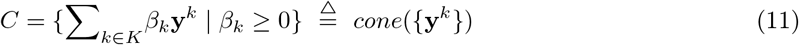

where the **y**^*k*^’s, *k* ∈ *K* (finite index set), are representatives of the extreme rays (unique up to positive scalar multiplication) and *cone* is the conical hull. More precisely, we get an upper bound for the number of extreme vectors that are sufficient to decompose any given nonzero vector 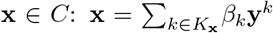 with |*K*_**x**_| ≤ *min*(*dim*(*C*), |*supp*(**x**)| + |*supp*(**Ax**)|). This result can be demonstrated first for a flux vector **v** of a pointed flux cone *FC* with an upper bound given by |*K*_**v**_| ≤ *min*(*dim*(*C*), |*supp*(**v**)|) and then for a vector **x** of a general pointed polyhedral cone *C* by using the correspondence (9) between *C* and *FC_C_* which maps the extreme vectors of *C* onto the extreme vectors of *FC_C_*. Extreme vectors of a pointed cone *C* will be noted ExVs.

The Double Description (DD) method [29, 8], known as Fourier-Motzkin, is an incremental algorithm (which processes one by one each linear inequality (**Ax**)_*j*_ ≥ 0) to build an explicit description of a pointed cone *C*, as a minimal generating matrix (whose columns are in 1-to-1 correspondence with the extreme rays), from an implicit description of *C* by a representation matrix, i.e., to enumerate its extreme rays.

If *C* is not pointed, it is still finitely generated:

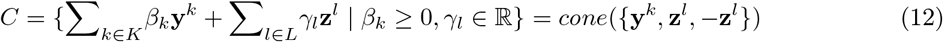

with (not unique this time) minimal set of generators consisting of basis vectors **z**^*l*^ of the lineality space and suitable vectors **y**^*k*^ not in the lineality space (e.g., the extreme vectors of the pointed cone obtained by intersecting *C* with the orthogonal complement of its lineality space). Actually, Minkowski-Weyl’s theorem for cones states that it is equivalent for a set *C* to being a polyhedral cone (7) or to being a finitely generated cone, i.e., the conical hull of a finite set of vectors (as the **y**^*k*^’s, **z**^*l*^’s and –**z**^*l*^’s).

### 1.5 Elementary vectors and conformal generating sets

Now, if the existence and uniqueness of a minimal set of generators for conical decomposition in a pointed polyhedral cone *C* is satisfactory for an explicit geometric description of *C*, it is not in general meaningful for a flux cone *FC* representing the steady-state flux vectors of a metabolic network. In fact, for a metabolic pathway, only a conical decomposition without cancelations is biochemically meaningful, since a reversible reaction cannot have a net rate in opposite directions in the contributing pathways. Indeed, the second law of thermodynamics states that a reaction can only carry flux in the direction of negative Gibbs free energy of the reaction, which is imposed by the values of the concentrations of the metabolites. This means that, when decomposing a flux vector, only so-called conformal sums, i.e., sums without cancelations, are biochemically admissible. A sum **v** = **v**^1^ + **v**^2^ of vectors is called conformal if, for all *i* ∈ {1,…, *r*}:

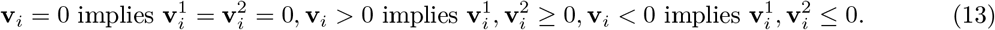

An equivalent definition is to define a sum **v** = **v**^1^ + **v**^2^ as conformal if:

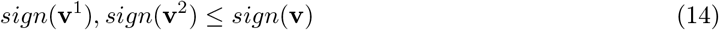

where the sign vector *sign*(**v**) ∈ {–, 0, +}^*r*^ is defined by applying the sign function component-wise, i.e., *sign*(**v**)_*i*_ = sign(**v**_*i*_), and the partial order ≤ on {–, 0, +}^*r*^ is defined by applying component-wise the partial order on {–, 0, +} induced by 0 < – and 0 < +. For example, there is a one-to-one mapping between closed orthants *O* and sign vectors **η**, defined by: 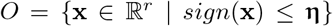 (*O* will be called defined by **η** and noted *O*_**η**_), which induces a one-to-one mapping between closed *r*-orthants and full support (i.e., with only nonzero components) sign vectors. For **ξ, η** sign vectors and **v** a vector, we say that **ξ** conforms to **η** if **ξ** ≤ **η** and that **v** conforms to **η** if *sign*(**v**) ≤ **η**. We call two vectors **v**^1^, **v**^2^ conformal if *sign*(**v**^1^), *sign*(**v**^2^) ≤ **η** for a certain sign vector **η** or, equivalently, if 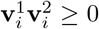 for all *i* (so, **v**^1^ + **v**^2^ is a conformal sum if and only if **v**^1^ and **v**^2^ are conformal). A conformal sum **v** = **v**^1^ + **v**^2^, i.e., verifying (14), will be noted **v** = **v**^1^ ⊕ **v**^2^. It is therefore natural to look for generators as conformally non-decomposable vectors, where a nonzero vector **x** of a convex polyhedral cone *C* is called conformally non-decomposable, if:

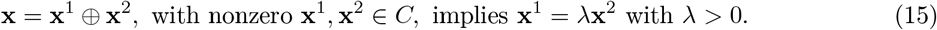

A vector **x** (resp., flux vector **v**) of a convex polyhedral cone *C* (resp., a flux cone *FC*) is called elementary [22] if it is conformally non-decomposable. All nonzero elements of the ray defined by an elementary vector are elementary, i.e., elementary vectors are unique up to positive scalar multiplication (and thus we will often not distinguish elementary vectors and elementary rays). Elementary vectors (resp., flux vectors) will be noted EVs (resp., EFVs).

The elementary rays constitute the unique minimal (finite) set of conformal generators of *C*, i.e., generators for conformal conical sum:

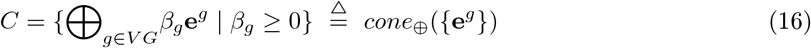

where the **e**^*g*^’s, *g* ∈ *VG* (finite index set), are representatives of the elementary rays (unique up to positive scalar multiplication) and *cone*_⊕_ is the conical conformal hull. More precisely, we get an upper bound for the number of elementary vectors that are sufficient to decompose any given nonzero vector 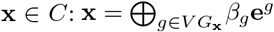 with |*VG*_**x**_| ≤ *min*(*dim*(*C*), |*supp*(**x**)| + |*supp*(**Ax**)|). This result can be demonstrated first for a flux vector **v** of a flux cone *FC* with an upper bound given by |*VG*_**v**_| ≤ *min*(*dim*(*C*), |*supp*(**v**)|) and then for a vector **x** of a general polyhedral cone *C* by using the correspondence (9) between *C* and *FC_C_* which maps the elementary vectors of *C* onto the elementary flux vectors of *FC_C_*.

It follows from (10) and (15) that an extreme vector of *C* is elementary, i.e., ExVs ⊆ EVs, but the converse is generally false since a conformally non-decomposable vector may be conically decomposable. Nevertheless, if *C* is contained in a closed orthant, there is identity between extreme vectors and elementary vectors, i.e., ExVs = EVs. More precisely, for nonzero **x** ∈ *C* and *O* a closed orthant with **x** ∈ *O*, then **x** is elementary in *C* if and only if it is extreme in *C* ∩ *O*. It results that the elementary vectors of *C* are the extreme vectors of intersections of *C* with any closed orthant:

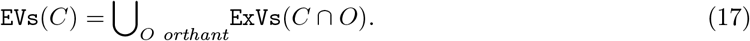

Elementary vectors of *C* can thus be obtained by using algorithms, such as DD, to compute extreme vectors ExVs of the pointed polyhedral cones *C* ∩ *O*. By doing this, it is convenient to select in (17) only a minimal subset of closed orthants *O*, in order to avoid equality or inclusion between the *C*∩*O*’s (nonempty intersection can obviously not be avoided as orthants are closed). It is clearly enough to consider only the 2^r^ closed r-orthants of maximal dimension *r*, but this does not avoid equality or inclusion. Let {**η**^*i*^} be the maximal (for the partial order defined above on {–, 0, +}^*r*^) sign vectors of *sign*(*C*) = {*sign*(**x**) | **x** ∈ *C*}. It is then enough in (17) to consider the closed orthants 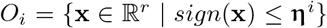 and there is no equality or inclusion between the *C* ∩ *O_i_*’s, where *C* ∩ *O_i_* = *C*_≤**η**^*i*^_ = {**x** ∈ *C* | *sign*(**x**) ≤ **η**^*i*^}. The *C*_≤**η**^*i*^_’s are called topes, noted Ts (flux topes [12] noted FTs for a flux cone *FC*). *C* is thus decomposed into topes and (17) can be rewritten as:

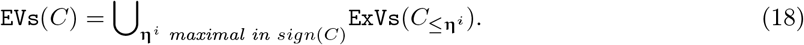

Note that, for a flux cone *FC* (6), a FT is defined by specifying a maximal subset of reactions with fixed directions (thus fixing the directions of reversible reactions), the others having necessarily a zero flux. This simplifies if *FC* is consistent, i.e., without unused reaction, which means that every reaction, in every possible direction for reversible reactions, is supported by a flux vector: ∀*i* ∈ {1,…, *r*} ∃**v** ∈ *FC* **v**_*i*_ > 0 and ∀*i* ∈ {1,…, *r*}\**Irr** ∃**v** ∈ *FC* **v**_*i*_ < 0. We can always assume *FC* consistent after a preprocessing step (practically, this can be achieved by using flux variability analysis [14]) that removes all reactions that cannot carry nonzero steady-state flux and changes all reversible reactions that cannot carry flux in both directions into irreversible ones. In this case, every remaining reaction in every possible direction is supported by a flux vector with full support (i.e., with nonzero flux in any reaction) and all FTs *FC*_≤**η**^i^_ have full support, i.e., the **η**^*i*^’s have full support or, equivalently, the *O_i_*’s are *r*-orthants. An obvious upper bound for the number of FTs is thus 2^*r−r*_*I*_^.

Now, for a flux cone *FC*, another commonly used method is, at the extreme opposite, to have it included into a single (positive) orthant in a higher dimension by splitting each reversible reaction i into a forward *i*^+^ and a backward *i*^−^ irreversible reaction. This means decomposing a flux in *i* as 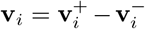 with 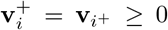 and 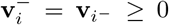. Columns of the stoichiometric matrix **S** corresponding to reversible reactions *i* are negated (which means exchanging the roles of substrates and products in *i*) and appended to **S** as new columns to form the new stoichiometric matrix 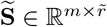, where 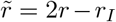 is the new number of reactions and all 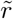 reactions are now irreversible, 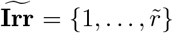. *FC* is in one-to-one correspondence with vectors **v** of 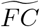 such that **v**_*i*_+.**v**_*i*^−^_ = 0 for *i* reversible. In particular, the fluxes of the form **v**_*i*^+^_ = **v**_*i*^−^_ > 0 with all other components being null are obtained as extreme vectors of 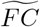 but represent futile cycles (involving reactions *i*^+^ and *i*^−^) without biological reality and must be eliminated. Finally, EFVs(*FC*) are in one-to-one correspondence with 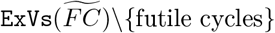 (called at the origin extreme currents in stoichiometric network analysis [6]). 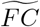 is included in the positive 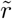-orthant and has thus only one FT.

We therefore have two opposite ways of dealing with reversible reactions for computing EFVs of a flux cone *FC*: either splitting each reversible reaction into two irreversible ones, such that *FC* is reduced to a single FT at the price of an increase in the space dimension by *r* – *r_I_* (which can cause serious efficiency problems to algorithms such as DD) or keeping the reversible reactions unchanged and decomposing *FC* into FTs, in each of which the directions of reversible reactions are fixed, at the price of a potentially exponential (in terms of *r* – *r_I_*) number of FTs to consider. All intermediate cases, where only a subset of reversible reactions are split into irreversible ones and the others are processed by decomposition into FTs, are obviously possible. Independently of the solution adopted, we will work most of the time in a given FT for *FC*, defined by a (full support if *FC* is consistent) sign vector **η**, and the EFVs of *FC* which conform to **η** are thus given by the ExVs of this FT *FC*_≤**η**_.

### 1.6 Elementary modes

The null value 0 plays a component-wise crucial role in definitions of the support of a vector (4), of a flux cone (6) and of a conformal sum (13),(14). A close relationship results between support-minimal vectors and elementary vectors in a flux cone. A nonzero vector **x** ∈ *C* is called support-minimal, if:

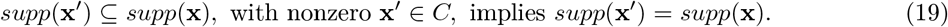

A nonzero vector **x** (resp., flux vector **v**) of a convex polyhedral cone *C* (resp., a flux cone *FC*) is called elementary mode (resp., elementary flux mode) if it is support-minimal (the concept of elementary flux mode was first introduced, under the name of elementary vector [35], for a subspace of 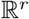, i.e., for a flux linear subspace *FS* (3) and then [39] for a flux cone *FC* with a definition actually closer to that of a support-wise non-decomposable vector). All nonzero elements of the ray defined by an elementary mode are elementary modes having the same support, i.e., elementary modes are unique up to positive scalar multiplication (and we will therefore in general identify two positively proportional elementary modes). Elementary modes (resp., flux modes) will be noted EMs (resp., EFMs). The concept of EFM in a flux cone FC is biologically significant as it represents a minimal pathway operating in a steady state, i.e., with all reactions involved necessarily active (with a nonzero net rate), which means that no proper sub-pathway can operate in a steady state.

Note that, if **x** is an EFM, then *sign*(**x**) is a minimal (for the partial order defined above on {–, 0, +}^*r*^) nonzero element of *sign*(*FC*) and, conversely, it is shown that a minimal nonzero sign vector **σ** ∈ *sign*(*FC*) determines an EFM **x** with *sign*(**x**) = **σ** by: *FC*_≤σ_ = {**v** ∈ *FC* | *sign*(**v**) ≤ **σ**} = {**v** ∈ *FC* | *sign*(**v**) = **σ**} ∪ {**0**} = {*λ***x** | λ ≥ 0}. There is thus a one-to-one mapping between EFMs and minimal nonzero sign vectors of *sign*(*FC*). Comparing with the one-to-one mapping between FTs and maximal sign vectors of *sign*(*FC*), we see that EFMs and FTs are dual concepts.

Now, for a flux cone *FC*, support-minimality and conformal non-decomposability are equivalent properties, i.e., there is identity between elementary flux modes and elementary flux vectors: EFMs = EFVs. For metabolic pathways in a flux cone *FC*, there is therefore identity between minimal (for support inclusion) pathways and non-decomposable (for conformal sum) pathways. From (16), the EFMs constitute a conformal generating set (i.e., generating set for conformal sum) for *FC*, and in fact the unique minimal such set (for that matter one way of proving (16) for flux cones *FC* is to prove it with EFMs as a conformal generating set and to prove that EFMs = EFVs). From (18), for any maximal sign vector **η** of *sign*(*FC*), the EFMs of *FC* which conform to **η** are the EFMs of the FT *FC*_≤**η**_ and coincide with the ExVs of the said FT, this result being the basis of methods for computing EFMs [9, 7]. EFMs are thus decomposed into subsets according to the decomposition of *FC* into flux topes [12]: the EFMs of the FT *FC*_≤**η**_ correspond to the *FC*_≤**σ**_’s, where the **σ**’s are the minimal nonzero sign vectors of *sign*(*FC*) such that **σ** ≤ **η**.

For a general polyhedral cone *C*, there is no direct relationship between elementary modes and elementary vectors but the correspondence (9) between *C* and the higher dimensional flux cone *FC_C_* maps the elementary vectors of *C* onto the elementary flux vectors of *FC_C_*, i.e., the elementary flux modes of *FC_C_*:

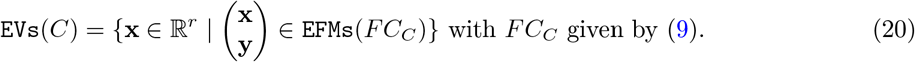

And it remains true that any vector which is support-minimal in a given *C*_≤**η**_ is actually support minimal in *C*, as it depends only on the convexity of *C*: if **x** and **x′** are vectors in a convex set with *supp*(**x′**) ⊂ *supp*(**x**), then a vector **x″** exists in this convex set with *sign*(**x″**) ≤ *sign*(**x**) and *supp*(**x″**) ⊂ *supp*(**x**) (we take **x″** = *λ***x′** + (1 – *λ*)**x** with *λ* minimal in (0, 1] such that 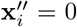 for a certain i with **x**_*i*_ = 0). Thus EMs of *C* can be computed tope by tope:

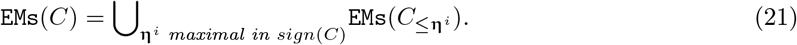

### 1.7 Inhomogeneous linear constraints and polyhedra

Additionally, in this standard setting, fluxes may be constrained by other constraints, typically lower and upper bounds regarding reaction rates:

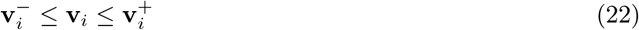

or, more generally, any set of inhomogeneous linear constraints, noted **ILC**, that can be written in the general form:

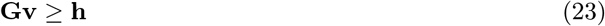

where 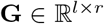 is a matrix and 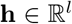 a vector with nonzero components, defining a general inhomogeneous convex polyhedron 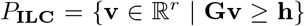. The solution space *Sol*_**s,irr,ILC**_ is thus now a s-polyhedron or flux polyhedron noted *FP* and defined by:

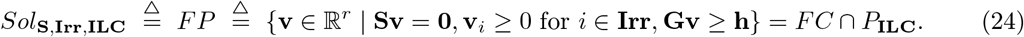

This is a particular case of (general) convex polyhedron that is defined implicitly by finitely many linear inequalities:

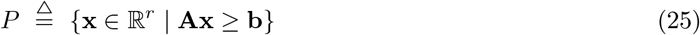

with a suitable matrix 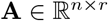 and vector 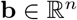 and is thus the intersection of *n* (affine) half-spaces. Its dimension is defined as the dimension of its affine span. In this way, a flux polyhedron *FP* corresponds to a particular matrix 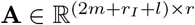 and vector 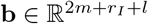 given by:

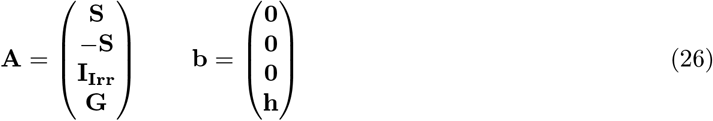

meaning that the inequalities that are homogeneous actually divide into *m* equalities defining a lower-dimensional subspace given by the nullspace of the stoichiometric matrix **S** and into *r_I_* nonnegativity constraints regarding certain single coordinate variables (given by the irreversible reactions). A polyhedral cone (resp., flux cone) is a special case of polyhedron (resp., flux polyhedron) where **b** = **0** (resp., 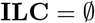).

To any nonempty polyhedron *P* given by (25) is associated its so-called recession cone 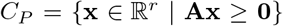, which is the polyhedral cone containing all unbounded directions (rays) of *P* (if *P* is a polyhedral cone, then *P* = *C_P_*). A bounded polyhedron is called a polytope and thus *P* is a polytope if and only if its recession cone is trivial: *C_P_* = {**0**}. *P* is called pointed if its recession cone *C_P_* is pointed, i.e., if its lineality space 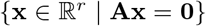, also called the lineality space of *P* (as it contains all unbounded lines of *P*), is trivial. For a flux polyhedron *FP*, we have: *C_FP_* = *FC* ∩ *C_P_**ILC**__* (thus *FP* can be a polytope without *P*_**ILC**_ being so and be pointed without either *FC* or *P*_**ILC**_ being so). Note that *C_FP_* is not in general a flux cone.

#### 1.7.1 Extreme points and vectors and generating sets

A vector **x** ∈ *P* is called an extreme point, if it cannot be written as a convex combination of distinct vectors of *P*:

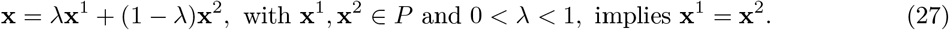

Extreme points coincide with vertices of *P*, where a vertex of *P* is defined as a face of dimension 0. *P* is pointed if and only if it has a vertex and in this case, according to Minkowski’s theorem, the vertices of *P* and the extreme rays of *C_P_* constitute the unique minimal (finite) set of (“bounded” and “unbounded”, resp.) generators of *P* for (convex and conical, resp.) combination:

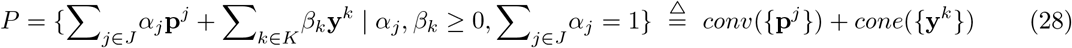

where the **p**^*j*^’s, *j* ∈ *J* (finite index set), are the extreme points (vertices) of *P*, noted ExPs, and the **y**^*k*^’s, *k* ∈ *K* (finite index set), are the extreme vectors of *C_P_* (unique up to positive scalar multiplication), noted ExVs, and *conv* is the convex hull (if *P* is a pointed polyhedral cone, then it has only one vertex, which is the zero vector, and the formula (28) reduces to (11)). More precisely, we get an upper bound for the number of extreme points and vectors that are sufficient to decompose any given vector 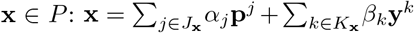 with |*J*_**x**_| + |*K*_**x**_| ≤ *min*(*dim*(*P*) + 1, |*supp*(**x**)| + |*supp*(**Ax – b**)| + 1). This result can be deduced from result (11) for a pointed flux cone *FC* by using the following correspondence between *P* and such a flux cone.

In fact, to any convex polyhedron *P* in 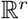 defined by a matrix 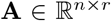 and vector 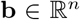 (25), we can associate a flux cone *FC_P_* in a higher dimension 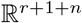 defined by:

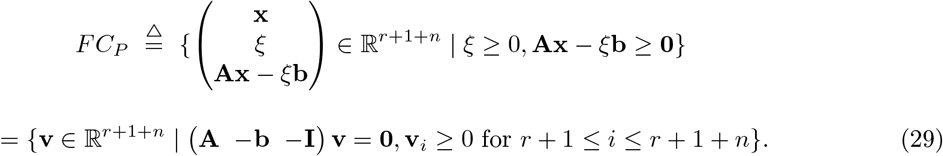

This introduces a correspondence between vectors **x** of *P* and vectors 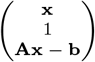 of *FC_P_*, which maps vertices of *P* onto extreme vectors of *FC_P_* with component *ξ* = 1, and between vectors **x** of *C_P_* and vectors 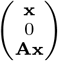 of *FC_P_*, which maps extreme vectors of *C_P_* onto extreme vectors of *FC_P_* with component *ξ* = 0. Thanks to this correspondence, several properties, proven for flux cones, can be lifted to general convex polyhedra.

For the particular case where *P* is a flux polyhedron *FP* given by (24), the correspondence (29) simplifies by associating to *FP* the flux cone *FC_FP_* in dimension 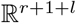 defined by:

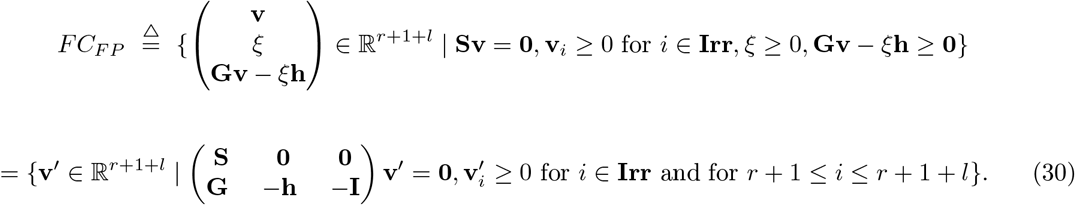

The DD method builds an explicit description of a pointed polyhedron *P*, in the form of two generating matrices whose columns are respectively the **p**^*j*^’s and the **y**^*k*^’s, from an implicit description of *P* as in (25), i.e., enumerates its vertices ExPs and extreme vectors ExVs.

If *P* is not pointed, it is still finitely generated:

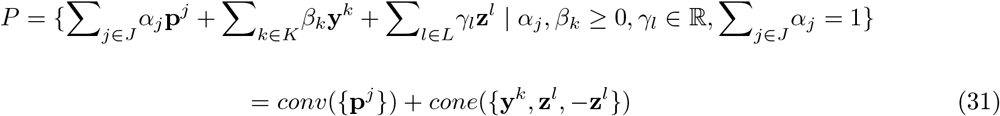

with a (not unique this time) minimal set of generators consisting of basis vectors **z**^*l*^ of the lineality space and suitable vectors **p**^*j*^ and **y**^*k*^ (e.g., the vertices and extreme vectors of the pointed polyhedron obtained by intersecting *P* with the orthogonal complement of its lineality space; if *P* is a non-pointed polyhedral cone, there is no nonzero **p**^*j*^ and formula (31) reduces to (12)). In fact, Minkowski-Weyl’s theorem for polyhedra states that it is equivalent for a set *P* to be a polyhedron (25) or to be finitely generated, i.e., to be the Minkowski sum of the convex hull of a finite set of vectors (as the **p**^*j*^’s) and of the conical hull of a finite set of vectors (as the **y**^*k*^’s, **z**^*l*^’s and –**z**^*l*^’s).

#### 1.7.2 Elementary points and vectors and conformal generating sets

As was the case for a flux cone, however, such decomposition into a finite set of generators is not in general satisfactory for a flux polyhedron as only a decomposition without cancelations is biochemically meaningful. In the same way we replaced as generators for a polyhedral cone extreme vectors by conformally non-decomposable vectors, we will replace as generators for a polyhedron *P* extreme points (vertices) by convex-conformally non-decomposable vectors (and for its recession cone *C_P_* extreme vectors by conformally non-decomposable vectors). A vector **x** of a polyhedron *P* is called convex-conformally non-decomposable, if:

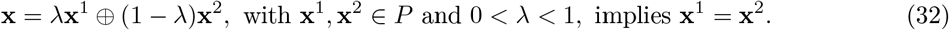

Given a polyhedron *P* (resp., flux polyhedron *FP*), a vector (resp., flux vector) **x** is called an elementary point (resp., elementary flux point) - also called “bounded” elementary vector - of *P* if **x** ∈ *P* is convex-conformally non-decomposable and is called an elementary vector (resp., elementary flux vector) - also called “unbounded” elementary vector - of *P* if **x** ∈ *C_P_* is conformally non-decomposable (it is unique only up to positive scalar multiplication) [22]. Elementary points (resp., flux points) will be noted EPs (resp., EFPs) and elementary vectors (resp., flux vectors) will be noted EVs (resp., EFVs), which is consistent with the same notation for polyhedral cones and flux cones. We will note Es = EPs ∪ EVs (resp., EFs = EFPs ∪ EFVs) the elementary elements (resp., elementary fluxes) of *P* (resp., *FP*).

The elementary points and the elementary rays constitute the unique minimal (finite) set of conformal generators of *P*, i.e., generators for convex-conformal (for elementary points) and conformal (for elementary vectors) sum:

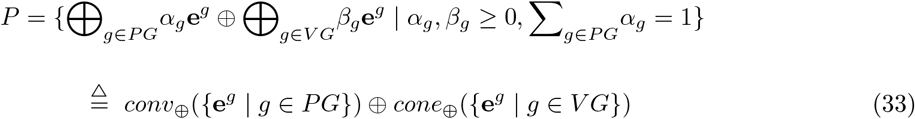

where the **e**^*g*^’s, *g* ∈ *PG* (finite index set), are the elementary points and the **e**^*g*^’s, *g* ∈ *VG* (finite index set), are the elementary vectors (unique up to positive scalar multiplication), and *conv*_⊕_ is the convex conformal hull. More precisely, we get an upper bound for the number of elementary points and vectors that are sufficient to decompose any given vector 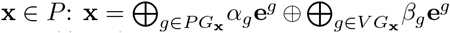 with |*PG*_**x**_| + |*VG*_**x**_| ≤ *min*(*dim*(*P*) + 1, |*supp*(**x**)| + |*supp*(**Ax** – **b**)| + 1). This result can be demonstrated from result (16) for a flux cone by using the correspondence (29) between *P* and *FC_P_* which maps the elementary points of *P* onto the elementary flux vectors of *FC_P_* with component *ξ* = 1 and the elementary vectors of *P* onto the elementary flux vectors of *FC_P_* with component *ξ* = 0.

We already know that an extreme vector of *C_P_* is elementary, and is therefore an elementary vector of *P*, i.e., ExVs ⊆ EVs. It follows from (27) and (32) that an extreme point (vertex) of *P* is an elementary point of *P*, i.e., ExPs ⊆ EPs, but the converse is generally false since a convex-conformally non-decomposable vector may be convex decomposable. Nevertheless, if *P* is contained in a closed orthant (and thus *C_P_* too), any sum of vectors of *P* (resp., *C_P_*) is conformal and thus there is identity between extreme points (vertices) and elementary points (resp., between extreme vectors and elementary vectors), i.e., ExPs = EPs and ExVs = EVs. More precisely, for **x** ∈ *P* (resp., **x** ∈ *C_P_* and nonzero) and *O* a closed orthant with **x** ∈ *O*, then **x** is an elementary point (resp., elementary vector) of *P* if and only if it is a vertex in *P*∩*O* (resp., an extreme vector in *C_P_*∩*O* = *C*_*P*∩*O*_). It follows that the elementary points (resp., elementary vectors) of *P* are the vertices (resp., extreme vectors) of intersections of *P* (resp., *C_P_*) with any closed orthant, which are pointed subpolyhedra (resp., pointed subcones):

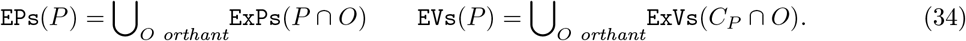

Note in particular that, if **0** ∈ *P*, then **0** ∈ EPs. Elementary points and vectors can therefore be obtained by using algorithms, such as DD, to compute vertices ExPs and extreme vectors ExVs of the pointed polyhedra *P* ∩ *O*. It is obvious that considering only the 2^*r*^ closed *r*-orthants is enough. Now, as for polyhedral cones, decomposing the polyhedron into topes is better:

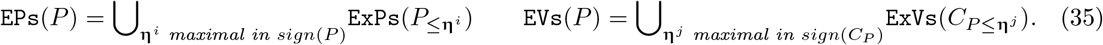

If *O_i_* is the closed orthant defined by **η**^*i*^, then the corresponding tope for *P* is *P* ∩ *O_i_* = *P*_≤**η**^*i*^_ = {**x** ∈ *P* | *sign*(**x**) ≤ **η**^*i*^} and, as seen for polyhedral cones, *C_P_* ∩ *O_j_* = *C*_*P*_≤**η**^*j*^__ is a tope for the recession cone *C_P_*. Examine how the equality *C*_*P*_≤**η**__ = *C*_*P*_≤**η**__, for **η** an arbitrary sign vector, can be expressed in terms of topes. Note first that any **η**^*j*^ is dominated by at least one **η**_*i*_ for the partial order ≤ on { –, 0, +}^*r*^: ∀**η**^*j*^∃**η**^*i*^**η**^*j*^ ≤ **η**_*i*_, which means that *O_j_* is a sub-orthant of *O_i_*, and we have *C*_*P*_≤**η**^*j*^__ = *C*_*P*_≤**η**^*i*^__, expressing the relation between the topes for the recession cone of the polyhedron and the recession cones of certain of the polyhedron topes (precisely those topes *P*_≤**η**^*i*^_ for which **η**^*i*^ dominates an **η**^*j*^, necessarily unique). More generally, the recession cone of any tope *P*_≤**η**^*i*^_ for *P* can be expressed as a subcone of a tope for the recession cone of *P* by: *C*_*P*_≤**η**^*i*^__ = *C*_*P*_≤**c**(**η**^*i*)^__, where **c**(**η**^*i*^) = *max*{**η** ∈ *sign*(*C_P_*) | **η** ≤ **η**^*i*^} is the greatest (it is necessarily unique) sign vector in *sign*(*C_P_*) dominated by **η**^*i*^ (thus, if **η**^*i*^ does not dominate any **η**^*j*^, *C*_*P*≤_**c**(**η**^*i*^)__ is not a tope for *C_P_*; this is the case for example if *P* is not a polytope but *P*_≤**η**^*i*^_ is, implying that **c**(**η**^*i*^) = **0** and that *C*_*P*≤_**c**(**η**^*i*^)__ = {**0**} is not a tope for *C_P_* ≠ {**0**}).

For the particular case of a flux polyhedron *FP* (24), we have *FP*_≤**η**_ = *FC*_≤**η**_ ∩ *P*_**ILC**_ and *C*_*FP*_≤**η**__ = *C*_*FP*≤_**η**__ = *FC*_≤**η**_ ∩ *C*_*P*_**ILC**__, for any sign vector **η**. Any flux tope *FP*_≤**η**^*i*^_ for *FP* can thus be expressed as: *FP*_≤**η**^*i*^_ = *FC*_≤**η**^*k*^_ ∩ *P*_**ILC**_, for a certain flux tope *FC*_≤**η**^*k*^_ for *FC*, i.e., by applying the constraints **ILC** to a flux tope for *FC*. The same holds for the recession cone: 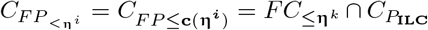, i.e., by applying the homogeneous counterparts of constraints **ILC** to a flux tope for *FC*. If *FP* is assumed to be consistent (same definition as for a flux cone, i.e., without unused reaction, always with a zero flux), all FTs *FP*_≤**η**^*i*^_ then have full support, i.e., the **η**^*i*^’s have full support or, equivalently, the *O_i_*’s are *r*-orthants. An obvious upper bound for the number of FTs is thus 2^*r–r_I_*^. Note that *C_FP_* to be consistent is a sufficient (but not necessary) condition for *FP* to be consistent. And that, if *FP* is consistent, so is *FC* (but *FP* can be inconsistent even if both *FC* and *P*_**ILC**_ are consistent).

The method used to include a flux cone into a single (positive) orthant in higher dimension, by splitting each reversible reaction *i* into a forward *i*^+^ and a backward *i*^−^ irreversible reaction, applies as well to a flux polyhedron *FP*. Matrix **G** is extended into matrix 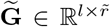 in the same way that **S** is extended into 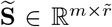, where 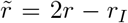 is the new number of reactions and all 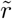 reactions are now irreversible, 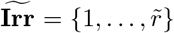. *FP* is in one-to-one correspondence with vectors **v** of 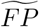 such that **v**_*i*+_.**v**_*i*^−^_ = 0 for *i* reversible. In particular, the fluxes with **v**_*i*^+^_, **v**_*i*^−^_ > 0 for a certain *i*, which are obtained as vertices or extreme vectors of 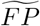, are futile (involving a net rate in opposite directions in reactions *i*^+^ and *i*^−^) without biological reality and must be eliminated (they are not generally limited to futile cycles as in flux cones). Finally, the set of elementary fluxes EFs(*FP*) is in one-to-one correspondence with 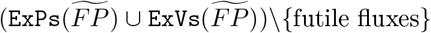. As 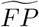 is included in the positive 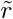-orthant, it has only one FT.

As for flux cones, there are two opposite ways of dealing with reversible reactions for computing EFs = EFPs ∪ EFVs of a flux polyhedron *FP*: either splitting each reversible reaction into two irreversible ones, reducing *FP* to a single FT in higher dimension, or keeping the reversible reactions unchanged and decomposing *FP* into FTs without increasing the dimension. And all intermediate cases are possible. Whatever the solution adopted, EFs are obtained as the union of ExPs and ExVs for each flux tope for *FP* and thus we will generally work in a given FT for *FP*, defined by a (full support if *FP* is consistent) sign vector **η**, and the EFs of *FP* which conform to **η** are thus given by the ExPs and ExVs of this FT *FP*_≤**η**_.

#### 1.7.3 Elementary modes

For a general polyhedron *P* (and even for a flux polyhedron *FP*), there is no direct relationship between support-minimal (19) vectors and elementary elements as it was the case for flux cones *FC*. Nevertheless, from (33), it follows that, for any support-minimal nonzero vector in *P* (still called elementary mode EM), there is an elementary element with the same support. Thus, all minimal support patterns of vectors appear in the set of supports of elementary elements and are actually the minimal elements in this set for subset inclusion:

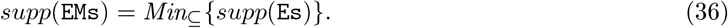

In particular, for a metabolic network, this means that any set of reactions involved in a minimal pathway (EFM) appears as the set of reactions involved in a certain elementary flux (EF).

In addition, for a general polyhedron *P*, the correspondence (29) between *P* and the higher dimensional flux cone *FC_P_* maps the elementary points (resp., elementary vectors) of *P* onto the elementary flux vectors, i.e., the elementary flux modes, of *FC_P_* with component *ξ* = 1 (resp., with component *ξ* = 0):

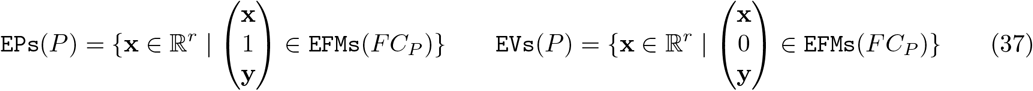

with *FC_P_* given by (29). The same formula holds for the particular case of a flux polyhedron *FP* (24) with *FC_FP_* given by (30).

And, as seen in the proof of (21), any vector which is support-minimal in a certain *P*_≤**η**_ is actually support minimal in *P*, thus EMs of *P* can be computed tope by tope:

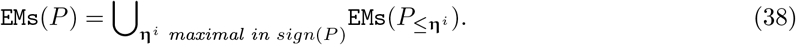

### 1.8 Complexity results

Enumerating the extreme vectors of a polyhedral cone or the vertices of a polytope has been proven to be a #P-complete problem. A consequence is that enumerating EFs or EFMs of a metabolic network is #P-complete too [1, 2]. The problem of enumerating the extreme vectors of a polyhedral cone or the vertices of a polytope with a polynomial delay (i.e., the time between one output item and the next is bounded by a polynomial function in terms of the input size) is an open problem. So the possibility of enumerating EFs or EFMs of a metabolic network with a polynomial delay is an open problem (except for a flux cone with all reactions being reversible, i.e., a flux linear subspace (3), as the EFMs are then the circuits of a linear matroid [35]). Given a metabolic network, deciding if an EF or EFM exists with a given support of size at least two is an NP-complete problem. The same result holds for deciding if an EF or EFM exists whose support size is bounded above by a given positive integer [1, 2].

The McMullen’s upper bound theorem [27] states that, for any fixed positive integers *d* and *n*, the maximum number of *j*-faces of a *d*-polytope with *n* facets (i.e., faces of dimension *d* – 1) is attained by the dual cyclic polytope *c*\*(*d, n*) for all *j* = 0, 1,…, *d*–2. A consequence is that the maximum number of vertices of a *d*-polytope with *n* facets is given by 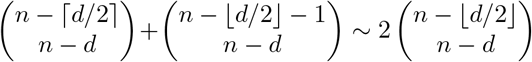. We thus obtain that the number of EFs or EFMs in a metabolic network (after having split each reversible reaction into two irreversible ones) is bounded above by a quantity approximately equal to 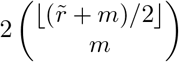 with 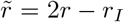 (so 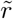 varies between *r* and 2*r*). If the number *m* of internal metabolites is small compared to the total number 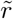 of reactions (after having split reversible reactions), the number of EFs or EFMs is then bounded above by a quantity of order 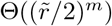. If *m* is close to 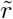, this number is bounded above by a quantity of order 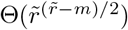. The worst case occurs when *m* is close to 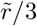 with an upper bound approximately equal to 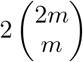. We obtain the same results with *r* instead of 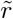 if we do not split reversible reactions but fix their signs arbitrarily, i.e., if we consider EFs or EFMs in an arbitrary closed *r*-orthant *O*, i.e., in an arbitrary flux tope for *FC* (or *FP*).

## 2 Metabolic pathways in the presence of biological constraints

Although the current improved implementations of the DD method [40, 43] allow the computation of millions, even billions, of EFMs or EFs, tackling genome-scale metabolic models (GSMMs) is still beyond our reach. Moreover, most of the computed EFMs or EFs are not biologically valid, because only stoichiometry and certain flux constraints, such as irreversibility of reactions or bounds on reaction rates, are taken into account. For both scaling up to large systems and limiting the number of biologically invalid solutions, it is necessary to consider additional biological constraints, such as thermodynamic, kinetic or regulatory constraints.

### 2.1 Biological constraints

#### 2.1.1 Thermodynamic constraints

Assuming constant pressure and a closed system, according to the second law of thermodynamics, a reaction *i* proceeds spontaneously only in the direction of its negative Gibbs free energy Δ_*r*_*G_i_* [3], given by:

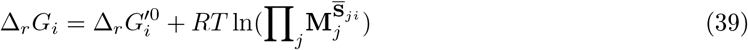

where: 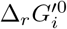 is the standard Gibbs free energy of reaction *i, R* the molar gas constant, *T* the absolute temperature, **M**_*j*_ the (positive) concentration of metabolite *j* and 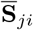 is the stoichiometric coefficient of metabolite *j* in reaction *i* (i.e., 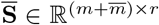 is the extension of the stoichiometric matrix **S** to the 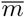 external metabolites). This means that Δ*_r_G_i_* < 0, (resp., Δ*_r_G_i_* > 0) is a necessary condition to get **v**_*i*_ > 0 (resp., **v**_*i*_ < 0), which can be expressed by the constraint: *sign*(**v**_*i*_) ≤ –*sign*(Δ*_r_G_i_*). And that a flux vector **v** is thermodynamically feasible [16] if and only if all its components **v**_*i*_ satisfy such a constraint (it is enough to consider those *i* ∈ *supp*(**v**) as the constraint is trivially satisfied when **v**_*i*_ = 0). The thermodynamic constraint for **v**, that depends on the vector **M** of metabolite concentrations, can thus be defined as:

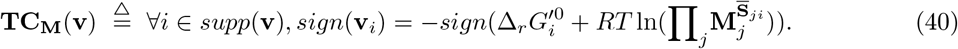

As at equilibrium Δ*_r_G_i_* is null, we obtain: 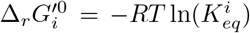, where 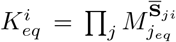 is the equilibrium constant of reaction *i*. Thus, Δ*_r_G_i_* can be rewritten as 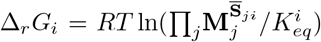 and the thermodynamic constraint **TC_M_**(**v**) as:

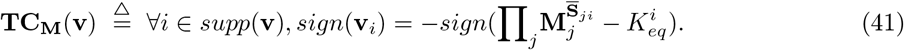

Often, the concentrations of external metabolites can be measured and included in the constraint as known parameters, keeping only a dependency of the constraint on the concentrations of internal metabolites. The formula 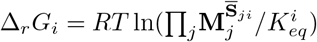 can be rewritten, by dividing the numerator and denominator of the fraction by the terms dealing with external metabolites, as 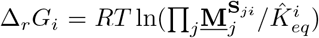, where 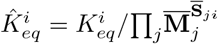 is the apparent equilibrium constant of the reaction *i* and <u>**M**</u> (resp., 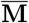) the vector of internal (resp., external) metabolite concentrations. The thermodynamic constraint (41) can thus be rewritten as:

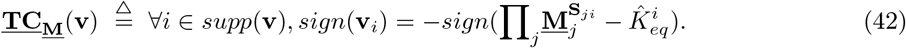

For given metabolite concentrations vector **M** (resp., internal metabolite concentrations vector <u>**M**</u>), let **ts_M_** ∈ {–, 0, +}^*r*^ (resp., **ts_<u>M</u>_**) be the fixed thermodynamic sign vector defined by:

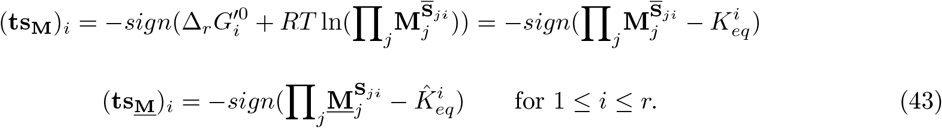

Then the thermodynamic constraint **TC_M_** (resp., <u>**TC_M_**</u>) can be rewritten as:

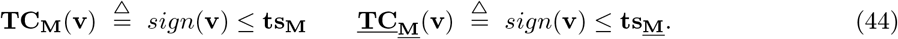

Thus the set *Sol*_**TC_M_**_ (resp., *Sol*_<u>**TC_M_**</u>_) of vectors **v** satisfying the constraint **TC_M_**(**v**) (resp., <u>**TC_M_**</u>(**v**)), given by 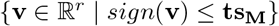 (resp., 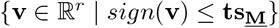), is the closed orthant *O*_**tS_M_**_ (resp., *O*_**ts_<u>M</u>_**_) defined by **ts_M_** (resp., **ts_<u>M</u>_**), of dimension *r* if **ts_<u>M</u>_** (resp., **ts_<u>M</u>_**) has full support, i.e., (**ts_M_**)_*i*_ ≠ 0 (resp., (**ts_<u>M</u>_**)_*i*_ ≠ 0) for all *i*, of lesser dimension otherwise.

##### Lemma 2.1.

*Given metabolite concentrations **M** (resp., internal metabolite concentrations **<u>M</u>**), the set Sol_**TC_M_** (resp., Sol_<u>**TC_M_**</u>__) of vectors in* 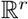 *satisfying the thermodynamic constraint **TC_M_** (resp., <u>**TC_M_**</u>) is the set of vectors that conform to the thermodynamic sign vector **ts_M_** (resp., **ts_<u>M</u>_**)* (43), *i.e., the closed orthant O_**tS_M_**_ (resp., O_ts_<u>M</u>__) defined by **ts_M_** (resp., **ts_<u>M</u>_**)*.

#### 2.1.2 Kinetic constraints

Metabolic reactions are catalyzed by enzymes. The catalytic mechanisms of key enzymes have been investigated in great detail and described by mathematical formulas. However many kinetic equations are still unknown and have to be substituted by standard rate laws such as mass-action kinetics, power laws, reversible Hill kinetics, lin-log kinetics, convenience kinetics, generic rate equations or TKM rate laws [24]. What is important for our study is that these modular rate laws share the general form:

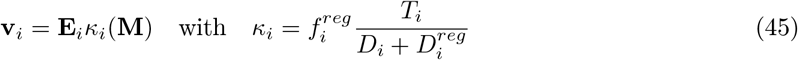

where **E**_*i*_ is the (nonnegative) level of the enzyme catalyzing the reaction *i* (given either as an amount or as a concentration, in which case the rate law is pre-multiplied by the compartment volume) and *κ_i_* depends on the concentrations of the metabolites occurring in *i* (reactants of *i*) and on reaction *i* stoichiometry, rate law considered, allosteric regulation and parameters. In the general form of *κ_i_, T_i_* is the thermodynamic numerator (which can be written in the compact form 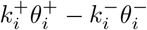 with turnover rate parameters 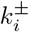) that gives its sign to *κ_i_* (and thus to **v**_*i*_) and reflects the relationship between chemical potentials and reaction directions and ensures that the rate vanishes at chemical equilibrium, 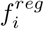 and 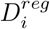, both positive, implement enzyme regulation (partial or complete for the first one, specific for the second) and *D_i_* is the (positive) kinetic denominator, a polynomial of scaled reactant concentrations whose terms correspond to different binding states of the enzyme (reducing the enzyme amount available for catalysis), which depends on the rate law considered.

The kinetic constraint for **v** depends both on the vector **E** of enzyme concentrations and on the vector **M** of metabolite concentrations and can thus be defined as:

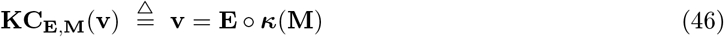

where ∘ is the component-wise product of vectors: (**E** o ***κ***)_*i*_ = **E***_i_κ_i_*. This means that the flux vector is a component-wise linear function of the vector of enzyme concentrations (and a nonlinear function of the metabolite concentrations vector). For nonzero **E**_*i*_, the sign of *T_i_* gives the direction in which reaction *i* proceeds, so in this sense the kinetic constraint includes the thermodynamic constraint.

This can be highlighted on a widely-used rate law, the reversible Michaelis-Menten kinetics [32]. In the simple case of a reaction *i* where the enzyme can only exist in one of three distinct states: free, all substrates bound, or all products bound, it can be written as:

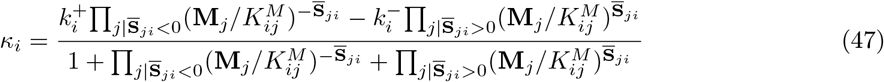

where 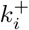 and 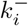 are the maximal forward and backward rates in reaction *i* per unit of enzyme and the 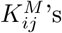 are the Michaelis constants. Equating the numerator to zero at equilibrium, we obtain: 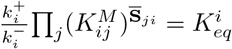. This gives:

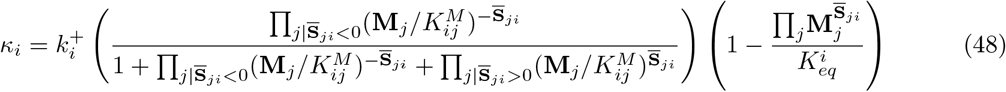

that is, the product of three terms: the positive capacity term per unit of enzyme, the positive (smaller than one) fractional saturation term depending on **M** and the thermodynamic term, which can be rewritten as 1 – *e*^Δ*_r_G_i_/RT*^ and gives the sign of *κ_i_* (and thus the sign of **v**_*i*_): *κ_i_* > 0 ⇔ Δ*_r_G_i_* < 0, i.e., the thermodynamic constraint. Thus *sign*(***κ***(**M**)) = **ts_M_** and we will assume that this equality holds for all kinetic laws we consider.

Note that, for given metabolite and enzyme concentrations **M** and **E**, the kinetic constraint **KC_E,M_** defines completely and uniquely the only vector that satisfies it: *Sol*_**KC_E,M_**_ = {**E** o ***κ***(**M**)}.

#### 2.1.3 Regulatory constraints

Coupling metabolic networks with Boolean transcriptional regulatory networks allows us to express the additional constraints imposed by gene regulatory information on a metabolic network and to take them into account when computing EFMs [20, 21, 19]. In all generality, such a constraint may be given by an arbitrary Boolean formula in terms of the reactions *i*, viewed as propositional symbols, i.e., the positive literal *i* meaning that reaction *i* is active (nonzero flux) and the negative literal ¬*i* meaning it is inactive (zero flux). Thus, when applying this Boolean constraint, noted *Bc*, to a flux vector **v**, the positive literal *i* is interpreted as **v**_*i*_ = 0, i.e., *i* ∈ *supp*(**v**) (4), and the negative literal ¬*i* as **v**_*i*_ = 0, i.e., *i* ∉ *supp*(**v**).

The regulatory constraint **RC**_*Bc*_ for **v** induced by *Bc* can thus be defined as:

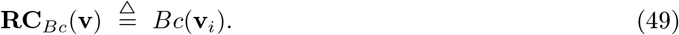

Note that coupled reactions as used by Flux Coupling Analysis (FCA) [5, 23, 36] can be easily represented by such constraints. For example, *i* directionally coupled to *j*, meaning that zero flux through *i* implies zero flux through *j*, is expressed by *Bc* = *i* ∨ ¬*j*, and *i* partially coupled to *j*, meaning that zero flux through *i* is equivalent to zero flux through *j*, is expressed by *Bc* = (*i* ∧ *j*) ∨ (¬*i* ∧ ¬*j*).

By rewriting the Boolean constraint *Bc* in DNF (Disjunctive Normal Form), 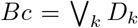, the set 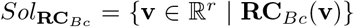 of vectors **v** satisfying the constraint **RC**_*Bc*_ is a union of the solution spaces for each disjunct *D_k_* (and this union can be assumed to be disjoint by taking the disjuncts *D_k_* two by two inconsistent). Now a disjunct is a conjunction of literals, where a negative literal ¬*i* corresponds to the constraint **v**_*i*_ = 0 and a positive literal *i* to the constraint **v**_*i*_ ≠ 0, which can be rewritten as the disjunctive constraint (**v**_*i*_ < 0) ∨ (**v**_*i*_ > 0). Finally, a propositional symbol *i* that does not appear in the disjunct corresponds to an absence of constraint on **v**_*i*_, which can be rewritten as the disjunction (**v**_*i*_ < 0) ∨ (**v**_*i*_ = 0) ∨ (**v**_*i*_ > 0). Thus, the solution space for a disjunct is itself the disjoint union of subspaces each one defined by constraints of type **v**_*i*_ *op_i_* 0 for all *i*, with *op_i_* ∈ {<, =, >}, that is, defined by 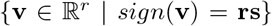 for a given sign vector **rs** ∈ {–, 0, +}^*r*^, i.e., an open orthant 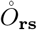 (which, for **rs** ≠ **0**, is topologically open in the vector subspace it spans and is the interior in this subspace of the closed orthant 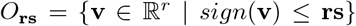, which is the closure of 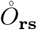 in 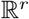). In summary, *Sol*_**RC**_*Bc*__ is thus the disjoint union of such open orthants. Now, instead of keeping this partition of *Sol*_**RC**_*Bc*__ in open orthants, it can be more practical to generalize this concept and deal with what we will call semi-open orthants *O*°, i.e., orthants *O* without some of their faces of lesser dimension (open orthant is thus a particular case of semi-open orthant, without any facet thus without any face of lesser dimension). To do this, we can group together with any given open orthant 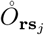 in *Sol*_**RC**_*Bc*__, in a same cluster 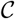, all other open orthants 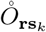 in *Sol*_**RC**_*Bc*__ such that *O*_**rs**_*k*__ ⊂ *O*_**rs**_*j*__, i.e. *O*_**rs**_*k*__ is a face of *O*_**rs**_*j*__, and, for any arbitrary orthant *O*_**rs**_*l*__ with *O*_**rs**_*k*__ ⊂ *O*_**rs**_*l*__ ⊂ *O*_**rs**_*j*__ (which is equivalent to **rs**_*k*_ < **rs**_*l*_ < **rs**_*j*_), then 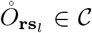. In this case 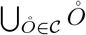 is a semi-open orthant *O*°. We can iteratively process like this by considering each time the open orthants in *Sol*_**RC**_*Bc*__ that have not yet been selected in any cluster, which guarantees that the semi-open orthants built in this way are disjoint. In addition, by choosing as open orthant 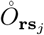, to start with at each iteration, a maximal one among those that remain, we are sure that no two of the semi-open orthants thus built can be grouped together to constitute a bigger semi-open orthant, i.e., the collection obtained is minimal in this sense. We finally obtain that *Sol*_**RC**_*Bc*__ can be written as a disjoint union of semi-open orthants, with no merging possible between any two of them. However note that such a decomposition is not unique and that an inclusion can still exist between the closures of two such semi-open orthants. If we consider the particular case of a Boolean constraint which is an arbitrary disjunct D, then for any closed *r*-orthant *O*_**η**_, i.e., for any full support sign vector **η**, *Sol*_**RC**_*D*_ ≤_**η**__ is a semi-open orthant, which is the face of *O*_**η**_ defined by **v**_*i*_ = 0 for all *i* such that ¬*i* is a literal of *D*, without its facets of equation **v**_*i*_ = 0 for all *i* such that *i* is a literal of *D*.

##### Lemma 2.2.

*The set Sol_**RC**_Bc__ of vectors in* 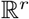 *satisfying the regulatory constraint **RC**_Bc_ is a disjoint union of open orthants* 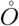. *These can be grouped together such that Sol_**RC**_Bc__ is a disjoint union of semiopen orthants O*°, *i.e., orthants O without some of their faces, such that no two of them can be grouped together to constitute a bigger semi-open orthant. If Bc is a conjunction of literals and* **η** *an arbitrary full support sign vector, then Sol_**RC**_Bc_ ≤_**η**__ is itself a semi-open orthant*.

### 2.2 Characterizing the solution space

We start from a flux cone (6) (or more generally a flux polyhedron (24)) representing the flux vectors of a metabolic network in a steady state, satisfying the stoichiometric equations, the inequalities regarding irreversibility of reactions (and possibly some inhomogeneous linear constraints). Then we consider additional biological constraints, such as those described in section (2.1). In all generality, these constraints will be noted **C_x_**(**v**), where **v** represents a flux vector in 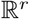 and **x** a vector of biochemical quantities involved in the constraint (typically, metabolite concentrations and enzyme concentrations). Some parameters (such as stoichiometric coefficients), that are not made explicit in the notation for sake of simplicity, are also present. In the following, the solution space in 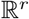 of an arbitrary constraint **C** will be noted *Sol*_**C**_, defined as: 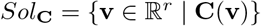. For given values of quantities **x**, the solution space is thus the constrained flux cone subset (not necessarily a cone, depending on **C_x_**):

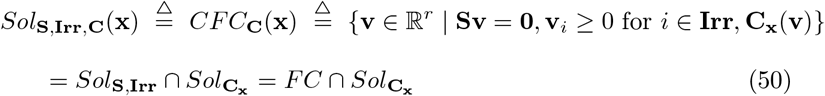

or, more generally, the constrained flux polyhedron subset (not necessarily a polyhedron, depending on **C_x_**):

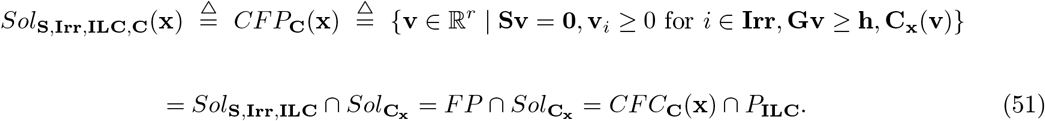

Now, very often, most of the values of quantities **x**, if not all, are unknown. We then only require consistency of the set of constraints **C_x_**, i.e., the existence of values for variables **x** such that the constraints are satisfied. This means that we replace the constraint **C_x_** by the constraint ∃**xC_x_** whose solution space is 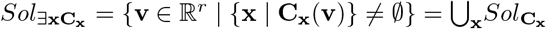. The solution space becomes:

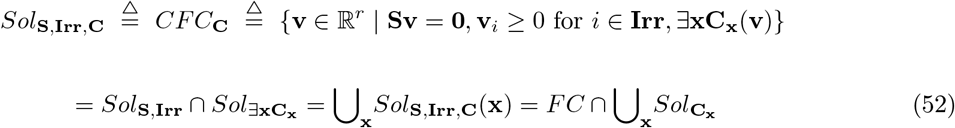

or:

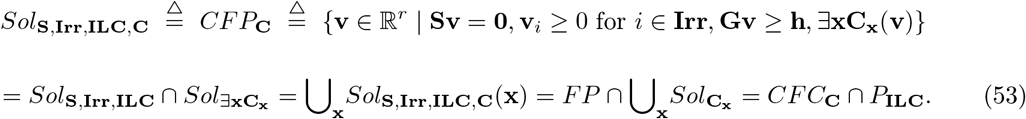

If some but not all of the values of quantities **x** are known, ∃**x** concerns only those **x** whose value is unknown, the known values being integrated in **C** as parameters to simplify the notations.

We will obtain examples of such constraints ∃**xC_x_**(**v**) from the cases of thermodynamic constraint **TC** and kinetic constraint **KC**.

Depending on the constraint considered, the solution space is in general no longer a cone or polyhedron and no longer convex. Nevertheless, the definitions of extreme vectors and extreme points, of elementary flux vectors and elementary flux points, and of elementary flux modes are still valid, as, respectively, non-decomposable and convex non-decomposable vectors, conformally non-decomposable and convex-conformally non-decomposable vectors, and support-minimal vectors, where decomposition and support minimality have to be understood w.r.t. vectors of the solution space. It is then clear that if a vector of *FC* (resp., *FP*) satisfies one of those properties in *FC* (resp., *FP*) and satisfies the given constraint, then it satisfies the same property in the solution space, i.e.:

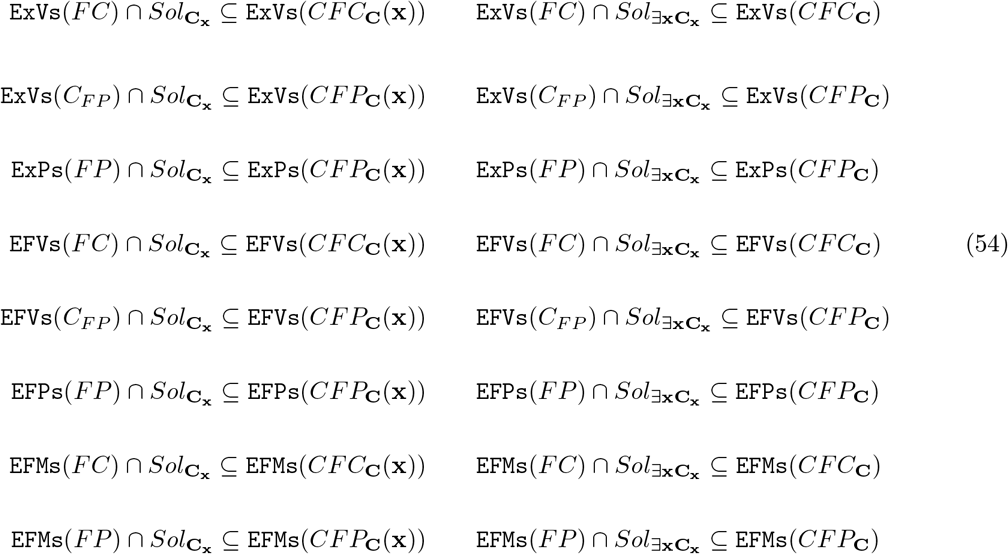

For EFMs, the above formulas are valid whatever the structure of *Sol*_**C**_. It is also the case for ExPs and EFPs but, in practice, only really meaningful if *CFP*_**C**_(**x**) (resp., *CFP*_**C**_) is a convex polyhedron. Finally, for ExVs and EFVs, this is only meaningful if *CFC*_**C**_(**x**) (resp., *CFC*_**C**_) is a convex polyhedral cone and *CFP*_**C**_(**x**) (resp., *CFP*_**C**_) is a convex polyhedron. This is the case when *Sol*_**C_x_**_ (resp., *Sol*_∃**xC_x_**_) is itself a convex polyhedral cone and we will see that, for almost all common biological constraints, this solution space is actually a union of such cones or of semi-open cones and thus the formulas will apply for each conical component. In this case we have 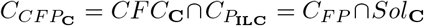. This can be generalized when *Sol*_**C**_ is a convex polyhedron, and thus *CFC*_**C**_ and *CFP*_**C**_ too, with *C*_*CFC*_**C**__ = *FC* ∩ *C*_*Sol*_**C**__ and *C*_*CFP*_**C**__ = *C*_*CFC*_**C**__ ∩ *C*_*P*_ILC__ = *C_FP_* ∩ *C*_*Sol*_**C**__, by replacing *Sol*_**C**_ by *C*_*Sol*_**C**__ in the formulas above regarding ExVs and EFVs.

However, in all the above cases, we generally do not have the reciprocal subset inclusion. This is what we will study for particular constraints.

#### 2.2.1 Application to thermodynamics

Assume first that the concentrations of the metabolites (resp., of the internal metabolites) are known and given. From (50), (51) and Lemma 2.1, we obtain:

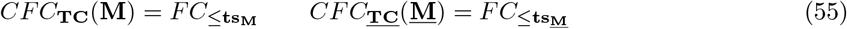

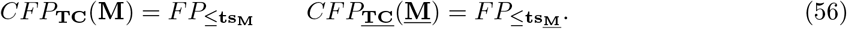

*CFC*_**TC**_(**M**), *CFC*_**<u>TC</u>**_(**<u>M</u>**) are thus flux cones and *CFP*_**TC**_(**M**), *CFP*_**<u>TC</u>**_(**<u>M</u>**) flux polyhedra (possibly equal to {**0**} or empty (for polyhedra) if the flux directions imposed by the concentrations of the metabolites and by the second law of thermodynamics are incompatible with the steady-state assumption). When nonempty, they consist of a single flux tope. From (18), (35) and (38), it follows that:

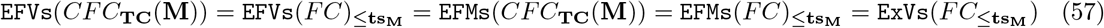

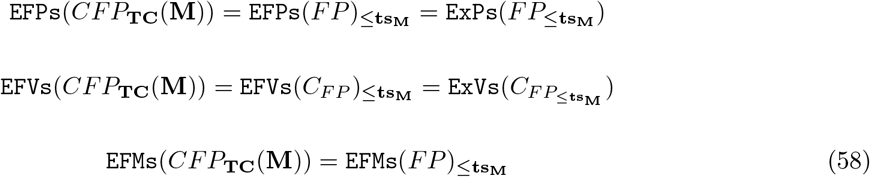

and the same for *CFC*_**<u>TC</u>**_(**<u>M</u>**) and *CFP*_**<u>TC</u>**_(**<u>M</u>**). Now, considering the decomposition of *FC* (resp., *FP*) into flux topes, we can proceed flux tope by flux tope for the computation of the sets above. Starting from a FT *FC*_≤**η**_ (resp., *FP*_≤**η**_), we have (*FC*_≤**η**_)_≤**ts_M_**_ = *FC*_≤**·**_ (resp., (*FP*_≤**η**_)_≤**ts_M_**_ = *FP*_≤**η′**_) with **η′** = *inf* (**η, ts_M_**) (where *inf* is defined component-wise with *inf* (–, +) = 0) and the four sets above are equal to ExVs(*FC*_≤**η**′_), ExPs(*FP*_≤**η**′_), ExVs(*C*_*FP*_≤**η**__) and EFMs(*FP*_≤**η**′_), respectively. This is in particular the case if we split each reversible reaction into two irreversible ones as, in higher dimension, 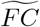 (resp., 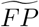) is reduced to a single flux tope defined by **η** = +, the all-positive sign vector, and thus **η′** is obtained from **ts_M_** by changing each – into 0. Note also that *FC* (resp., *FP*) consistent does not imply that *CFC*_**TC**_(**M**) (resp., *CFP*_**TC**_(**M**)) is consistent (**η** being full support does not imply that **η′** is full support).

##### Proposition 2.3.

*Given metabolite concentrations **M**, the space of flux vectors in FC (resp., FP) satisfying the thermodynamic constraint **TC_M_** is a flux cone (resp., a flux polyhedron), made up of vectors of FC (resp., FP) which conform to the thermodynamic sign vector **ts_M_** given in* (43). *Its elementary flux vectors, identical to elementary flux modes, (resp., elementary flux points, elementary flux vectors and elementary flux modes) are exactly those of FC (resp., FP) that satisfy the constraint, i.e., that conform to **ts_M_**. The same result holds for internal metabolite concentrations **<u>M</u>** and thermodynamic constraint **<u>TC_M_</u>** with **ts_<u>M</u>_***.

Let’s now consider the more usual case where the concentrations of the metabolites (at least those of internal metabolites) are unknown. From (44), we obtain one or the other form for the thermodynamic constraint (existentially quantified on metabolite concentrations):

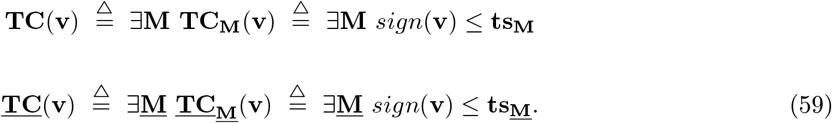

Though the metabolite concentrations **M**_*j*_ are unknown, some lower bounds 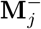 and upper bounds 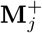 on these concentrations are often known. They are thus added as additional constraints:

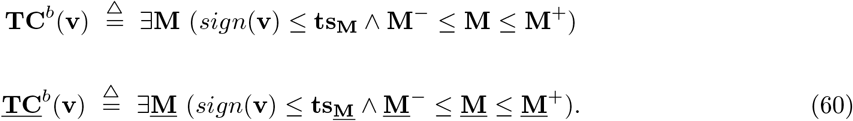

The solution space in 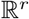 of these constraints is thus 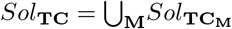 (idem for **<u>TC</u>**) and 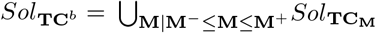 (idem for **<u>TC</u>**^*b*^). From the result of Lemma 2.1, it follows: 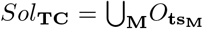 and 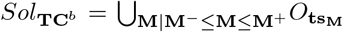 (idem for **<u>TC</u>** and **<u>TC</u>**^*b*^ with *O*_**ts_<u>M</u>_**_). This is obviously enough to take the union on the maximal **ts_M_**’s (resp., **ts_<u>M</u>_**’s) when **M** varies. As these are at most 2^*r*^, the union is finite.

##### Lemma 2.4.

*The set Sol_**TC**_ (resp., Sol_**TC**^b^_) of vectors in* 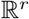 *satisfying the thermodynamic constraint **TC** (resp., **TC**^b^) is a union of closed orthants. More precisely, it is the (finite) union for all **M** (resp., all bounded **M**) of the sets of vectors that conform to **ts_M_**, i.e., O_**ts**_**M**__’s. The same result holds for **<u>TC</u>** and **<u>TC</u>**^b^ with **<u>M</u>** and O_ts_<u>M</u>__*.

It follows from (52), (53), (55), (56) that:

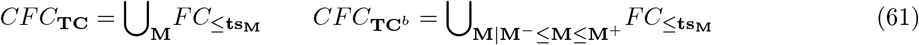

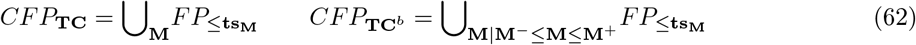

and the same for **<u>TC</u>** and **<u>TC</u>**^*b*^ with **<u>M</u>** and **ts_<u>M</u>_**, where all unions are finite. From this and (18), (35) and (38), we get:

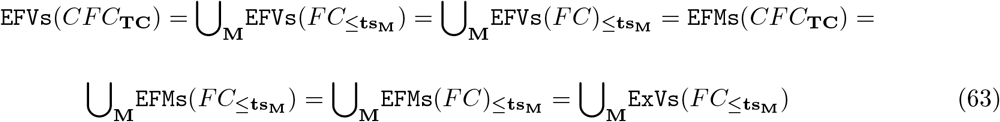

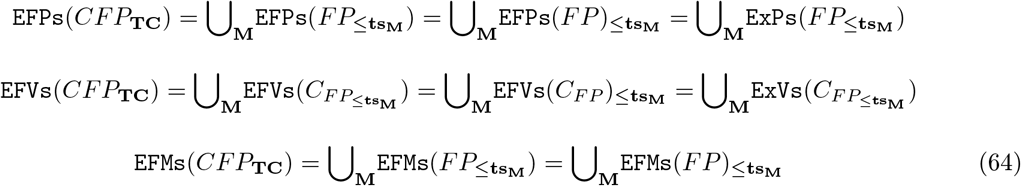

and the analog for **TC**^*b*^, **<u>TC</u>** and **<u>TC</u>**^*b*^. Now, considering the decomposition of *FC* (resp., *FP*) into flux topes, we can proceed flux tope by flux tope for the computation of the sets above. Starting from a FT *FC*_≤**η**_ (resp., *FP*_≤**η**_), we have 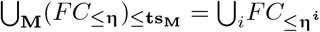 (resp., 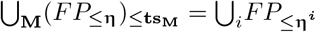) where {**η**^*i*^} are the maximal sign vectors in 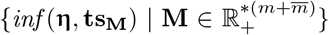 and the four sets above are equal to 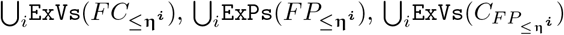 and 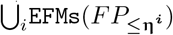, respectively. This is in particular the case if we split each reversible reaction into two irreversible ones as, in higher dimension, 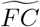 (resp., 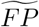) is reduced to a single flux tope defined by **η** = +. The analog holds for **TC**^*b*^, **<u>TC</u>** and **<u>TC</u>**^*b*^.

##### Proposition 2.5.

*The space of flux vectors in FC (resp., FP) satisfying the thermodynamic constraint **TC**, or **TC**^b^, is a finite union of flux cones (resp., flux polyhedra), obtained as those vectors of FC (resp., FP) which conform to **ts_M_**, for a certain **M**, or bounded **M**. It is thus no longer convex but made up of particular faces FC_≤**η**^i^_ (resp., FP_≤**η**^i^_) of each flux tope FC_≤**η**_ for FC (resp., FP_≤**η**_ for FP), with* **η**^*i*^ ≤ **η**. *Its elementary flux vectors, identical to elementary flux modes, (resp., elementary flux points, elementary flux vectors and elementary flux modes) are exactly those of *FC* (resp., FP) that satisfy the constraint, i.e., that conform to a certain **ts_M_**, and coincide thus with the extreme vectors (resp., the extreme points, extreme vectors and elementary modes) of the FC_≤**η**^i^_’s (resp., FP_≤**η**^i^_’s). The same result holds for **<u>TC</u>**, or **<u>TC</u>**^b^, with **<u>M</u>** and **ts**_<u>M</u>_*.

We will now refine this result in order to characterize the faces involved. Consider the case of a flux cone where the concentrations of external metabolites are given and assume first there are no bounds on the concentrations of the internal metabolites [33]. As 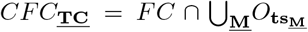 and 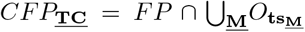, let’s begin by studying 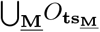. We have: 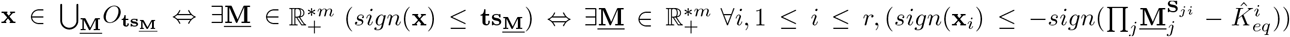 from (43). By applying the monotonic logarithm function and noting 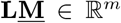 the vector whose components are given by **L<u>M</u>**_*j*_ = *ln*(**<u>M</u>**_*j*_), we obtain: 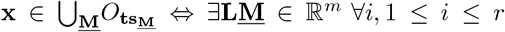, 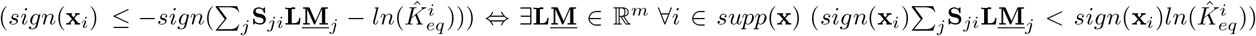. We will now make use of Gale’s theorem (or Kuhn-Fourier theorem), which is a form of Farkas’ duality lemma (for two vectors **x, y, x** < **y** means **x**_*i*_ < **y**_*i*_ for all *i*):

**Gale’s theorem.** *For any* 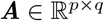 *and* 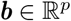, *exactly one of the following statements holds:*

a. *there exists* 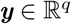 *such that **Ay** < **b**;*
b. *there exists* 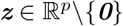 *such that **z** ≥ **0, z**^T^ **A** = **0** and **z**^T^**b*** ≤ 0.

Applying this theorem in the formula above to 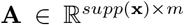 (where we note 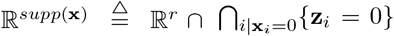 the subspace of vectors having a null component outside the support of **x**), given by **A**_*ij*_ = *sign*(**x**_*i*_)**S**_*ji*_, and 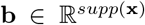, given by 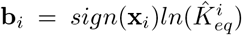, provides: 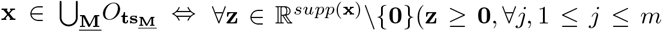, 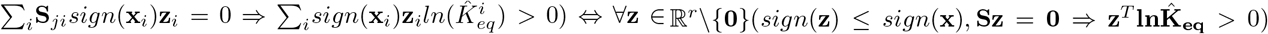 where 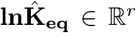 is defined by: 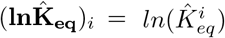.

We thus obtain: 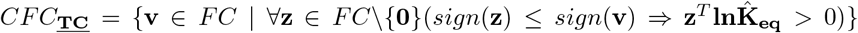. This can be rewritten as: 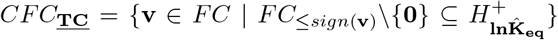, where 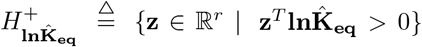 is an open half-space with the hyperplane 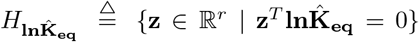 as a frontier. Considering the decomposition of *FC* into FTs *FC*_≤**η**_, this is equivalent to: 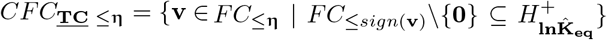 for all **η**’s maximal sign vectors in *sign*(*FC*). Now, note that, in this formula, *FC*_≤*sign*(**v**_ is the minimal face (for set inclusion) of *FC*_≤**η**_ containing **v**. We finally obtain finally:

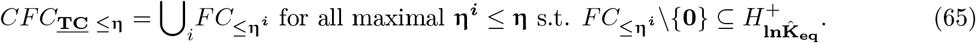

That is to say *CFC*_**<u>TC</u>**_ is the union of all the maximal faces *FC*_≤**η**_ of the FTs for *FC* such that 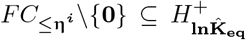 or, equivalently, the union of all maximal faces *F_i_* of the *FC* ∩ *O*’s for all *r*-orthants *O* such that 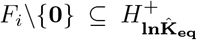. A topological consequence of this characterization is that *CFC*_**<u>TC</u>**_\{**0**} is a connected set (actually arc-connected). Another consequence, regarding EFMs (or, equivalently, EFVs), is:

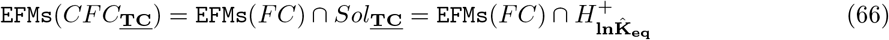

i.e., the elementary flux modes in the (non-convex) cone of those flux vectors in *FC* satisfying the thermodynamic constraint **<u>TC</u>** are exactly those elementary flux modes in *FC* that satisfy **<u>TC</u>**, i.e., that belong to the open half-space 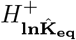. We can equivalently write:

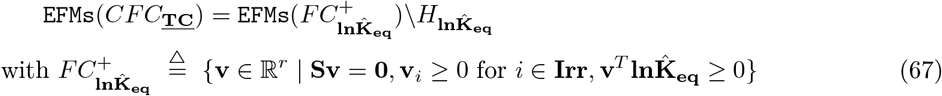

where 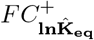 is the polyhedral cone obtained from *FC* merely by adding the homogeneous linear inequality 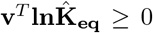 [34]. Each flux vector of *CFC*_<u>**TC**</u>_ is a conformal conical sum of these EFMs, but the converse is false as *CFC*_**<u>TC</u>**_ is not convex. Thus the set of EFMs, considered globally, does not characterize the solution space *CFC*_**<u>TC</u>**_. More precisely, we have:

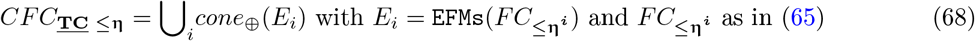

i.e., *CFC*_**<u>TC</u>**_ is characterized by the decomposition of the set of EFMs into the (non-disjoint) subsets *E_i_*. Now, the *E_i_*’s are exactly the maximal subsets of EFMs included in a given flux tope for *FC* (i.e., in a given *r*-orthant *O*) and whose conical hull is included in *CFC*_**<u>TC</u>**_, i.e., all vectors in this hull must satisfy the constraint **<u>TC</u>**: *E_i_* maximal such that *E_i_* ⊆ EFMs(*CFC*_**<u>TC</u>**_) and *E_i_* ⊆ *O* with *O r*-orthant and *cone*_⊕_(*E_i_*) ⊅ *CFC*_**<u>TC</u>**_. Such an *E_i_* is called a largest thermodynamically consistent set (LTCS) of EFMs [10] in *O* (or, equivalently, in the flux tope *FC*_≤**η**_ = *FC* ∩ *O* defined by *O*).

We could want to estimate the ratio of thermodynamically feasible EFMs on all EFMs, i.e., the ratio of EFMs(*CFC*_**<u>TC</u>**_) on EFMs(*FC*). If all reactions are reversible (*r_I_* = 0), then the function **v** ↦ –**v** maps the EFMs on one side of hyperplane 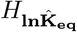 onto the EFMs on the other side. Thus, if we neglect the EFMs that might belong to this hyperplane, it means that 50% of the EFMs are thermodynamically feasible (and thus only 50% eliminated). Now, intuitively, irreversible reactions given in **Irr** come from an expert knowledge that can be seen as a form of compiled thermodynamic knowledge, as we saw that it is thermodynamics which imposes the direction in which a reaction may proceed. So, we can assume, provided the adequacy of the model for some given environment (such as the concentrations of external metabolites, supposed here to be known), that any thermodynamically feasible EFM satisfies these irreversibility constraints. Under this assumption, the irreversibility constraints rule out only thermodynamically unfeasible EFMs so if we then apply the thermodynamic constraint **<u>TC</u>** we only eliminate at most than 50% of the remaining EFMs (from 50% when all reactions are reversible to 0% when all are irreversible, without splitting any reversible reaction into two irreversible ones).

If inhomogeneous linear constraints are added, we have *CFP*_**<u>TC</u>**_ = *CFC*_**<u>TC</u>**_ ∩ *P*_**ILC**_, with 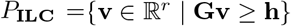 (24) and we obtain in the same way, for all **η**’s maximal sign vectors in *sign* (*FC*):

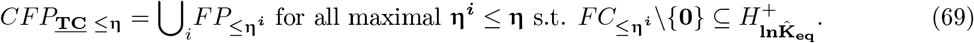

That is to say *CFP*_**<u>TC</u>**_ is the union of all the *FP*_≤**η**^*i*^_ = *FC*_≤**η**^*i*^_ ∩ *P*_**ILC**_ such that *FC*_≤**η**^*i*^_ is a maximal face of a FT for *FC* verifying 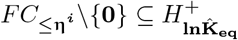 or, equivalently, the union of the *F_i_* ∩ *P*_**ILC**_ for all maximal faces *F_i_* of the *FC*∩*O*’s for all *r*-orthants *O* such that 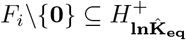. Take care because the *FP*_≤**η**^*i*^_’s involved are faces of *FP*_≤**η**_ included in 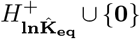, but not any face of *FP*_≤**η**_ included in 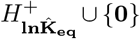 is included in *CFP*_**<u>TC</u>**_ (as such a face may be defined by inhomogeneous equality constraints coming from **ILC** and not only by nullity constraints of the form **v**_*i*_ = 0). *CFP*_<u>**TC**</u>_ is no longer a connected set in general.

We can sum up these results as follows.

##### Proposition 2.6.

*The space of flux vectors in FC satisfying the thermodynamic constraint **<u>TC</u>** is a finite union of flux cones, obtained as all the maximal faces of all the flux topes FC*_≤**η**_ *that are entirely contained (except the null vector) in the open half-space* 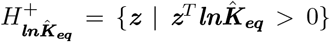. *The thermodynamically feasible* EFMs *are thus those* EFMs *which belong to this half-space and can be simply computed as the* EFMs *of the flux cone obtained from FC by adding to it the homogeneous linear inequality* 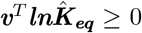 *(and removing those* EFMs *that would belong to the frontier hyperplane of* 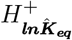 *). These* EFMs *can be decomposed into (non-disjoint) maximal subsets of* EFMs *belonging to a same flux tope (i.e., a same r-orthant) and whose conical hull is made up of thermodynamically feasible vectors, each of these subsets thus representing the set of* EFMs *of one of the maximal faces above. At most 50% of the* EFMs *can thus be ruled out as thermodynamically infeasible, if we assume that no thermodynamically feasible flux vector violates given irreversibility constraints. In the presence of additional inhomogeneous linear constraints on flux vectors given by **Gv** ≥ **h**, the space of flux vectors in FP satisfying **<u>TC</u>** is a finite union of flux polyhedra, obtained as intersections of the flux cones above with the polyhedron defined by **Gv** ≥ **h**. All these results hold for constraint **TC** by just replacing the* 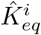*’s by the* 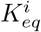*’s*.

This applies in particular to 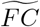 and 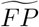 (after splitting each reversible reaction into two irreversible ones) with the simplification, as 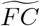 (resp., 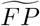) is reduced to a single flux tope defined by **η** = +, that we must just consider the maximal faces of 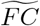 that are entirely contained (except the null vector) in the open half-space 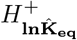.

Consider now the case where certain bounds on the concentrations of internal metabolites are known, i.e., the case of the thermodynamic constraint **TC**^*b*^. We have 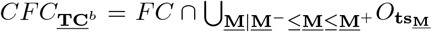. From what precedes, we obtain: 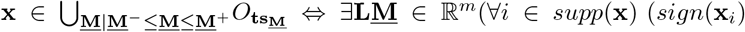 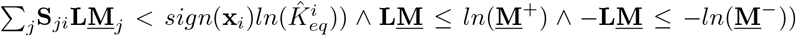. Applying Gale’s theorem in this formula to 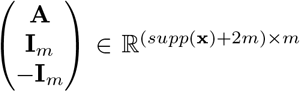, with **A**_*ij*_ = *sign*(**x**_*i*_)**S**_*ji*_, and 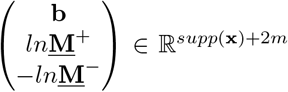, with 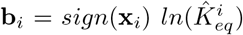, provides: 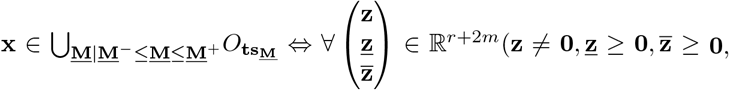 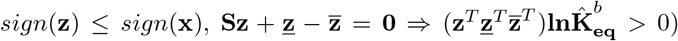 where 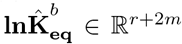 is defined by: 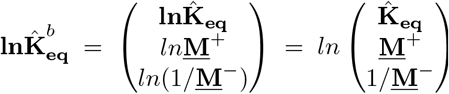. Let 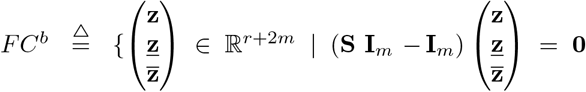, **z**_*i*_ ≥ 0 for 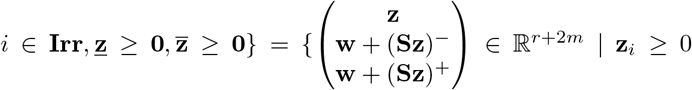 for *i* ∈ **Irr, w** ≥ **0**}, where 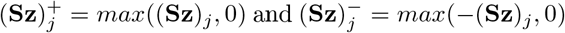. *FC* is a flux cone in 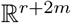 of dimension *r+m* and 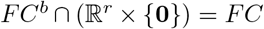. We thus obtain: 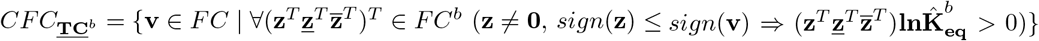. This is equivalent to: 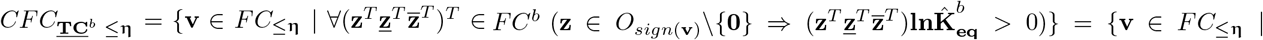 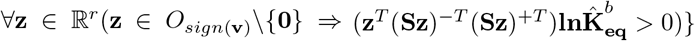 for all flux topes *FC*_≤**η**_ for *FC*. This can be rewritten as: 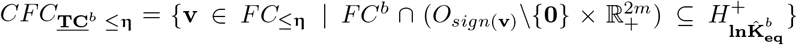, where we note 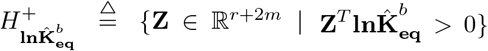 the open half-space with hyperplane 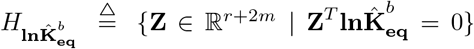 as a frontier. We have 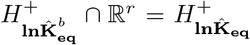 and 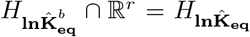. Now, note that, in the formula above, 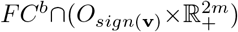 is a face of the cone 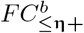 (we note **η**+ the sign vector of dimension *r*+2*m* obtained by concatenation of **η** of dimension *r* and + of dimension 2*m*, thus 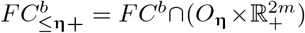) whose intersection with 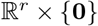 is equal to *FC*_≤*sign*(**v**)_, the minimal face of the tope *FC*_≤**η**_ containing **v**. Actually, among all the faces of 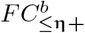 whose intersection with 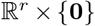 is equal to *FC*_≤*sign*(**v**)_, this is the maximal one (without any constraint on the 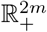 factor). Thus finally:

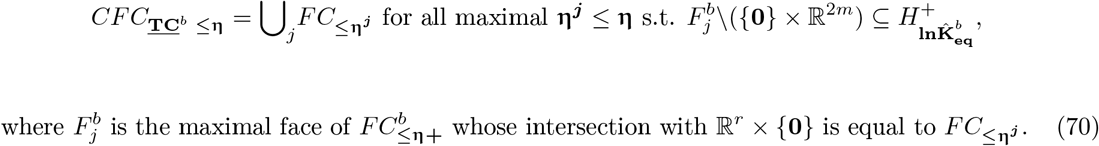

Note that any *FC*_≤**η**^*j*^_ given by (70) is a face of a certain *FC*_≤**η**^*i*^_ given by (65) (i.e., ∀**η^*j*^**∃**η^*i*^ η^*j*^** ≤ **η^*i*^**). *CFC*_<u>**TC**</u>^*b*^_\{**0**} is no longer a connected set in general. A consequence, for EFMs (or, equivalently, EFVs), is:

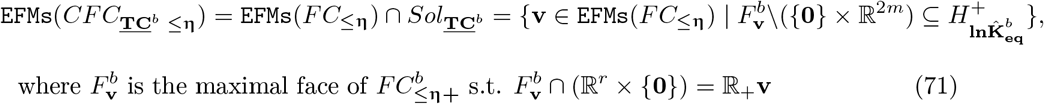

i.e., the elementary flux modes in the (non-convex) cone of those flux vectors in *FC*_≤**η**_ satisfying the thermodynamic constraint **<u>TC</u>**^*b*^ are exactly those elementary flux modes **v** in *FC*_≤**η**_ (or in *CFC*_**<u>TC</u>** ≤**η**_) that satisfy **<u>TC</u>**^*b*^, i.e., such that the maximal face of 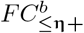 whose intersection with 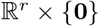 is equal to the ray 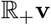 is included in 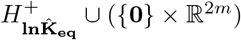. Note that EFMs(*CFC*_**<u>TC</u>**^*b*^_) is thus given by:

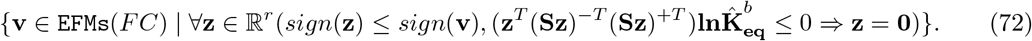

Each flux vector of *CFC*_**<u>TC</u>**^*b*^_ is a conformal conical sum of these EFMs, but the converse is false as *CFC*_**<u>TC</u>**^*b*^_ is not convex. More precisely, we have:

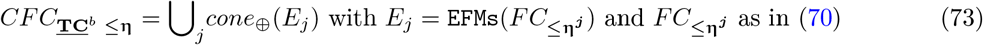

i.e., *CFC*_**<u>TC</u>**^*b*^_ is characterized by the decomposition of the set of EFMs into the (non-disjoint) subsets *E_j_*. Now, the *E_j_*’s are exactly the maximal subsets of EFMs included in a given flux tope for *FC* (i.e., in a given *r*-orthant *O*) and whose conical hull is included in *CFC*_**<u>TC</u>**^*b*^_, i.e., all vectors in this hull must satisfy the constraint **<u>TC</u>**^*b*^: *E_j_* maximal such that *E_j_* ⊅ EFMs(*CFC*_**<u>TC</u>**^*b*^_) and *E_j_* ⊅ *O* with *O r*-orthant and *cone*_⊕_(*E_j_*) ⊅ *CFC*_**<u>TC</u>**^*b*^_. Such an *E_j_* is called a largest thermodynamically consistent (for bounded concentrations of internal metabolites) set (LTCbS) of EFMs [10] in *O* (or, equivalently, in the flux tope *FC*_≤**η**_ = *FC* ∩ *O* defined by *O*) and is included in a certain LTCS *E_i_* as in (68).

If inhomogeneous linear constraints **ILC** are added, we have, for all **η**’s maximal sign vectors in *sign*(*FC*):

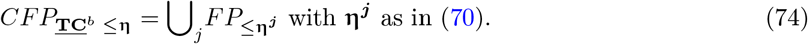

That is to say *CFP*_**<u>TC</u>**^*b*^_ is the union of all the *FP*_≤**η^*j*^**_ = *FC*_≤**η^*j*^**_ ∩ *P*_**ILC**_ such that *FC*_≤**η^*j*^**_ is given as in (70).

We can sum up these results as follows.

##### Proposition 2.7.

*For FC a flux cone in* 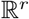, *let’s define its lift to* 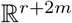 *as the flux cone* 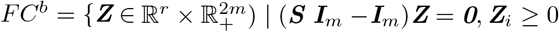 *for i ∈ **Irr***, 1 ≤ *i* ≤ *r*}. *Thus*, 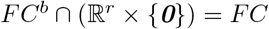. *For any flux tope FC*_≤**η**_ *for FC and any face FC′ of FC*_≤**η**_, *its lift to* 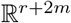 *is defined as the maximal face of* 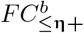 *whose intersection with* 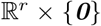 *is equal to FC′. The space of flux vectors in FC satisfying the thermodynamic constraint **<u>TC</u>**^b^ is a finite union of flux cones, obtained as all the maximal faces of all the flux topes FC*_≤**η**_ *whose lifts to* 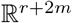 *are entirely contained (except vectors from* 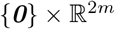*) in the open half-space* 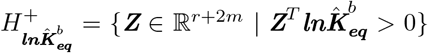, *where* 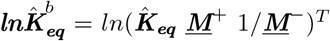. *The thermodynamically feasible* EFMs *in FC*_≤**η**_ *are those* EFMs *in FC*_≤**η**_ *whose lifts (as rays) are contained (except vectors from* 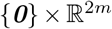*) in this half-space. They can be decomposed into (non-disjoint) maximal subsets of* EFMs *belonging to a same flux tope (i.e., a same r-orthant) and whose conical hull is made up of thermodynamically feasible vectors, each of these subsets representing thus the set of* EFMs *of one of the maximal faces above. In the presence of additional inhomogeneous linear constraints on flux vectors given by **Gv** ≥ **h**, the space of flux vectors in FP satisfying **<u>TC</u>**^b^ is a finite union of flux polyhedra, obtained as intersections of the flux cones above with the polyhedron defined by **Gv** ≥ **h***.

#### 2.2.2 Application to kinetics

In the same way, we obtain for the kinetic constraint in the absence of knowledge regarding values of concentrations of enzymes and metabolites:

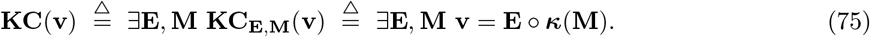

Once more we can add optional lower and upper bounds 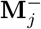 and 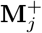 on metabolite concentrations and/or lower and upper bounds 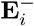 and 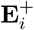 on enzyme concentrations if they are known, and also a global enzymatic resource constraint, which is often considered, as **c**^*T*^**E** ≤ *W*, where **c** is a constant positive vector of size *r* and *W* a positive constant:

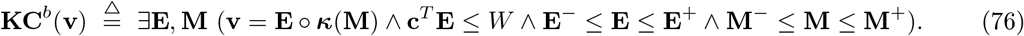

We can also consider intermediate constraints, existentially quantified only on metabolite concentrations if enzyme concentrations are known:

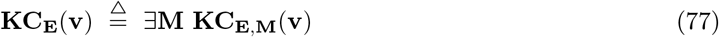

possibly with bounds on metabolite concentrations in the quantification:

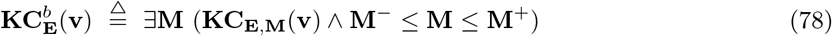

or only on enzyme concentrations if metabolite concentrations are known:

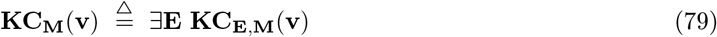

possibly with enzymatic resource constraint and bounds on enzyme concentrations in the quantification:

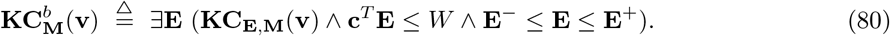

The solution space in 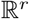 of the constraint **KC_M_** is 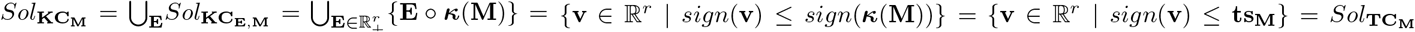, as the sign of ***κ***(**M**) is the thermodynamic sign vector **ts_M_**. It is thus the same as the solution space of the thermodynamic constraint **TC_M_** and is the closed orthant *O*_**tS_M_**_. It follows that the solution space of the constraint **KC**, given by 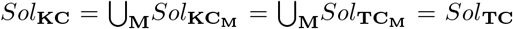, is the same as the solution space of the thermodynamic constraint **TC** and is the finite union of the closed orthants *O*_**ts_M_**_. Similarly, *Sol*_**KC**^*b*^_ = *Sol*_**TC**^*b*^_ if the bounds are only on the metabolite concentrations, i.e., there are no bounds on enzyme concentrations. From 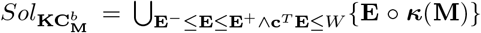, it follows that the flux vectors in 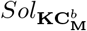 are defined by the linear inequalities: 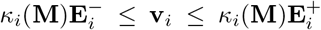 for those *i*’s such that *κ_i_*(**M**) > 0, 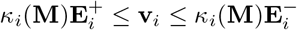 for those *i*’s such that *κ_i_*(**M**) < 0, **v**_*i*_ = 0 for those *i*’s such that *κ_i_*(**M**) = 0 and 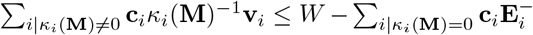. Thus 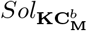 is a convex polyhedron (a parallelepiped contained in the closed orthant *O*_**tS_M_**_, truncated by a hyperplane). Now, if **E**^-^ = **0**, i.e., in the absence of positive lower bounds on enzyme concentrations, this polyhedron has **0** as a vertex and is nothing else than *Sol*_**KC_M_**_, i.e., the closed orthant *O*_**ts_M_**_, truncated as a parallelepiped by the faces given by 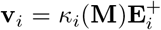 for those *i*’s such that *κ_i_*(**M**) ≠ 0 and by the hyperplane 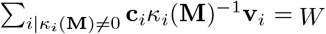. Consequently 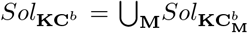 is equal to a certain truncation of *Sol*_**KC**_ = *Sol*_**TC**_, defined as the finite union of truncations of the closed orthants *O*_ts_M__ (each *O*_**ts**_ becoming a parallelepiped, after being truncated by hyperplanes according to equations 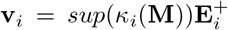 if **ts**_*i*_ = + (resp., 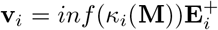 if **ts**_*i*_ = –), where *sup* (resp., *inf*) applies to those **M** such that *sign*(***κ***(**M**)) = **ts**, which gives a polyhedron, but also, in the presence of an enzymatic resource constraint, by an algebraic, non-linear, surface, which gives in this case a local solution space in *O*_**ts**_ that is no longer a polyhedron and is not necessarily convex).

##### Proposition 2.8.

*Given metabolite concentrations **M**, the kinetic constraint **KC_M_** is identical to the thermodynamic constraint **TC_M_** and thus the set Sol_**KC_M_**_ of vectors in* 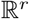 *satisfying **KC_M_** is the closed orthant defined by the thermodynamic sign vector **ts_M_**. The kinetic constraint **KC** is identical to the thermodynamic constraint **TC** and thus the set Sol_KC_ of vectors in* 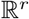 *satisfying **KC** is a union of closed orthants. In the presence of bounds only on metabolite concentrations (and not on enzyme concentrations), the kinetic constraint **KC**^b^ is identical to the thermodynamic constraint **TC**^b^ and thus the set Sol_KC^b^_ of vectors in* 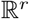 *satisfying **KC**^b^ is a union of closed orthants. So, these kinetic constraints boil down to thermodynamic constraints and the results regarding the geometrical structure of the corresponding spaces of flux vectors and the characterization of elementary flux vectors (or elementary flux modes) given by propositions 2.3, 2.5, 2.6 and 2.7 apply: in particular, CFC_KC_*(***M***) = *CFC_**TC**_*(***M***) *is a flux cone and CFC_**KC**_ = CFC_**TC**_ and CFC_KC^b^_ = CFC_**TC**^b^_ are finite unions of flux cones. For a given **M** and in the presence of bounds on enzyme concentrations, the set of vectors in* 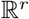 *satisfying* 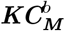 *is a convex polyhedron and CFC_**KC**^b^_* (***M***) *is thus a flux polyhedron. In the particular case where positive lower bounds on enzyme concentrations are absent, CFC_**KC**^b^_* (***M***) *is just the parallelepiped obtained by truncating the flux cone CFC_**KC**_*(***M***) = *CFC_**TC**_*(***M***) *by hyperplanes originating from the upper bounds on enzyme concentrations and by a hyperplane originating from the enzymatic resource constraint and coincides thus with the said flux cone in a certain adequate neighborhood of **0**, i.e., for sufficiently small values of the fluxes. In this case, CFC_**KC**^b^_ is thus a truncation (by an algebraic surface) of CFC_**TC**_ and coincides with this union of flux cones in a certain adequate neighborhood of **0** and the characterization of elementary flux vectors remains valid in this neighborhood. These results extend in the presence of additional inhomogeneous linear constraints on flux vectors given by **Gv** ≥ **h** by intersecting the solution spaces above with the polyhedron defined by **Gv** ≥ **h**, giving rise to flux polyhedra (results in a neighborhood of **0** holding only if **h** < **0**)*.

However the geometric structure of *Sol*_**KC**_*E*__, of *Sol*_**KC**^*b*^_ and of *Sol*_**KC**^*b*^_ in the presence of positive lower bounds on enzyme concentrations, depends on the kinetic function ***κ***(**M**) and *CFC*_**KC**_(**E**), *CFC*_**KC**^*b*^_ (**E**) and *CFC*_**KC**^*b*^_ are generally neither polyhedra nor convex.

Proposition 2.8 has important consequences on constrained enzyme allocation problems in kinetic metabolic networks. Considering a kinetic metabolic network, with possible bounds on metabolite concentrations, but not on enzyme concentrations, i.e., with solution space *CFC*_**KC**_ = *CFC*_**TC**_, or *CFC*_**KC**^*b*^_ = *CFC*_**TC**^*b*^_, which is a finite union, for **M** varying, of the flux cones *FC*_≤**ts_M_**_, the generic enzyme allocation problem consists in maximizing the specific flux (or specific rate, i.e., rate expressed per unit of biomass amount) of a given reaction, say *k*, defined as **v**_*k*_/*E_T_*, where **v** is a flux vector in *CFC*_**KC**_ or *CFC*_**KC**^*b*^_ and *E_T_* denotes the total protein content in the system. *E_T_* is expressed in all generality as a fixed weighted sum of the enzyme concentrations, 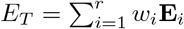 (the *w_i_*’s being given positive weights that denote the fraction of the resource used per unit of enzyme), able to encode different enzymatic constraints (such as limited cellular or membrane surface space) as well as other constraints regarding the abundance of certain enzymes. Likewise, the steady-state flux component **v**_*k*_ > 0 may stand for diverse metabolic processes, ranging from the synthesis rate of a particular product within a specific pathway to the rate of overall cellular growth. The formation rate of a metabolic product expressed per gram of biomass and the specific growth rate of a cell are both examples of such specific rates. We look for maximization in the solution space by varying the metabolite concentrations (inside their bounds, if any) and the enzyme concentrations (without bounds), which gives rise to a complex non-convex optimization problem. Now, maximizing **v**_*k*_/*E_T_* is equivalent to fixing the rate **v**_*k*_ to a positive value, e.g., to 1, and minimizing the *E_T_* needed to attain this level of **v**_*k*_. This means minimizing the function 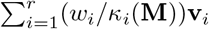 by varying **M** (with possible bounds) and, for each given **M**, the flux vector **v** in *FP*_**M**_ = *FC*_≤**ts_M_**_ ∩ {**v** | **v**_*k*_ = 1} (without bounds on **v** as we assume there are no bounds on enzyme concentrations). If all *FP*_**M**_’s are empty, the problem is unsolvable, i.e., **v**_*k*_ > 0 is incompatible with the kinetics. Otherwise, i.e., when the problem is solvable, we consider successively each nonempty *FP*_**M**_. Such an *FP*_**M**_ is a flux polyhedron whose elementary flux points (which are equal to the extreme points or vertices) correspond to the intersections of the hyperplane {**v** | **v**_*k*_ = 1} with extreme rays (edges) of *FC*_≤**ts_M_**_, i.e., EFMs of *CFC*_**KC**_ or *CFC*_**KC**^*b*^_, and whose elementary vectors (equal to extreme vectors), if any, correspond to the extreme rays of *FC*_≤**ts_M_**_ ∩ {**v** | **v**_*k*_ = 0 }. As, for **M** fixed, the function to minimize is linear in **v**, its minimum on *FP*_**M**_ is reached on a lower-dimensional face of *FP*_**M**_ (as **v**_*i*_/*κ_i_*(**M**) ≥ 0, *w_i_* > 0 and *FP*_**M**_ is included in a closed orthant, this face is necessarily a convex hull of certain extreme points of *FP*_**M**_ even if it is not a polytope, i.e., no extreme vector may occur as one of the generators of this face), and thus reached in particular at at least one extreme point, i.e., at an EFM. Now, as the total number of EFMs is finite, so is the number of those for which the minimum of the function occurs for any given **M**, considering all nonempty *FP*_**M**_’s. So, when **M** varies in its domain, we obtain the result that the maximum of the specific flux **v**_*k*_/*E_T_* occurs (at least) at an EFM of *CFC*_**KC**_ or *CFC*_**KC**^*b*^_. In the case of *CFC*_**KC**^*b*^_ and assuming the *κ_i_*(**M**)’s are continuous, this maximum is attained at an EFM at finite metabolite concentrations as **M** then varies in the compact set 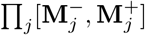. In the case of *CFC*_**KC**_, the maximum might not be attained at finite metabolite concentrations. This is the result already obtained in [31, 47] and we followed a similar proof, but relying this time on a precise characterization of the solution space *CFC*_**KC**_ or *CFC*_**KC**^*b*^_ and of the EFMs given by the Proposition 2.8, which was not the case in the above-quoted works. Finally, the enzyme allocation problem can theoretically be solved by computing all the thermodynamically feasible EFMs having *k* in their support (i.e., satisfying the Boolean constraint *Bc* = *k*, see next subsection), say {**e**^*l*^} (all components of which are fixed by 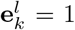), and, for each one, by computing the minimum (if it exists) of 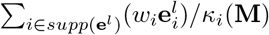 for **M** varying in its domain, such that 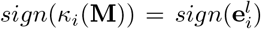 for all *i* ∈ *supp*(**e**^*l*^), which is a much simpler optimization problem than the initial one. The global minimum, if it exists, is then the smallest of these local minima, for all **e**^*l*^’s. We then obtain the maximum value of the specific flux **v**_*k*_/*E_T_*, an EFM where this maximum occurs and the values of the metabolite concentrations for which it occurs.

##### Proposition 2.9.

*Given a kinetic metabolic network with possible bounds on metabolite concentrations, but not on enzyme concentrations, i.e., with solution space CFC_**KC**_ or CFC_**KC**^b^_, if the enzyme allocation problem, which consists in maximizing the specific flux (rate per unit of biomass amount) in a given reaction k, i.e., **v**_k_/E_T_, where **v** is a flux vector in CFC_**KC**_ or CFC_**KC**^b^_ and 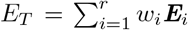 is the total protein content in the system (the w_i_’s being fixed positive weights), has an optimal solution (which is always the case for CFC_KC^b^_ if the problem is solvable), then this solution is reached in particular at an* EFM *of CFC_**KC**_ or CFC_**KC**^b^_*.

#### 2.2.3 Application to regulatory constraints

From Lemma 2.2, we obtain that *CFC*_**RC**_*Bc*__ (resp., *CFP*_**RC**_*Bc*__) is the disjoint union of 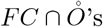 (resp., 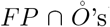), for certain open orthants 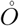 and, after merging, the disjoint union of *FC* ∩ *O*°’s (resp., *FP* ∩ *O*°’s), for certain semi-open orthants *O*° (orthants without some of their faces), without any further merging possible.

We will call open polyhedral cone (resp., open polyhedron) in dimension *r* > 0 an *r*-polyhedral cone *C* (resp., *r*-polyhedron *P*) without its facets, and we will note it 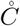 (resp., 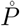) as it coincides with the topological interior of *C* (resp., *P*) in the affine span of *C* (resp., *P*). Reciprocally, *C* (resp., *P*) is the topological closure of 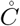 (resp., 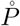) in 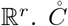 (resp., 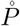) is defined as in (7) (resp., in (25)) but with strict inequalities (and equalities for defining its affine space). We will in particular consider open faces of a cone *C* or polyhedron *P* as 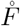 for *F* a face of *C* or *P*. *C* (resp., *P*) is the disjoint union of its open faces (by convention, a face of dimension 0, i.e., a vertex, is equal to its open face).

It follows from these definitions that, for any closed *r*-orthant *O* and any open orthant 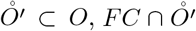 (resp., 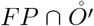) is an open face of *FC* ∩ *O* (resp., a disjoint union of open faces of *FP* ∩ *O*, as we have to keep faces corresponding to **Gv** = **h**). In all, we obtain that *CFC*_**RC**_*Bc*__ ∩ *O* (resp., *CFP*_**RC**_*Bc*__ ∩ *O*) is a disjoint union of open faces of the flux cone *FC* ∩ *O* (resp., the flux polyhedron *FP* ∩ *O*). Or, equivalently, that, for any flux tope *FC*_≤**η**_ for *FC* (resp., *FP*_≤**η**_ for *FP*), *CFC*_**RC**_**Bc**_ ≤ **η**_ (resp., *CFP*_**RC**_*Bc*_ ≤ **η**_) is a disjoint union of open faces of *FC*_≤**η**_ (resp., *FP*_≤**η**_). As we grouped together open orthants into semi-open orthants in Lemma 2.2, we can also group together with such an open face 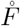 all those other open faces 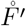 in question where *F*′ is a face of *F* to obtain thus a (minimal) disjoint union of semi-open polyhedral cones (resp., semi-open polyhedra). Here, we call semi-open polyhedral cone *C*° (resp., semi-open polyhedron *P*°) a polyhedral cone *C* (resp., polyhedron *P*) without certain (between zero and all) of its faces of lesser dimension, that can be thus expressed as a disjoint union of certain (between all and only 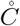, resp., 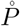) of the open faces of *C* (resp., *P*). We have: 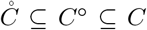 (resp., 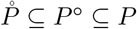).

##### Proposition 2.10.

*Given an arbitrary Boolean constraint Bc, the solution space CFC_**RC**_Bc__ (resp., CFP_**RC**_Bc__) for the regulatory constraint **RC**_Bc_ is a finite disjoint union of open, polyhedral cones (resp., open polyhedra), which are certain open faces of all the flux topes FC_≤**η**_ (resp., FP_≤**η**_). Grouping together certain of these open faces according to Lemma 2.2, we obtain a disjoint union of semi-open (i.e., without certain of their faces of lesser dimension) faces of the FC_≤**η**_’s (resp., FP_≤**η**_’s), without any possible further merging of two of them to make up a bigger semi-open face*.

Note that the rules presented in the proof of Lemma 2.2 to group together open faces into semi-open faces are applied globally to all flux topes. If we choose to apply them separately for each flux tope, then the union of semi-open faces obtained is no longer disjoint in general as two such semi-open faces for two different flux topes may have a nonempty intersection (as *CFC*_**RC**_*Bc*__ ∩ *O* (resp., *CFP*_**RC**_*Bc*__ ∩ *O*) and *CFC*_**RC**_*Bc*__ ∩ *O*′ (resp., *CFP*_**RC**_*Bc*__ ∩ *O*′), for two different closed *r*-orthants *O* and *O*′ may have open faces in common). Note also that, even after this merging, there may exist a strict inclusion relationship between the closures of two semi-open faces.

From this geometrical structure of the solution space, we will now deduce what its EFMs and its EFVs (or EFPs) are. Let’s begin with a preliminary remark. We know that, by definition of EFVs, we have, for a flux cone 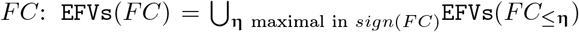 (and idem for a flux polyhedron *FP* with EFPs and EFVs), which remains true for any constrained flux cone subset *CFC*_**C**_ (or constrained flux polyhedron subset *CFP*_**C**_) whatever the biological constraint **C** is. We saw that this equality was also satisfied by EFMs for 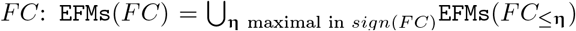 (and idem for *FP*) and remained true for *CFC*_**C**_ for **C** any thermodynamic constraint or any kinetic constraint of the form **KC_M_, KC** or **KC**^*b*^ (in the absence of bounds on enzyme concentrations), thus allowing in these cases to decompose the identification of EFMs flux tope by flux tope, as for EFVs. However this is no longer true for regulatory constraints. Actually, counter-examples can be found in the presence of reversible reactions, used in a given direction in an EFM local to a certain flux tope and in the other direction in an EFM local to another flux tope, choosing the Boolean constraint such that it imposes a strict set inclusion between the supports of these two local EFMs.

##### Example 2.1.

Consider the simple network comprising one reaction *R*: *A* → *B*, where *A* and *B* are the two internal metabolites, and four transfer reactions *T*1: → *A, T*2: → *B, T*3: *A* →, *T*4: *B* →, and assume that *R* and *T*1 are reversible (we will note par *rev*() the reversed reaction), the three other reactions being irreversible. It is obvious that **e**^1^ = (1,1,0,0,1)^*T*^, made up of *T*1, *R*, *T*4, **e**^4^ = (−1, −1, 1, 0, 0)^*T*^, made up of *T*2, *rev*(*R*), *rev*(*T*1), and **e**^3^ = (0, 0, 1, 0, 1)^*T*^, made up of *T*2, *T*4, are EFMs of *FC*. Consider now the Boolean constraint *Bc* = *T*1 ∧ *T*4. Then, **e**^1^ is an EFM of *CFC*_**RC**_*Bc*__, belonging to *CFC*_**RC**_*Bc* ≤(+,+,+,+,+)^*T*^__. And any **e** = λ**e**^3^ + **e**^4^ with λ > 0 is an EFM of *CFC*_**RC**_*Bc* ≤(−,−,+,+,+)^*T*^__ (obviously, the sub-pathways **e**^3^ and **e**^4^ of **e** do not belong to *CFC*_**RC**_*Bc*__). As *supp*(**e**^1^) = {*R,T*1,*T*4} and *supp*(**e**) = {*R,T*1,*T*2,*T*4}, we have *supp*(**e**^1^) ⊂ *supp*(**e**) and thus **e** is not an EFM of *CFC*_**RC**_*Bc*__.

Now, from a biological point of view, it is not relevant to compare supports of two pathways with a certain reaction in a given direction in the first support and in the other direction in the second support (case of *R* and *T*1 in the example). This means that the useful concept concerning minimality is not support-minimality, but sign-minimality (exactly in the same way as, concerning decomposition, we saw that the useful concept is not non-decomposability but conform non-decomposability), which is equivalent to comparing supports separately for each closed *r*-orthant, i.e., for each flux tope. We will thus identify the EFMs flux tope by flux tope (note that this is analog to distinguishing a positive flux from a negative flux in a regulatory constraint, e.g., distinguishing the constraints *T*1^+^ ∧ *T*4 and *T*1^−^ ∧ *T*4 in the example above, which could be done by adopting a tri-valued logic instead of a Boolean logic to represent these constraints; this is obviously done automatically when splitting each reversible reaction into two irreversible ones, where only the positive *r*-orthant has to be considered).

So, we will consider in the following an arbitrary closed *r*-orthant *O* given by a full support sign vector **η** (with **η**_*i*_ = + for *i* ∈ **Irr**) and consider the part of the solution space limited to this orthant, i.e., *CFC*_**RC**_*Bc*≤**η**__ (resp., *CFP*_**RC**_*Bc*≤**η**__), thus we can limit ourselves to sign vectors **η** that are maximal in *sign*(*CFC*_**RC**_*Bc*__) (resp., *sign*(*CFP*_**RC**_*Bc*__)). We saw that we could rewrite the Boolean constraint as a disjunction of two by two exclusive disjuncts, 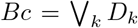, decomposing thus the solution space *CFC*_**RC**_*Bc* ≤**η**__ (resp., *CFP*_**RC**_*Bc* ≤**η**__) into the disjoint solution spaces for each disjunct, 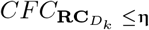 (resp., 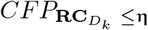). The elementary flux vectors (i.e., faces of dimension one) of *FC*_≤**η**_ that satisfy the constraint **RC***_D_k__* are obviously elementary flux vectors of 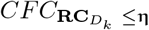 and the reciprocal is also true: if a flux vector of the semi-open polyhedral cone 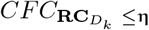 is not an elementary flux vector of *FC*_≤**η**_, i.e., is not a face of dimension one, then it belongs to the interior of a face of 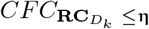 of dimension at least two and is thus conformally decomposable in this face, i.e., is not elementary in 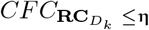. It follows that the elementary flux vectors of *CFC*_**RC**_*Bc* ≤ **η**__ are made up of all the elementary flux vectors of the *CFC*_**RC**_*Bc* ≤ **η**__’s. We can sum up the results regarding EFVs as:

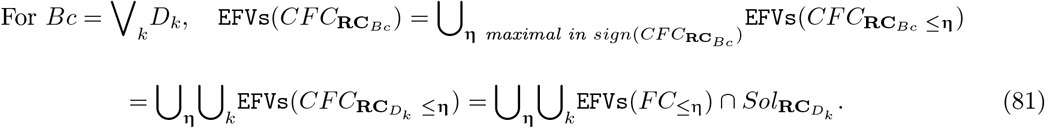

This means that EFVs can be computed flux tope by flux tope and constraint-disjunct by constraint-disjunct. And the result holds also for EFPs (by considering vertices of *FP*_≤**η**_) and EFVs of *CFP*_**RC**_*Bc*__. However, for EFMs, we have to take care that a phenomenon similar to that described in the example above still arises and that, even in a given flux tope, an EFM of 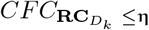 is not necessarily an EFM of *CFC*_**RC**_*Bc* ≤**η**__.

##### Example 2.2.

Consider the network comprising three irreversible reactions and one internal metabolite *A: R*1: → *A, R*2: → *A, R*3: *A* →, and the Boolean constraint *Bc* = ¬ *R*1 ∨ *R*2, decomposed as *Bc* = *D*_1_ ∨ *D*_2_, with *D*_1_ = ¬*R*1 and *D*_2_ = *R*1 ∧ *R*2. Take **η** = +. Then **e**^1^ = (0,1,1)^*T*^, made up of *R*2, *R*3, which is the only EFM of 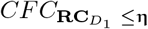, is also the only EFM of *CFC*_**RC**_*Bc* ≤**η**__. However any **e**^2^ = (λ, 1,1 + λ)^*T*^ with λ > 0, made up of *R*1, *R*2, *R*3, is an EFM of 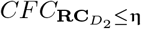 and is not an EFM of *CFC*_**RC**_*Bc* ≤ **η**__. Note that the way the constraint is decomposed matters. With the decomposition 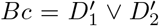 with 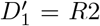 and 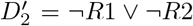, the result for 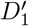 is identical to that for *D*_1_, but 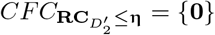.

This means that, if it is natural to study each 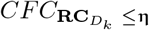 (resp., 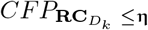) separately in order to characterize the solution space and if EFVs are obtained in this way by collecting all the local EFVs, it is not the case for EFMs and, after collecting all local EFMs, we must only keep those with minimal support:

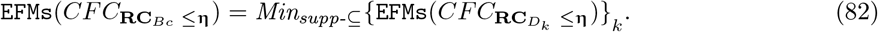

##### Proposition 2.11.

*Given an arbitrary regulatory constraint **RC**_Bc_ with 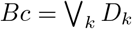, where the disjuncts D_k_ are taken two by two inconsistent, and the associated constrained flux cone subset CFC_**RC**_Bc__ (resp., constrained flux polyhedron subset CFP_**RC**_Bc__), its EFVs (resp*., EFPs *and* EFVs*) are obtained by collecting these for each flux tope (i.e., in each r-orthant O*_≤**η**_*) and for each disjunct, i.e., for each* 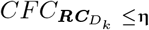 *(resp*., 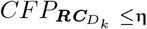*), and are nothing else than the* EFVs *(resp*., EFPs *and* EFVs*) of FC (resp., FP) satisfying the constraint **RC**_**Bc**_. This is not the case for* EFMs. *First, an* EFM *of CFC*_***RC***_*Bc* ≤**η**__ *is not necessarily an* EFM *of CFC_**RC**_Bc__, but actually the biologically relevant minimality concept being sign-minimality and not support-minimality, we will consider* EFMs *for each flux tope CFC_**RC**_Bc ≤**η**__. Second, an* EFM *of* 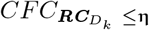 *is not necessarily an* EFM *of CFC*_***RC***_*Bc* ≤**η**__. EFMs *of CFC*_***RC***_*Bc*_ ≤**η**_ *are actually obtained by collecting* EFMs *of* 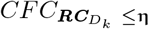 *for all k and keeping only those with minimal support*.

This being said, we will now focus on an arbitrary disjunct *D* of the form 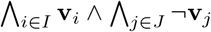 with *I, J* ⊆ {1,…, *r*}, 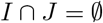. Thus 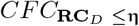 (resp., 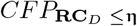) is the semi-open face *F*° of *FC*_≤**η**_ (resp., *FP_≤**η**_*), obtained from the face *F* defined by {**v**_*j*_ = 0, *j* ∈ *J*} (i.e., 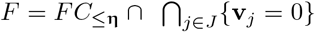, idem with *FP*) by removing its facets {**v**_*i*_ = 0} for all *i* ∈ *I*. Let’s note 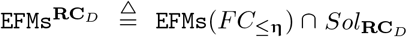 those EFMs of *FC*_≤**η**_ that satisfy the constraint **RC**_*D*_ and 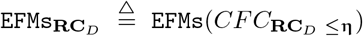 the EFMs of the part of the solution space in *O*_**η**_. Obviously, 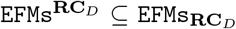. If 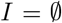, *F*° = *F* and 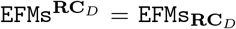, so we will consider the case 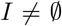. In this case, and contrary to what happens for thermodynamic and kinetic (as described in Proposition 2.8) constraints, there is generally no longer identity between 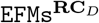 and 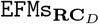 [28].

##### Example 2.3.

Consider the network of Example 2.1 (thus *D* = *T*1 ∧ *T*4) and let **η** = + and **e**^2^ = (0,1, 0,1,0)^*T*^ be the EFM of *FC*_≤**η**_ made up of *T*1 and *T*3. Then, for any λ > 0, **e**^2^ + λ**e**^3^, positive conformal combination of the two EFMs **e**^2^ and **e**^3^ of *FC*_≤**η**_, has support {*T*1,*T*2,*T*3,*T*4} and belongs to 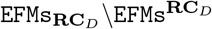.

EFMs^**RC**_*D*_^ correspond to the faces of dimension one (edges or extreme rays), if any, of the semi-open face *F*°, i.e., the edges of the face *F* that have not been removed, which means that they are not included in any hyperplane of equation **v**_*i*_ = 0, for a certain *i* ∈ *I*, or equivalently that their supports contain *I*. Now consider a face *F*′ of *F*, of dimension at least two, such that 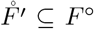 but no facet of *F*′ has its interior included in *F*° (i.e., any facet of *F*′ is included in an hyperplane of equation **v**_*i*_ = 0, for a certain *i* ∈ *I*), if any. Note that several such *F*′ may exist, but none can be included in another one, i.e., they are not comparable for inclusion in the lattice of the faces of *F*. Then all vectors of 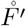 have the same minimal support, i.e., 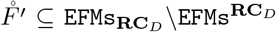. If {**e**^*k*^}_*k∈K*_ (with |*K*| ≥ 2) are representatives of the extreme vectors of *F*′, we have *F*′ = *cone*_⊕_({**e**^*k*^}) (which is actually the same as *cone*({**e**^*k*^}) as we are in orthant *O*_≤**η**_) and thus 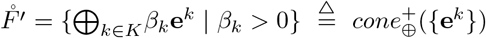 and the common minimal support of all vectors of 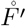 is 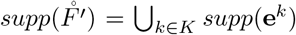. Conversely, if 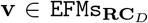, let *F*′ be the minimal face of *F* containing **v**. If *F*′ has dimension one (extreme ray), then *F*′\{**0**} ⊆ *F*° and 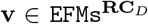. If *F*′ has dimension at least two, then no facet of *F*′ has its interior included in *F*°, because if it were the case for one facet, then any vector of its interior would have its support strictly included in the support of **v**, which would contradict the minimality of the latter. Finally, 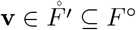 and all vectors of 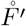 have the same support as **v** and belong to 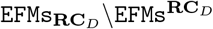. We have thus characterized both 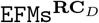 and 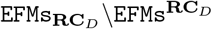. We stipulate now, for the latter one, the decomposition of its vectors into 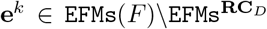, in order to compute their supports, which is generally the only useful information (the precise decomposition into fluxes not often being very relevant). So, we consider a face *F*′ of *F*, of dimension at least two, such that 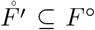 but no facet of *F*′ has its interior included in *F*° and {**e**^*k*^}_1≤*k*≤*N*_ a minimal set of vectors in EFMs(*F*′) such that 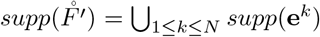. Note that, for any *k*, 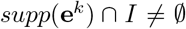 (and, as we have seen, 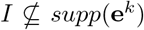), because if for a certain *k*_0_ we had 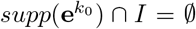, then **e**^*k*_0_^ would belong to all hyperplanes of equation **v**_*i*_ =0 for *i* ∈ *I*, thus to all facets of *F*′, which is impossible for a non-null vector. Let’s note *S*_1_ = *supp*(**e**^1^) ∪ *I* and, for any *k*, 2 ≤ *k* ≤ *N*, 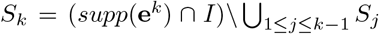. Then, for any *k*, 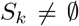, because if for a certain *k*_0_ we had 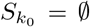, then the vectors of 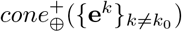 would verify the constraint **RC**_*D*_ and have their supports included in 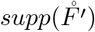, thus equal to it as it is minimal for vectors in 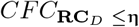, which would contradict the minimality of {**e**^*k*^}. Finally, as by construction 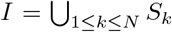 and 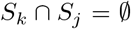 for *k* ≠ *j*, we obtain that {*S_k_*}_1≤*k*≤*N*_ constitutes a partition of *I* and {**e**^*k*^}_1≤*k*≤*N*_ is a set of vectors in 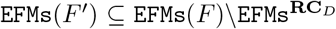 such that *supp*(**e**^*k*^) ⊅ *S_k_* by construction and 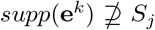 for any *j* ≠ *k*, otherwise, by the same reasoning as above, **e**^*j*^ could be suppressed from the set {**e**^*k*^}, contradicting its minimality. Finally, we obtain that the support of any vector in 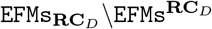 can be written as 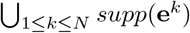, where {**e**^*k*^}_1≤*k*≤*N*_, *N* ≥ 2, are vectors in EFMs(*F*) verifying *supp*(**e**^*k*^) ⊅ *S_k_* and 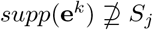 for all *k* and *j* ≠ *k*, where {*S_k_*}_1≤*k*≤*N*_ is a partition of *I* (note that we have the same result for 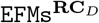 by taking *N* = 1). Now, a given 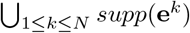 is not necessarily minimal among the whole collection when we vary *N*, {**e**^*k*^} and {*S_k_*}. It is also possible that it strictly contains the support of a vector in 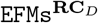. So, to obtain exactly the supports of vectors in 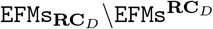, we must only keep the minimal elements for inclusion w.r.t. the whole collection extended by 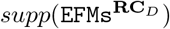.

##### Example 2.4.

Let’s continue with the network of Examples 2.1 and 2.3, so *D* = *T*1 ∧ *T*4 and **η** = +. We have EFMs(*F*) = {**e**^1^, **e**^2^, **e**^3^} and EFMs^**RC**_*D*_^ = {**e**^1^}. The only partition of {*T*1,*T*4} with a size ≥ 2 is given by: *S*_1_ = {*T*1} and *S*_2_ = {*T*4}. The only vector in EFMs(*F*) whose support contains *S*_1_ and not *S*_2_ is **e**^2^ and the only one whose support contains *S*_2_ and not *S*_1_ is **e**^3^. Thus, 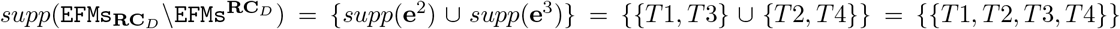. Actually, we have from Example 2.3: 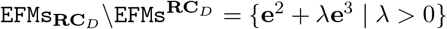.

Consider now the following network comprising two internal metabolites and seven irreversible reactions, six of which are transfer reactions, *R*: *A* → *B, T*1: → *A, T*2: *B* →, *T*3: → *A, T*4: *B* →, *T*5: *A* →, *T*6:→ *B*. Let *D* = *T*1 ∧ *T*2 and **η** = +. Let **e**^1^ = (1,1,1,0,0,0,0)^*T*^ made up of *T*1,*R, T*2, **e**^2^ = (1,0,0,1,1,0,0)^*T*^ made up of *T*3, *R, T*4, **e**^3^ = (1,1,0,0,1,0,0)^*T*^ made up of *T*1, *R, T*4, **e**^4^ = (1,0,1,1,0,0,0)^*T*^ made up of *T*3, *R, T*2, **e**^5^ = (0,1,0,0,0,1,0)^*T*^ made up of *T*1, *T*5, **e**^6^ = (0,0,0,1,0,1,0)^*T*^ made up of *T*3, *T*5, **e**^7^ = (0,0,1,0,0,0,1)^*T*^ made up of *T*6, *T*2 and **e**^8^ = (0,0,0,0,1,0,1)^*T*^ made up of *T*6, *T*4. We have EFMs(*F*) = {**e**^1^, **e**^2^, **e**^3^, **e**^4^, **e**^5^, **e**^6^, **e**^7^, **e**^8^} and EFMs^**RC**_*D*_^ = {**e**^1^}. The only partition of {*T*1,*T*2} with a size ≥ 2 is given by: *S*_1_ = {*T*1} and *S*_2_ = {*T*2}. The vectors in EFMs(*F*) whose support contains *S*_1_ and not *S*_2_ are **e**^3^ and **e**^5^ and those whose support contains *S*_2_ and not *S*_1_ are **e**^4^ and **e**^7^. We have *supp*(**e**^3^) ∪ *supp*(**e**^4^) = {*R, T*1,*T*2,*T*3,*T*4}, *supp*(**e**^3^) ∪ *supp*(**e**^7^) = {*R,T*1,*T*2,*T*4,*T*6}, *supp*(**e**^5^) ∪ *supp*(**e**^4^) = {*R,T*1,*T*2,*T*3,*T*5} and *supp*(**e**^5^) ∪ *supp*(**e**^7^) = {*T*1,*T*2,*T*5,*T*6}. Each one of the four supports obtained is minimal in this collection, but the first three contain *supp*(**e**^1^) = {*R,T*1,*T*2}. Thus, 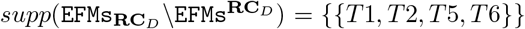. Actually, 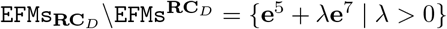.

Finally consider another network comprising three internal metabolites and seven irreversible reactions, five of which are transfer reactions, *R*1: *B* → *A, R*2: *C* → *A, T*1:→ *B* + *C, T*2: *A* →, *T*3: *A* →, *T*4:→ *B*, *T*5:→ *C*. Let *D* = *R*1 ∧ *R*2 ∧ *T*2 and **η** = +. Let **e**^1^ = (1,1,1,2,0,0,0)^*T*^ made up of *T*1, *R*1, *R*2, *T*2, **e**^2^ = (1,1,1,0,2,0,0)^*T*^ made up of *T*1, *R*1, *R*2, *T*3, **e**^3^ = (1,0,0,1,0,1,0)^*T*^ made up of *T*4, *R*1, *T*2, **e**^4^ = (1,0,0,0,1,1,0)^*T*^ made up of *T*4, *R*1, *T*3, **e**^5^ = (0,1,0,1,0,0,1)^*T*^ made up of *T*5, *R*2, *T*2 and **e**^6^ = (0,1,0,0,1,0,1)^*T*^ made up of *T*5, *R*2, *T*3. We have EFMs(*F*) = {**e**^1^, **e**^2^, **e**^3^, **e**^4^, **e**^5^, **e**^6^} and EFMs^**RC**_*D*_^ = {**e**^1^}. There are four partitions of {*R*1,*R*2,*T*2} with a size ≥ 2. The partition {{*R*1}, {*R*2}, {*T*2}} gives nothing because there is no vector in EFMs(*F*) whose support contains {*T*2} but neither {*R*1} nor {*R*2}. For the partition *S*_1_ = {*R*1,*T*2} and *S*_2_ = {*R*2}, the vector in EFMs(*F*) whose support contains *S*_1_ and not *S*_2_ is **e**^3^ and those whose support contains *S*_2_ and not *S*_1_ are **e**^2^, **e**^5^ and **e**^6^, providing supp(**e**^3^) ∪ *supp*(**e**^2^) = {*R*1, *R*2, *T*1, *T*2, *T*3, *T*4}, *sup(*(**e**^3^) ∪ *supp*(**e**^5^) = {*R*1, *R*2, *T*2, *T*4, *T*5} and *supp*(**e**^3^) ∪ *supp*(**e**^6^) = {*R*1,*R*2,*T*2,*T*3,*T*4,*T*5}. For the partition *S*_1_ = {*R*2,*T*2} and *S*_2_ = {*R*1}, the vector in EFMs(*F*) whose support contains *S*_1_ and not *S*_2_ is **e**^5^ and those whose support contains *S*_2_ and not *S*_1_ are **e**^2^, **e**^3^ and **e**^4^, providing *supp*(**e**^5^) ∪ *supp*(**e**^2^) = {*R*1,*R*2,*T*1,*T*2,*T*3,*T*5}, *supp*(**e**^5^) ∪ *supp*(**e**^3^) = {*R*1,*R*2,*T*2,*T*4,*T*5} and *supp*(**e**^5^) ∪ *supp*(**e**^4^) = {*R*1,*R*2,*T*2,*T*3,*T*4,*T*5}. For the partition *S*_1_ = {*R*1,*R*2} and *S*_2_ = {*T*2}, the vector in EFMs(*F*) whose support contains *S*_1_ and not *S*_2_ is **e**^2^ and those whose support contains *S*_2_ and not *S*_1_ are **e**^3^ and **e**^5^, providing *supp*(**e**^2^) ∪ *supp*(**e**^3^) = {*R*1,*R*2,*T*1,*T*2,*T*3,*T*4} and *supp*(**e**^2^) ∪ *supp*(**e**^5^) = {*R*1,*R*2,*T*1,*T*2,*T*3,*T*5}. The minimal elements of this collection of supports are {*R*1,*R*2,*T*1,*T*2,*T*3,*T*4}, {*R*1,*R*2,*T*1,*T*2,*T*3,*T*5} and {*R*1,*R*2,*T*2,*T*4,*T*5}. Minimizing also w.r.t. *supp*(**e**^1^) = {*R*1,*R*2,*T*1,*T*2} gives thus 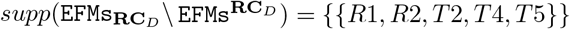. Actually, 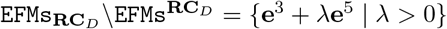.

We have thus proved the following result.

##### Proposition 2.12.

*Given a Boolean constraint D as a conjunction of literals* 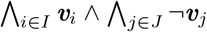 *and an arbitrary closed r-orthant O*_≤**η**_, *let CFC*_***RC**_D_* ≤**η**_ *be the constrained flux cone subset for the regulatory constraint **RC**_D_ in O*_≤**η**_. *Let F be the face of the flux cone FC*_≤**η**_ *defined as* 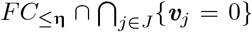, *then CFC*_***RC**_D_* ≤**η**_ *is the semi-open flux cone F° obtained from F by removing its facets* {**v**_*i*_ = 0} *for all i* ∈ *I. Let* 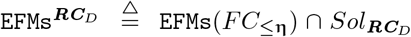 *be the* EFMs *of FC*_≤**η**_ *that satisfy **RC**_d_ and* 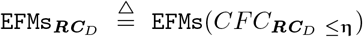 *be the* EFMs *of CFC*_***RC**_D_* ≤**η**_. *We get* EFMs^***RC**_D_*^ ⊅ EFMs_***RC**_D_*_ *but, unlike the case of thermodynamic and kinetic (as in Proposition 2.8) constraints, there is generally no longer identity between* EFMs^***RC**_D_*^ *and* EFMs_***RC**_D_*_. EFMs^***RC**_D_*^ *correspond to the extreme rays (faces of dimension one) of F°, i.e., the edges of F whose support contains I*. EFMs_***RC**_D_*_\EFMs^***RC**_D_*^ *are the vectors of the* 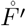’*s for all F′ faces of F of dimension at least two, such that* 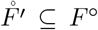 *(i.e., F′ is not included in any hyperplane* {***v**_i_* = 0} *with i* ∈ *I) but no facet of F′ has its interior included in F° (i.e., any facet of F′ is included in a certain Fyperplane* {***v**_i_* = 0 } *with i* ∈ *I). The supports of those vectors in* EFMs_***RC**_D_*_\EFMs^***RC**_D_*^ *are obtained as* 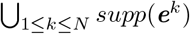, *where* {***e**^k^*}_1≤*k*≤*N*_, *N* ≥ 2, *are vectors in* EFMs(*F*) *verifying supp*(***e**^k^*) ⊇ *S_k_ and supp*(***e**^k^*) ⊉ *S_j_ for all k and j* = *k, where* {*S_k_*}_1≤*k*≤*N*_ *is a partition of I (they are actually the minimal elements, for subset inclusion and including the supports of* EFMs^***RC**_D_*^ *when checking minimality, obtained like this, for any partition* {*S_k_*} *of I, of size at least two, and any choice of* {***e**^k^*} *as above)*.

This characterization of EFMs for flux cones in the presence of regulatory constraints does not hold as simply for flux polyhedra, because the added inhomogeneous linear constraints **ILC**, given by **Gv** ≥ **h**, may not respect the structure of *CFC*_***RC**_Bc_*_ at all and may cut the interior of flux cones: we already mentioned that there is no direct relationship between EFMs and extreme or elementary fluxes for a flux polyhedron. Nevertheless, the ideas developed above for flux cones can be applied to flux polyhedra *FP*_≤**η**_, where certain of their faces have to be removed due to the constraint *D* because they are included in certain hyperplanes {**v**_*i*_ = 0}, in order to determine EFMs_***RC**_D_*_\EFMs^***RC**_D_*^. And in practice, **ILC** is generally only used to bound (below and/or above) fluxes, in which case each extremal ray 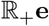 of *FC*_≤**η**_ is still partly present in *FP*_≤**η**_ as for example an edge [*α*^−^, *α*^+^]**e**, defined by the two vertices *α*^−^**e** and *α*^+^**e**. The results above can then be transposed by using convex-conformal decomposition into vertices and conformal decomposition into elementary vectors to characterize EFMs_***RC**_D_*_\EFMs^***RC**_D_*^.

We are now interested in looking for vectors in the solution space that are (in some sense) non-decomposable while not being support-minimal, and in characterizing them, if any. We could think of EFVs (or EFPs) but, from the study above regarding EFMs, we note that, for a flux cone constrained by the regulatory constraint *D*, the vectors in EFMs_***RC**_D_*_\EFMs^***RC**_D_*^ are necessarily conformally decomposable, as they can be described as the interiors of polyhedral cones in *O*_≤**η**_ of dimension at least two (for example, **e**^2^ +**e**^3^ in Example 2.3 can be decomposed as (0.7**e**^2^+0.3**e**^3^) + (0.3**e**^2^ +0.7**e**^3^)). More straightforwardly, we can notice that EFMs^***RC**_D_*^ is equal to EFVs(*CFC*_***RC**_D_* ≤**η**_), i.e., precisely the conformally non-decomposable vectors. As already pointed out, what matters more than the precise decomposition into fluxes is the decomposition of the supports (in the decomposition above of **e**^2^ + **e**^3^, the supports of the components are unchanged, i.e., equal to *supp*(**e**^2^) ∪ *supp*(**e**^3^) = {*T*1, *T*2, *T*3, *T*4}). It follows that a relevant concept for a solution vector is not to be conformally non-decomposable but, less strictly, to be (conformally) support-wise non-decomposable, in the sense that the vector cannot be conformally decomposed into two vectors of different (necessarily not greater) supports. Now, for all faces *F′* of *F* of dimension at least two such that 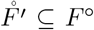, not covered by Proposition 2.12, i.e., owning at least one facet *F″* with interior included in *F*°, it so happens that all vectors of 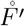 are actually support-wise decomposable, as each such vector can always be decomposed into a vector of 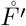 of same support and into a vector of 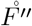 of smaller support. Therefore, in the decomposition, we do not authorize the same support for one of the component vectors. Thus the really relevant (less strict) concept is that the vector cannot be conformally decomposed into two vectors of smaller supports and we call a nonzero vector **x** of a convex polyhedral cone *C* as (conformally) support-wise non-strictly-decomposable, if:

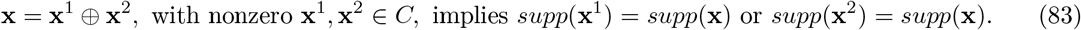

The (conformally) support-wise non-strictly-decomposable vectors in *C* (resp., flux vectors in a flux cone *FC*) will be noted swNSDVs (resp., swNSDFVs). Note that we could have defined support-wise non-strictly-decomposability more generally without imposing a conformal decomposition, but, as already pointed out, this is not relevant for biological fluxes and we will only use this concept in a certain tope. It is obvious that, in such a tope (resp., flux tope), EMs, ExVs = EVs ⊆ swNSDVs (resp., ExVs = EFVs = EFMs = swNSDFVs, so that all four definitions coincide). We will introduce a similar definition with a convex combination for a polyhedron *P* (the previous definition and the associated relationships above being valid for its recession cone *C_P_*). A vector **x** of a polyhedron *P* is called support-wise convex(- conformally) non-strictly-decomposable, if:

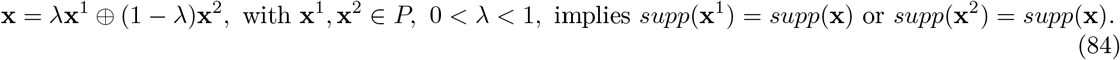

The support-wise convex(-conformally) non-strictly-decomposable vectors in *P* (resp., flux vectors in a flux polyhedron FP) will be called support-wise non-strictly-decomposable points (resp., support-wise non-strictly-decomposable flux points) and noted swNSDPs (resp., swNSDFPs). It is obvious that, in any given tope (resp., flux tope), EMs, ExPs = EPs ⊆ swNSDPs (resp., EFMs, ExPs = EFPs ⊆ swNSDFPs).

With the notations of Proposition 2.12, let 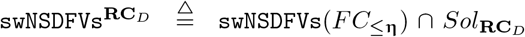 and 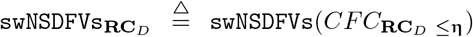. We have thus 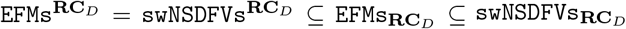 but, unlike the case of thermodynamic and kinetic (as in Proposition 2.8) constraints for flux cones, not only there is no longer identity between swNSDFVs^**RC**_*D*_^ and swNSDFVs_**RC**_*D*__ (consequence of the non-identity between EFMs^**RC**_*D*_^ and EFMs_**RC**_*D*__), but we will now see that there is generally no longer identity between EFMs_**RC**_*D*__ and swNSDFVs_**RC**_*D*__.

##### Example 2.5.

Continuing with the network of Example 2.3, **e**^1^ + **e**^3^ ∈ swNSDFVs_**RC**_*D*__ \EFMs_**RC**_*D*__. More precisely, we have (by considering only a representative for each ray) EFMs^**RC**_*D*_^ = swNSDFVs^**RC**_*D*_^ = {**e**^1^} ⊂ EFMs_**RC**_*D*__ = EFMs^**RC**_*D*_^ ∪ {**e**^2^ + λ**e**^3^ | λ > 0} ⊂ swNSDFVs_**RC**_*D*__ = EFMs_**RC**_*D*__ ∪ {**e**^1^ + λ**e**^2^ | λ > 0} ∪ {**e**^1^ + λ**e**^3^ | λ > 0}.

So, we extend the results of Proposition 2.12 by refining the structure in *F* of swNSDFVs_***RC**_D_*_. Let {**e**^*m*^}_*m*∈*M*_ be representatives of the extreme vectors of *F*, thus *F* = *cone*_⊕_({**e**^*m*^}_*m*∈*M*_). We get the following structure for *F*° regarding EFMs and swNSDFVs, given in the form of an algorithm:

- Let 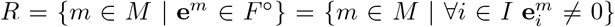. Then EFMs^**RC**_*D*_^ = {**e**^*m*^}_*m*∈*R*_. Note that *R* can vary from 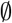 to *M*, thus EFMs^**RC**_*D*_^ from 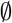 to EFMs(*F*). If *R* = *M*, then EFMs^**RC**_*D*_^ = EFMs^**RC**_*D*_^ = swNSDFVs_**RC**_*D*__ = EFMs(*F*) and the analysis is done (if **η** has been chosen maximal in *sign*(*CFC*_**RC**_*D*__), this case corresponds to 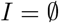). We consider the case *R* ⊂ *M* here below.
- Let’s consider successively all faces *F′* of *F* of dimension at least two, such that 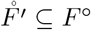, i.e., *F′* is not included in any hyperplane {**v**_*i*_ = 0} with *i* ∈ *I* (the lattice of faces of *F* can be explored for example in a way such that a sub-face is visited before a super-face; once such an *F′* has been found, all faces of *F* containing it are also suitable). Let {**e**^*k*^}_*k*∈*K*_, with *K* ⊆ *M*, be representatives of the extreme vectors of *F′*. Thus 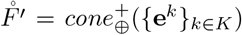 and 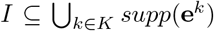. For each of these *F′*, three exclusive cases can now be distinguished.
- If no facet of *F′* has its interior included in *F*°, i.e., any facet of *F′* is included in a certain hyperplane {**v**_*i*_ = 0} with *i* ∈ *I* (a necessary but insufficient condition is: *K* ⊂ *M\R*), then 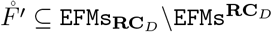 and 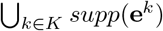 is a minimal support for vectors in *F*°.
- If exactly one facet of *F′* has its interior included in *F*°, i.e., it is the only facet not included in any hyperplane {**v**_*i*_ = 0} with *i* ∈ *I*, then 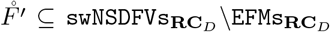 and 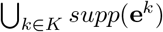 is a non-strictly-decomposable non-minimal support for vectors in *F*° (non-strictly-decomposable support means that any vector which is a conical sum of the **e**^*k*^’s having this support is support-wise non-strictly-decomposable, independently of the choice of the nonnegative coefficients fixing the contribution of each **e**^*k*^ in the distribution of the fluxes). This result follows immediately from the facts that one facet is not enough to decompose a certain vector in 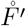 strictly in terms of supports and that the support of the vectors in the interior of the facet in question is strictly included in the support of the vectors in 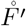.
- If at least two facets of *F′* have their interior included in *F*°, i.e., these facets are not included in any hyperplane {**v**_*i*_ = 0} with *i* ∈ *I*, then let {**e**^*l*^}_*l*∈*L*_ with *L* ⊆ *K*, be representatives of the extreme vectors of all these facets (note that we have then necessarily *K* ∩ *R* ⊆ L, i.e., *K\L* ⊂ *K\R*). Thus, the strict conical sum of the interiors of these facets, which is equal to 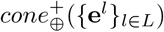, is not empty in 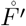 (as there are at least two such facets) and is made up of the support-wise strictly-decomposable vectors of 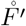 (by construction): 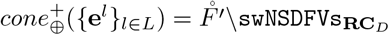. Two subcases must therfeore be distinguished.
- If *L* = *K*, i.e., the strict conical sum of the interiors of these facets is equal to 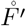, then 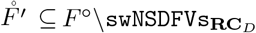 and 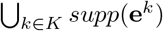 is a strictly-decomposable support for vectors in *F*° (which means that any vector which is a conical sum of the **e**^*k*^’s having this support is support-wise strictly-decomposable, independently of the choice of the nonnegative coefficients fixing the contribution of each **e**^*k*^ in the distribution of the fluxes).
- If *L* ⊂ *K*, then 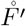 is split into two nonempty subsets: 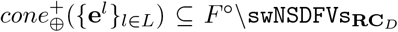 and 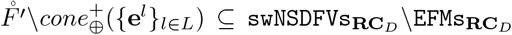 (note that 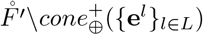) is made up of the vectors of 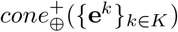 the conical decomposition of which on the **e**^*k*^’s requires at least one **e**^*k*^ with *k* ∈ *K\L*). This means that part of the vectors of 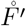 are support-wise non-strictly-decomposable and part are support-wise strictly-decomposable, while having the same non-minimal support 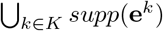 (still equal to 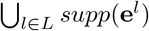). This proves that support-wise strict decomposability generally depends not only on the support but also on the positive values of the fluxes and that, unlike the particular cases above, we cannot speak of a strictly-decomposable or non-strictly-decomposable support. Note that, in this subcase, the support-wise non-strictly-decomposable vectors of 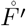 constitute the complementary, in the open cone 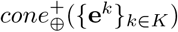, of the open subcone 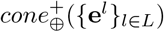 which is thus a finite disjoint union of semi-open cones, each of which is conically generated (with positive or nonnegative coefficients according to faces that are present or not) by extreme vectors **e**^*k*^’s with either *k* ∈ *K\L* ⊆ *K\R* or *k* ∈ *L\R*, thus in any case with *k* ∈ *K\R*, i.e., by extreme vectors **e**^*k*^ ∈ EFMs(*F*)\EFMs^**RC**_*D*_^.

We have finally proved the following result (keeping the notations of Proposition 2.12).

##### Proposition 2.13.

*Let* 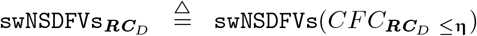 *be the support-wise non-strictly-decomposable vectors of CFC*_***RC**_D_* ≤**η**_. *We obtain* EFMs***RC**_D_* ⊆ swNSDFVs_***RC**_D_*_ *but, unlike the case of thermodynamic and kinetic (as in Proposition 2.8) constraints, there is in general no longer identity between* EFMs_***RC**_D_*_ *and* swNSDFVs_***RC**_D_*_. *Consider all faces F′ of F of dimension at least two, such that* 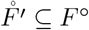 *(i.e., F′ is not included in any hyperplane* {***v**_i_* = 0} *with i* ∈ *I), and let* 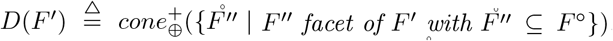. *Result of Proposition 2.12 can be stated as:* 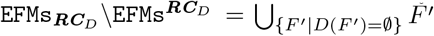. *Now we have:* 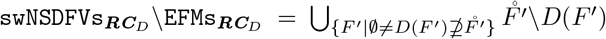. *Note that the F′’s considered here necessarily own at least one facet F″ that is not included in any hyperplane* {**v**_*i*_ = 0} *with i* ∈ *I. There are actually two cases. For those F′’s which own exactly one such facet F″ (thus* 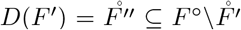*), we obtain* 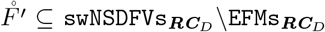 *and the common support of vectors in* 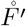 *is thus a non-strictly-decomposable (independently of the choice of the distribution of the fluxes) non-minimal support for vectors in F*°. *For those F′’s which own at least two such facets F″, we obtain* 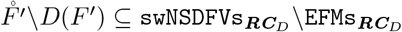 *and D*(*F*′) ⊆ *F*°\swNSDFVs_***RC**_D_*_, *thus part of the vectors of* 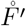 *(consisting of a finite disjoint union of semi-open cones) are support-wise non-strictly-decomposable and part (consisting of an open cone) are support-wise strictly-decomposable (depending on the choice of the distribution of the fluxes), while having the same non-minimal support*.

##### Example 2.6.

Let’s consider the simple network comprising one reaction *R*: *A* → *B*, where *A* and *B* are the two internal metabolites, and four transfer reactions *T*1: → *A*, *T*2: → *A*, *T*3: *B* →, *T*4: *B* →, and assume the five reactions irreversible. *FC* is a pointed cone of dimension 3 included in the positive orthant of 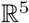 with axes {*T*1, *T*2, *T*3, *R*, *T*4} in this order. We have: *FC* = {(*x y z x* + *y x* + *y* — z)^*T*^ | *x,y,z* ≥ 0, *x* + *y* ≥ *z*}. *FC* has 4 facets and 4 extremal rays. Representatives of these extremal rays (EFMs) are: **e**^1^ = (1 0 1 1 0)^*T*^, **e**^2^ = (1 0 0 1 1)^*T*^, **e**^3^ = (0 1 1 1 0)^*T*^ and **e**^4^ = (0 1 0 1 1)^*T*^, defined by their supports: {*T*1, *R*, *T*3}, {*T*1, *R*, *T*4}, {*T*2, *R*, *T*3} and {*T*2, *R*, *T*4}, respectively.

Let’s consider the Boolean constraint *D* = *T*1 Λ *T*3. Then *F* = *FC* and the solution space is the semi-open cone *CFC*_**RC**_*D*__ = *F*° = {(*x y z x* + *y x* + *y* – *z*)^*T*^ | *x, z* > 0, *y* ≥ 0, *x* + *y* ≥ *z*}, i.e., *F* without its two facets {*x* = 0} and {*z* = 0}.

The only EFM still present in *F*° is **e**^1^, thus EFMs^**RC**_*D*_^ = {**e**^1^} with support {*T*1, *R*, *T*3}.

There are two faces of *F* of dimension two with their interiors included in 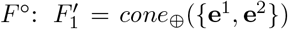 and 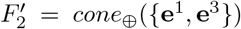, both with 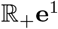 as their only facet with interior included in *F*°. Thus 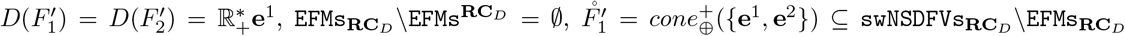 *and* 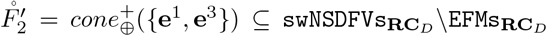. {*T*1, *R*, *T*3, *T*4} and {*T*1, *T*2, *R*, *T*3} are thus non-strictly-decomposable (independently of the respective values of the fluxes in *T*3 and *T*4 or in *T*1 and *T*2, respectively) non-minimal supports.

The last face of *F* with its interior included in *F*° is *F* itself, of dimension three, with 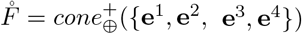 defined in *F*° by {*y* > 0, *x* + *y* > *z*}. *F* has exactly two facets with their interiors included in *F*°, namely 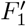 and 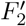 and thus 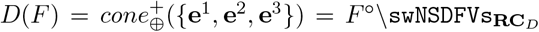. The support-wise strictly-decomposable vectors constitute the open sub-cone of 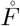 defined by {*y* < *z*}: *F*°\swNSDFVs_**RC**_*D*__ = {(*x y z x*+*y x*+*y*–*z*)^*T*^ | *x, y, z* > 0, *y* < *z* < *x*+*y*}. They are the pathways of support {*T*1, *T*2, *R*, *T*3, *T*4} such that the flux in *T*2 is smaller than the flux in *T*3. And actually any vector (*x y z x* + *y x* + *y* – *z*)^*T*^ with *x, y, z*> 0, *y<z<x* + *y* can be decomposed as: (*k*(*z* – *y*)**e**^1^ + (*x* + *y* – *z*)**e**^2^) ⊕ ((1 – *k*)(*z* – *y*)**e**^1^ + *y***e**^3^), with arbitrary *k*, 0 < *k* < 1, i.e., into two support-wise non-strictly-decomposable vectors respectively in 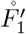 with support {*T*1, *R*, *T*3, *T*4} and in 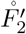 with support {*T*1, *T*2, *R*, *T*3}.

We thus have 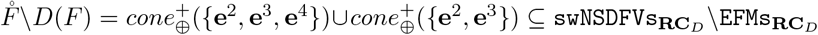. The supportwise non-strictly-decomposable vectors constitute the semi-open sub-cone of 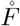 defined by {*z* ≤ *y*}: {(*x y z x* + *y x* + *y* – *z*)^*T*^ | *x, y, z* > 0, *z* ≤ *y*}. They are the pathways of support {*T*1, *T*2, *R*, *T*3, *T*4} such that the flux in *T*2 is not smaller than the flux in *T*3. Any vector (*x y z x* + *y x* + *y* – *z*)^*T*^ with *x, y, z* > 0, *z* ≤ *y* is equal to *x***e**^2^ + *z***e**^3^ + (*y* — *z*)**e**^4^ and belongs to 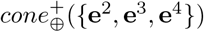 if *z* < *y* and to 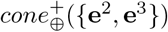 if *z* = *y*.

However take care, in the case of a general constraint 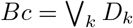, that, if the support-wise strictly-decomposable vectors of each *CFC*_**RC**_*D_k_*_ ≤ **η**_ are support-wise strictly-decomposable in *CFC*_**RC**_*B_c_*_ ≤ **η**_, it is not true for support-wise non-scrictly-decomposable vectors and some of them for *CFC*_**RC**_*D_k_*_ ≤ **η**_ may be decomposable in *CFC*_**RC**_*B_c_*_ ≤ **η**_.

This characterization of support-wise non-strictly-decomposable vectors in the presence of regulatory constraints does not extend directly from flux cones to flux polyhedra. The reason is that, due to inhomogeneous linear constraints ILC, certain faces of *FP*_≤ η_ are not defined by equalities of the form **v**_*k*_ = 0 and thus play no role in the definition of the support of the vectors they contain. Consequently, the basis of the reasoning above, namely that the support of vectors of the interior of any face is larger than the supports of vectors of the interior of any facet of this face, does not hold. Nevertheless, the ideas developed for flux cones for dealing with faces included in a certain hyperplane {**v**_*i*_ = 0} with *i* ∈ *I*, and thus removed from the solution space due to the Boolean constraint considered, can be applied. However it is necessary at each step, when considering an arbitrary face, to distinguish its facets resulting from ILC, not involved in the definition of the support, its facets included in a certain hyperplane {**v**_*k*_ = 0} with *k* ∉ *I* that are still present and contribute to the definition of the support, and its facets included in a certain hyperplane {**v**_*i*_ = 0} with *i* ∈ *I* that are removed.

#### 2.2.4 Case of several types of constraints

In general, when analyzing a metabolic pathway, all known biological constraints will have to be taken into account together, typically kinetic constraints and regulatory constraints. For two such constraints **C**_1_ (say, a kinetic constraint **KC**, equivalent to **TC**, or **KC**^*b*^ in the absence of bounds on enzyme concentrations, equivalent to **TC**^*b*^) and **C**_2_ (say a regulatory constraint **RC**_*Bc*_), the solution space, in the case of a flux cone *FC* (the reasoning would be similar for a flux polyhedron *FP*) is given by: *CFC*_**C**_1_∧**C**_2__ = *FC* ∩ *Sol*_**C**_1_∧**C**_2__ = *FC* ∩ *Sol*_**C**_1__ ∩ *Sol*_**C**_2__ = *CFC*_**C**_1__ ∩ *Sol*_**C**_2__. Now, from propositions 2.5 and 2.8, *CFC*_**C**_1__ is a finite union of flux cones *FC_i_*, which are certain particular faces of each flux tope of *FC* and the constraint **C**_2_ can be applied to each one as to an original flux cone: 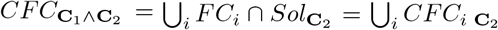. From proposition 2.10, each *CFC_i_*_**C**_2__ is a disjoint union of particular open faces of *FC_i_*. In all, the solution space is a disjoint union of particular open faces of the flux topes of *FC*. From propositions above and proposition 2.11, we can also conclude that the EFVs of *CFC*_**C**_1_∧**C**_2__ are exactly the EFVs of *FC* that satisfy both constraints **C**_1_ and **C**_2_.

##### Proposition 2.14.

*The space of flux vectors in FC (resp., FP) satisfying both the kinetic constraint **KC** (or **KC** in the absence of bounds on enzyme concentrations) and the regulatory constraint **RC**_Bc_ is a finite disjoint union of open polyhedral cones (resp., open polyhedra) which are certain open faces of the flux topes of FC (resp., FP). The elementary flux vectors (resp., elementary flux points and vectors) are those of FC (resp., FP) that satisfy both constraints*.

Note that proposition 2.13 applies also, by starting from each *FC_i_* instead of each flux tope of *FC*, to determine elementary flux modes and support-wise non-strictly-decomposable vectors.

## 3 General case of sign-compatible constraints

We are interested in determining, for general biological constraints **C**, what can be said about the structure of the solution spaces *CFC*_**C**_ or *CFP*_**C**_. More precisely, in identifying certain general features regarding constraints **C** (some of which are present in particular in biological constraints (41),(46),(49) we have considered so far), allowing us to clarify the mathematical and geometrical structure of the solution space, to determine the EFs (conformal non-decomposable fluxes) or EFMs (support-minimal fluxes) for this space and whether they characterize it and lastly how to compute them efficiently, in particular by checking the integration of this computation into the DD method. We will focus on deducing pertinent geometrical characteristics of the solution space only from general properties of the compatibility of the constraints with vector signs (i.e., with vector supports in each closed *r*-orthant or flux tope) and we will see that part of the results obtained above for thermodynamic, kinetic and regulatory constraints actually depends only on these general global properties.

### 3.1 Sign-invariant constraints

A constraint **C**_x_(**v**) (resp., ∃**xC**_x_(**v**)), is said to be sign-invariant if it depends only on *sign*(**v**), i.e., on the signs of the **v**_*i*_’s but not on their values (in the formulas below, free variables are assumed universally quantified):

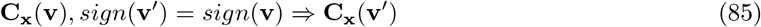

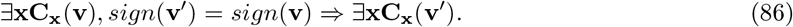

It follows from these definitions that, if the **C**_x_(**v**)’s are sign-invariant for all **x**, then ∃**xC**_x_(**v**) is sign-invariant. Note that the property for a constraint to be support-invariant (i.e., to depend only on *supp*(**v**)) is stronger than the property to be sign-invariant. Actually, in any flux tope (or in any given closed orthant), having the same sign for two vectors is equivalent to having the same support and thus being sign-invariant for a constraint means being support-invariant in each flux tope independently.

#### Example 3.1.

It follows then from definitions (40), (41) or (42) that the thermodynamic constraint **TC_M_**(**v**) or **<u>TC</u>**_<u>M</u>_(**v**) is sign-invariant. Thus this is also the case for **TC**^*b*^(**v**) and **<u>TC</u>**^*b*^(**v**) (59) and for **TC**^b^(**v**) and **TC**^b^(**v**) (60).

On the other hand, the kinetic constraint **KC_E,M_**(**v**) (46) is not sign-invariant, and this is also the case of **KC_E_**(**v**) (77), because *κ_i_*(**M**) is bounded below and above (e.g., for Michaelis-Menten kinetics, 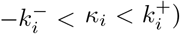 and thus, for a given nonzero enzyme concentration **E**_*i*_, there is no metabolite concentrations vector **M** allowing an arbitrary flux value **v**_*i*_ in reaction *i* (as soon as the value of **v**_*i*_/**E**_*i*_ is outside the *κ_i_* bounds).

The kinetic constraint **KC_M_**(**v**) (79) is however sign-invariant. This follows from the linear dependency of **v**_*i*_ on **E**_*i*_. Actually, if **KC_M_**(**v**) is satisfied (for a certain **E**) and *sign*(**v**′) = *sign*(**v**), then, for all *i* ∈ *supp*(**v**′), **v**_*i*_ and *κ_i_*(**M**) have the same sign, thus it is also the case for 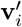 and *κ_i_*(**M**). Therefore by taking 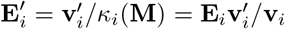 when 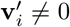, and 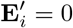 when 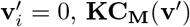 is satisfied (for **E**′). This sign-invariant property thus holds also for **KC**(**v**) (75) and for **KC**^*b*^(**v**) (76) if the only bounds are on metabolite concentrations. However it does not hold for 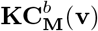 (80) and for **KC**^*b*^(**v**) (76) in the presence of bounds on enzyme concentrations.

Finally, by definition, the regulatory constraint **RC**_*Bc*_(**v**) (49) depends only on *supp*(**v**), so is support-invariant and thus sign-invariant.

#### Lemma 3.1.

*All thermodynamic constraints are sign-invariant. Only kinetic constraints **KC**_M_ and **KC** are sign-invariant, as well as **KC**^b^ with bounds only on metabolite concentrations (but not on enzyme concentrations, thus in particular without enzymatic resource constraint). The regulatory constraints are support-invariant, thus sign-invariant*.

The structure of the solution space of a sign-invariant constraint follows directly from its definition. Actually, if **v** satisfies a given sign-invariant constraint **C** = **C**_x_(**v**) (resp., **C** = ∃**xC**_x_(**v**)), then 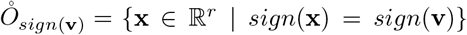 is included in *Sol*_**C**_ and thus 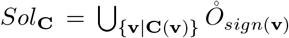 is a disjoint union of open orthants. This result can be compared to Lemma 2.2. More precisely, if we consider a support-invariant constraint **C**, the same reasoning applies and gives 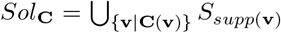, where 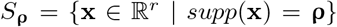 denotes the set of vectors of support **ρ**, where we code a support as a binary vector **ρ** (1 coding support membership and 0 non-membership). A support-invariant constraint **C** is thus equivalent (in extension) to a family {*S*_ρ_^*i*^}^*i*^ of such support sets and, as there are *2*^*r*^ possible binary vectors, there are thus 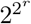 different support-invariant constraints. Now a given *S*_ρ_ is equivalent to the Boolean constraint 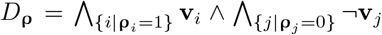 and thus an arbitrary support-invariant constraint **C** is equivalent to the Boolean constraint 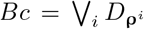, thus to the regulatory constraint **RC**_*Bc*_. Thus support-invariant constraints identify with regulatory constraints and all properties we have demonstrated for the latter apply to the first. For example, by associating with any sign vector **η** its support **ρ** (i.e., the support of any vector having the given sign), given by **ρ**_*i*_ = 1 if **η**_*i*_ = + or – and **ρ**_*i*_ = 0 if **η**_*i*_ = 0, we deduce that, for any binary vector **ρ**, there are 2^|*supp*(ρ)|^ sign vectors **η** with support **ρ**, given by **η**_*i*_ = + or – if **ρ**_*i*_ = 1 and 0 else, and we obtain 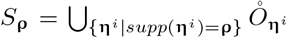 and 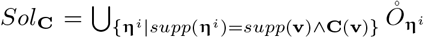, i.e., that Sol**c** is a disjoint union of open orthants, which is precisely the statement of Lemma 2.2. Note that a support-invariant constraint **C** is entirely defined by its restriction to an arbitrary closed *r*-orthant *O*_η_, i.e., for **η** an arbitrary full support sign vector, as *Sol*_C_ can be reconstituted from *Sol*_C_ ∩ *O*_η_. And, as to be sign-invariant and to be support-invariant coincide in any closed *r*-orthant, a sign-invariant constraint is nothing but a constraint whose restriction to any closed *r*-orthant is support-invariant, i.e., which coincides in any closed *r*-orthant with a well defined regulatory constraint (but such regulatory constraints differ in general from one *r*-orthant to another, while obviously coinciding on their intersection). A sign-invariant constraint **C** is equivalent to a family 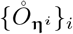 of open orthants and, as there are 3^*r*^ possible sign vectors, there are thus 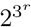 different sign-invariant constraints. Applying the merging method described in the proof of Lemma 2.2, these open orthants can be grouped together in order to obtain a family of semi-open orthants, without any possible further merging between any two of them.

#### Proposition 3.2.

*Support-invariant constraints coincide with Boolean constraints, i.e., regulatory constraints. They are completely characterized by their restriction to any closed* r*-orthant and number* 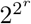. *Sign-invariant constraints coincide with constraints whose restriction to any closed r-orthant is given by a regulatory constraint (with identity of such regulatory constraints on the intersections of any two of these orthants). They number* 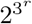 *The set Sol_C_ of vectors in* 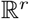 *satisfying the sign-invariant constraint **C*** = ***C***_*x*_(***v***) *(resp., **C*** = ∃***xC***_*x*_(***v***)) *is a disjoint union of open orthants, which can be grouped together according to Lemma 2.2 to provide a disjoint union of semi-open orthants without any possible further merging between any two of them*.

An important consequence is that, if we reason for each closed *r*-orthant separately, i.e., flux tope by flux tope in the solution space, then all results demonstrated in subsection 2.2.3 for regulatory constraints apply to sign-invariant constraints. With the previous definitions of open or semi-open polyhedral cones and open or semi-open polyhedra and, more generally, of open or semi-open faces of polyhedral cones or polyhedra, we get the following result.

#### Theorem 3.3.

*Let **C*** = ***C***_*x*_(***v***) *(resp., **C*** = ∃***xC***_*x*_(***v***)) *be a sign-invariant constraint and *CFC***C** (resp., CFP_C_) be the associated constrained flux cone subset (resp., the associated constrained flux polyhedron subset). Then *CFC*_C_ (resp., CFP_C_) is a finite disjoint union of open polyhedral cones (resp., open polyhedra), which are certain open faces of the flux topes *FC*_≤ η_ (resp., FP_≤ η_) for all* **η** *maximal sign vectors in sign*(*FC*) *(resp., *sign*(FP*)*). They can be grouped together according to Lemma 2.2 to provide a disjoint union of semi-open faces of the FC*_≤ **η**_’*s (resp., FP*_≤ **η**_’*s) without any possible further merging between any two of them. Elementary fluxes are obtained by collecting those for each flux tope (which is not the case for elementary flux modes where, after collecting them, only those with minimal support have to be kept) and are nothing but the elementary fluxes of *FC* (resp., FP) that satisfy the constraint **C** (which again is not the case for elementary flux modes):*

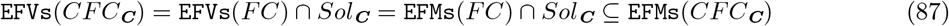

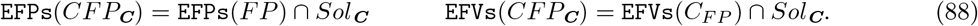

*Proof*. The disjoint decomposition of the solution space into open faces of the flux topes *FC*_≤ η_ (resp., FP_≤ η_) results from Proposition 3.2 or directly from the fact that all vectors of the strict conical hull cone^+^(**v**^1^, ⋯, **v**^*n*^) = {*β*_1_**v**^1^ + ⋯ + *β_n_***v**^*n*^ | *β*_1_, ⋯, *β_n_* > 0} and of the strict convex hull *conv*^+^(**v**^1^, ⋯, **v**^*n*^) = {*α*_1_**v**^1^ + ⋯ + *α_n_***v**^*n*^| *α*_1_, ⋯, *α_n_* > 0, *α*_1_ + ⋯ + *α_n_* = 1} of given vectors **v**^1^, ⋯, **v**^*n*^ in a flux tope, thus in a closed orthant (so the sum is conformal), have the same support *supp*(**v**^1^) ∪ ⋯ ∪ *supp*(**v**^*n*^), thus the same sign, and from the fact that an open face of a polyhedron is the Minkowski sum of the strict convex hull of its vertices and the strict conical hull of its extreme vectors. Thus an open face of a flux tope *FC*_≤ η_ (resp., *FP*_≤ η_) either does not intersect the solution space or is completely included in it. We have seen (54) that any elementary flux or any elementary flux mode of *FC* (resp., *FP*) that satisfies the constraint is an elementary flux or an elementary flux mode of the solution space (the conditions of validity of (54) for elementary fluxes are satisfied from the structure of the solution space). These elementary fluxes or elementary flux modes actually identify with the extreme vectors of the *FC*_≤ η_’s (resp., extreme points of the *FP*_≤ η_’s and extreme vectors of the *C_FP_*_≤ η_’s) that satisfy the constraint, i.e., whose sign satisfies the constraint. Reciprocally, any vector of the solution space that is not in this case necessarily belongs to an open face of a certain *FC*_≤ η_ (or *C_FP_* ≤_η_) of dimension at least two (resp., of a certain *FP*_≤ η_ of dimension at least one) and is thus conformally (resp., convex-conformally) decomposable in this open face and consequently cannot be an elementary vector (resp., elementary point) of the solution space (obviously, the decomposition involves two vectors of same support, thus nothing can be deduced regarding elementary flux modes). This gives the result for elementary fluxes.

Theorem 3.3 applies in particular to all thermodynamic constraints, to regulatory constraints and to those kinetic constraints described in Lemma 3.1. Especially, we directly obtain the results of Propositions 2.10 and 2.11 for regulatory constraints.

It is important to note that our only knowledge of the EFVs for *CFC*_**C**_ (resp., the EFPs and EFVs for *CFP*_**C**_) is really a long way from characterizing the solution space *CFC*_**C**_ (resp., *CFP*_**C**_): actually they are just the ExVs of each tope *CFC*_**C** ≤ η_ (resp., the ExPs and ExVs of each *CFP*_**C** ≤ η_), i.e., the only one-dimension open polyhedral cones, i.e., edges (resp., zero-dimension polyhedra, i.e., vertices, and edges for the recession cones) among all the open cones (resp., open polyhedra) of any dimension that constitute *CFC*_**C**_ (resp., *CFP*_C_).

As in each closed *r*-orthant, i.e., in each flux tope, a sign-invariant constraint identifies with a regulatory constraint, Propositions 2.12 and 2.13 demonstrated for regulatory constraints apply thus to sign-invariant constraints. Nevertheless, they are stated for constraints that are conjunctions of literals and, even if a decomposition into such disjuncts is always possible from the identity above, it is not necessarily natural for an arbitrary sign-invariant constraint and a global characterization for each flux tope is preferable, in particular for what concerns support-wise non-strictly-decomposable vectors. The difference is that one has to deal for *CFC*_**C** ≤ **η**_ with an arbitrary family of open faces of *FC*_≤ **η**_ instead of a single semi-open face of *FC*_≤ **η**_, obtained specifically as a face of *FC*_≤ **η**_ without certain of its facets, for *CFC*_**RC**_*D*_ ≥ **η**_.

Let us adopt notations similar to those used in Propositions 2.12 and 2.13, i.e., let 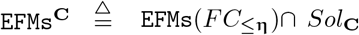 be the elementary flux modes of *FC*_≤**η**_ that satisfy **C** (i.e., the elementary flux vectors of *CFC*_**C** ≤**η**_), 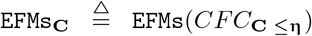 be the elementary flux modes of *CFC*_**C** ≤**η**_ and 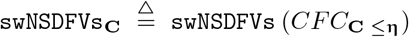 be the support-wise non-strictly-decomposable vectors of *CFC*_**C** ≤**η**_. We have thus EFMs^**C**^ ⊆ EFMs_**C**_ ⊆ swNSDFVs_**C**_. The proof of Proposition 2.12 adapts straightforwardly: EFMs^**C**^ correspond to the open faces of dimension one (extreme rays) of *FC*_≤**η**_ in *CFC*_**C** ≤**η**_ and EFMs_**C**_\EFMs^**C**^ are made up of all the open faces 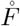 in *CFC*_**C** ≤**η**_ where *F* is a face of dimension at least two of *FC*_≤**η**_ but not any proper (i.e., with positive dimension, less than the dimension of *F*) face *F′* of *F* is such that 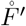 is in *CFC*_**C** ≤**η**_. The result is based on the properties that all vectors of an open face 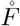 in *CFC*_**C** ≤**η**_ have the same support, and the common support of vectors of 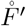, where *F′* is a face of *F*, is strictly included in that of vectors of 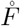. In the same way, the proof of Proposition 2.13 is easily adapted for what concerns swNSDFVs_**C**_\EFMs_**C**_. For this we consider successively all the open faces 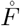 in *CFC*_**C** ≤**η**_ where *F* is a face of dimension at least two of *FC*_≤**η**_ which owns at least one proper face *F′* such that 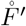 is in *CFC*_**C** ≤**η**_. For each such given 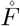, let 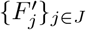 be the family of all such *F′*. Let {**e**^*k*^}_*k*∈*K*_ be representatives of the extreme vectors of *F* and {**e**^*l*^}_*l*∈*L*_, with *L* ⊆ *K*, be representatives of the extreme vectors of all these 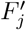. Thus 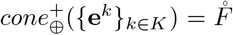 and 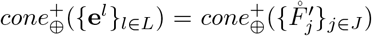 and we obtain 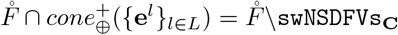, as by construction those vectors of 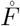 which belong to 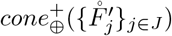 are precisely those in 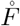 which are support-wise strictly-decomposable in *CFC*_**C** ≤**η**_, the decomposition being achieved along vectors in the 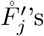’s, whose support is strictly included in the common support of vectors in 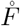. So three different exclusive cases can be distinguished for 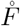:

- 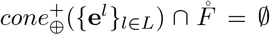, i.e., the 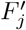’s are all included in a same facet of *F*, that is in a same hyperplane {**v**_*i*_ = 0} for a certain coordinate *i* of the affine span of *F*, which is still equivalent to 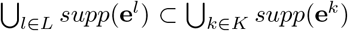 (this is for example the case if |*J*| = 1). Then 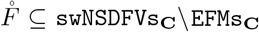 is entirely made up of support-wise non-strictly-decomposable vectors and 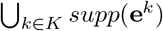 is a non-strictly-decomposable non-minimal support for vectors in *CFC*_**C** ≤**η**_.
- 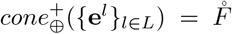, i.e., *L* = *K*. Then 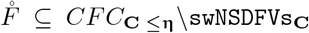 is entirely made up of support-wise strictly-decomposable vectors and 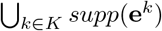 is a strictly-decomposable support for vectors in *CFC*_**C** ≤**η**_.
- 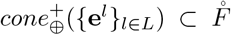, i.e., *L* ⊂ *K* and not all 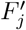’s are included in a same facet of *F*, which means that, for all *i* coordinates of the affine span of *F*, there exists *j* ∈ *J* such that 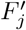 is not included in the hyperplane {**v**_*i*_ = 0}, or equivalently, there exists *l* ∈ *L* such that 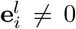, which is still equivalent to 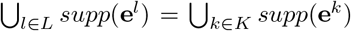. Then 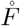 is split into two nonempty subsets: 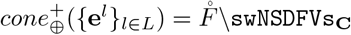, made up of support-wise strictly-decomposable vectors, and 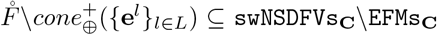, made up of support-wise non-strictly-decomposable vectors (note that 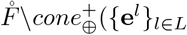) is made up of those vectors of 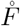 the conical decomposition of which on the **e**^*k*^’s requires at least one **e**^*k*^ with *k* ∈ *K\L*). As all vectors of 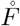 have the same non-minimal support 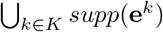, this proves that support-wise strict decomposability does not generally only depend on the support but also on the positive values of the fluxes and that, unlike the two particular cases above, we cannot speak of a strictly-decomposable or non-strictly-decomposable support. Note that the support-wise non-strictly-decomposable vectors of 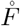 are obtained as the complementary, in this open cone, of the open sub-cone 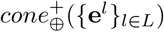, which is thus a disjoint union of semi-open cones (its connected components), each of which being conically generated both by certain extreme vectors **e**^*k*^ with *k* ∈ *K\L* (that necessarily do not belong to EFMs^**C**^), with nonnegative coefficients, and by certain extreme vectors **e**^*k*^ with *k* ∈ *L* (that may belong or not to EFMs^**C**^), with positive coefficients (in order to keep the concerned facets common with *cone*_⊕_({**e**^*l*^}_*l∈L*_)).

We have finally proved the following result (while keeping the notations of Theorem 3.3).

#### Theorem 3.4.

*For **C** a sign-invariant constraint, FC*_≤**η**_ *a flux tope and CFC*_***C*** ≤**η**_ *the associated subset of the solution space (disjoint union of open faces of FC*_≤**η**_ *from Theorem 3.3), let* 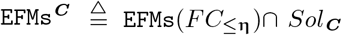 *be the elementary flux modes of FC*_≤*η*_ *that satisfy **C** (equal to* EFVs(*CFC*_***C*** ≤**η**_)), 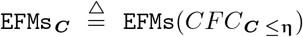 *be the elementary flux modes of CFC*_***C*** ≤**η**_, *and* 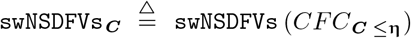 *be the support-wise non-strictly-decomposable vectors of CFC*_***C*** ≤**η**_, *with* EFMs^***C***^ ⊆ EFMs_***C***_ ⊆ swNSDFVs_***C***_. EFMs^***C***^ *correspond to the open faces of dimension one (extreme rays) of FC*_≤**η**_ *belonging to CFC*_***C*** ≤**η**_. *Consider now successively all the open faces* 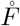 *in CFC*_***C*** ≤**η**_ *where F is a face of dimension at least two of FC*_≤**η**_, *and for each such given F, let* {**e**^*k*^}_*k*∈*K*_ *be representatives of the extreme vectors of F and* {**e**^*l*^}_*l*∈*L*_, *with L* ⊆ *K, be representatives of the extreme vectors of all proper (i.e., with positive dimension, less than the dimension of F) faces F′ of F such that* 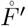 *belongs to CFC*_***C*** ≤**η**_. *Thus* 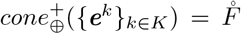 *and let* 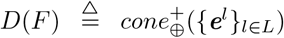. *Consequently: if* 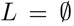, *then* 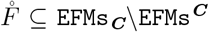*; if* 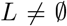 *and* 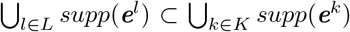, *then* 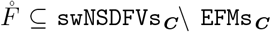 *and* 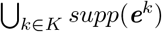 *is a non-strictly-decomposable non-minimal support for vectors in CFC*_***C*** ≤**η**_*; if L* = *K, then* 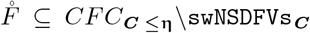 *and* 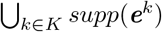 *is a strictly-decomposable support for vectors in CFC*_***C*** ≤**η**_*; if L* ⊂ *K and* 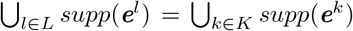, *then* 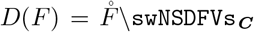, *a non-empty open cone, constitutes the support-wise strictly-decomposable vectors in* 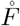 *and* 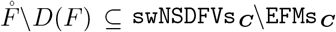, *a non-empty finite disjoint union of semi-open cones, constitutes the support-wise non-strictly-decomposable vectors in* 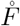, *all having the same non-minimal support (decomposability depending in this case not only on the support but also on the distribution of the fluxes). We finally obtain* EFMs_***C***_\EFMs^***C***^ *and* swNSDFVs_***C***_\EFMs_***C***_ *by collecting these vectors for each* 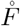.

Note that even the knowledge of swNSDFVs_**C**_ is not enough to completely reconstruct the tope *CFC*_**C** ≤**η**_ of the solution space.

In order to deal with the usual enzymatic resource constraint (more generally capacity constraints) in the kinetic constraints, we generalize the sign-invariance criterion. A constraint **C_x_**(**v**) (resp., ∃**xC_x_**(**v**)), is said to be contracting-sign-invariant if, when satisfied by one vector, it is satisfied by any vector having the same sign which is not greater (on each component), i.e., that belongs to the open rectangle parallelepiped defined by the null vector and the given vector:

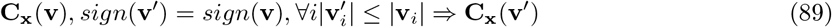

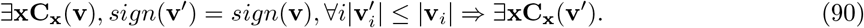

Obviously a sign-invariant constraint is contracting-sign-invariant and, if the **C_x_**(**v**)’s are contractingsign-invariant for all **x**, then ∈**xC_x_**(**v**) is contracting-sign-invariant.

#### Example 3.2.

The kinetic constraints 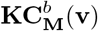 (80) and **KC**^*b*^(**v**) (76) are contracting-sign-invariant in the absence of positive lower bounds on enzyme concentrations (i.e., when **E**^-^ = **0**). Actually, if 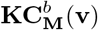 is satisfied (for a certain **E** verifying **c**^*T*^**E** ≤ *W* and **E** ≤ **E**^+^) and *sign*(**v**′) = *sign*(**v**) with 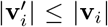 for all *i*, we saw that, by taking 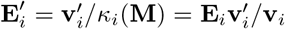 when 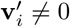, and 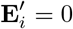 when 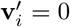, then **KC_M_**(**v**′) is satisfied (for **E**′). Now, as 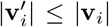, we have **E**′ ≤ **E** and thus **c**^*T*^**E**′ ≤ **c**^T^**E** ≤ *W* and **E**′ ≤ **E**^+^, so 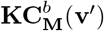 is satisfied.

#### Lemma 3.5.

*The kinetic constraints* 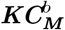 *and **KC**^b^ are contracting-sign-invariant in the absence of positive lower bounds on enzyme concentrations (i.e., when **E**^−^ = **0**)*.

We will call **0**-star domain a subset *SD* of 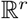 which has the property that the whole open segment joining **0** to any element of *SD* is included in *SD*: **v** ∈ *SD*, 0 < λ < 1 ⟹ *λ***v** ∈ *SD*. Any cone is a **0**-star domain.

#### Theorem 3.6.

*If **C** = **C**_x_*(***v***) *(or **C*** = ∃***xC**_x_*(***v***)*) is contracting-sign-invariant, then CFC_**C**_ is a **0**-star domain and there exists a neighborhood N* =]–*δ*, +*δ*[^*r*^ *of **0** for a certain δ >* 0 *such that CFC_**C**_* ∩ *N is a finite disjoint union of N-truncated open polyhedral cones, which are the intersection with N of open faces of the flux topes FC*_≤**η**_. *In addition, results* (87) *regarding* EFVs *and* EFMs *hold in N, i.e., for sufficiently small fluxes. The same holds for CFP_**C**_ with the flux topes FP*_≤*η*_ *if **0** is an interior point of FP, and in this case results* (88) *regarding* EFVs *and* EFPs *hold in N*.

*Proof*. The property of being a **0**-star domain is a direct consequence of the definition: if a contractingsign-invariant constraint is satisfied by a vector **v**, it is satisfied by *λ***v** for all 0 < *λ* < 1. From the proof of Theorem 3.3, it follows that if a nonzero vector **v** ∈ *FC*_≤**η**_ verifies a given contracting-sign-invariant constraint, then any vector **v**’ of the minimal open face of *FC*_≤**η**_ containing **v** (thus having the same sign as **v**) and belonging to]– *δ*_**v**_, +*δ*_**v**_[^*r*^ with *δ*_**v**_ = *min_i_*∈*supp*(**v**)|**v**_*i*_| (contracting condition) also verifies the constraint. The result is obtained by taking for *δ* the minimum of the *δ*_**v**_’s on all open faces of the *FC*_≤**η**_’s (whose number is smaller than the number of possible signs, i.e., 3^*r*^). The proof is still valid for *FP* but the result is meaningful only if *N* is included in *FP*, i.e., if **0** is an interior point of *FP*, which means that the inhomogeneous linear constraints defining *FP*, given by **Gv** ≥ **h**, verify **h** < **0**.

Theorem 3.6 tells us that the result of Theorem 3.3 for sign-invariant constraints regarding the geometrical structure of the solution space applies identically for contracting-sign-invariant constraints locally in a neighborhood of **0**, i.e., when considering only pathways with sufficiently small amounts of fluxes. This applies in particular to those kinetic constraints described in Lemma 3.5.

### 3.2 Sign-monotone constraints

We now consider a property of compatibility of a constraint with signs that is stronger than sign-invariance. A constraint **C**_x_(**v**) (resp., ∃**xC_x_**(**v**)), is said to be sign-monotone if, when satisfied by a vector, it is satisfied by any other vector with a smaller or equal sign (for the partial order on signs):

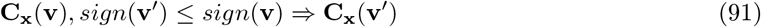

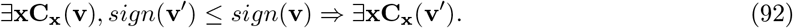

It follows from these definitions that, if the **C_x_**(**v**)’s are sign-monotone for all **x**, then ∃**xC_x_**(**v**) is sign-monotone and that any sign-monotone constraint is sign-invariant. Note that, in any given closed orthant, thus in any flux tope, *sign*(**v′**) ≤ *sign*(**v**) is equivalent to *supp*(**v′**) ⊆ *supp*(**v**) and thus being sign-monotone for a constraint means being support-monotone in each flux tope. Note also that if a sign-monotone constraint has a solution, then the null vector **0** is a solution.

#### Example 3.3.

It then follows from definitions (40), (41) or (42) that the thermodynamic constraint **TC_M_**(**v**) or <u>**TC_M_**</u>(**v**) is sign-monotone. This is thus also the case for **TC**(**v**) and <u>**TC**</u>(**v**) (59) and for **TC**^*b*^(**v**) and **TC**^*b*^(**v**) (60).

The argument given previously to establish that the kinetic constraint **KC_M_**(**v**) (79) is sign-invariant proves in fact that it is sign-monotone. This sign-monotone property holds thus also for **KC**(**v**) (75) and for **KC**^*b*^(**v**) (76) in the absence of bounds on enzyme concentrations.

The regulatory constraint **RC**_*Bc*_(**v**) (49) is not generally sign-monotone: consider for example *Bc* reduced to a positive literal. Actually, **RC**_*Bc*_(**v**) is sign-monotone if and only if *Bc* in DNF contains no positive literal, which is a very special case, requiring only certain fluxes to be zero but unable to express that a given flux is nonzero (in this particular case, it is support-monotone, which is stronger than sign-monotone).

#### Lemma 3.7.

*All thermodynamic constraints are sign-monotone. Only those kinetic constraints **KC_M_** and **KC** are sign-monotone, as well as **KC**^b^ in the absence of bounds on enzyme concentrations. The regulatory constraint **RC**_Bc_ is not sign-monotone, except if Bc in DNF contains only negative literals (i.e., assigns only zero fluxes, but no nonzero fluxes), in which case it is support-monotone*.

The structure of the solution space of a sign-monotone constraint follows directly from its definition.

#### Lemma 3.8.

*The set Sol_**C**_ of vectors in* 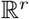 *satisfying the sign-monotone constraint **C** = **C_x_***(***v***) *(resp., **C*** = ∃***xC_x_***(***v***)*) is a union of closed orthants*.

Actually, if **v** satisfies **C**, then 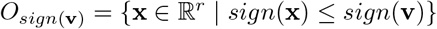 is included in *Sol*_**C**_ and thus 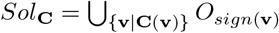. Obviously we can keep only those **v**’s for which *sign*(**v**) is maximal, in order to avoid any inclusion between the closed orthants. This result can be compared to Lemmas 2.1, 2.4 and first part of Proposition 2.8, which are obviously more precise but it shows that the mathematical structure of the solution space of these thermodynamic and kinetic constraints is mainly the only consequence of their sign-monotonicity.

#### Theorem 3.9.

*Consider indifferently a sign-monotone constraint **C_x_***(***v***) *or* ∃***xC_x_***(***v***) *(e.g., if **C**_x_*(***v***) *is sign-monotone for any **x**) and let us name it **C** and the associated constrained flux cone subset CFC_**C**_ (resp., the associated constrained flux polyhedron subset CFP_C_). Then CFC_C_ (resp., CFP_**C**_) is a finite union of polyhedral cones (resp., polyhedra), which are faces of the flux topes FC*_≤**η**_ *(resp., FP*_≤**η**_*) for all* **η** *maximal sign vectors in sign*(*FC*) *(resp., sign*(*FP*)*) and we obtain:*

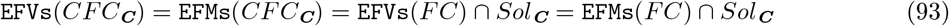

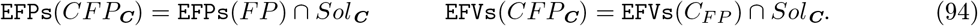

*Proof*. The (non-disjoint) decomposition of the solution space into faces of the flux topes *FC*_≤**η**_ (resp., *FP*_≤**η**_) follows from Lemma 3.8 or directly from the fact that all vectors of the conical hull *cone*(**v**^1^,…, **v**^*n*^) = {*β*_1_**v**^1^ + … + *β_n_***v**^*n*^ | *β*_1_,…, *β_n_* ≥ 0} and of the convex hull *conv*(**v**^1^,…, **v**^*n*^) = {*α*_1_**v**^1^ +…+ *α_n_***v**^*n*^ | *α*_1_,…, *α_n_* ≥ 0, *α*_1_ + … + *α_n_* = 1} of given vectors **v**^1^,…, **v**^*n*^ in a flux tope, thus in a closed orthant (so the sum is conformal), have their supports included in *supp*(**v**^1^) ∪ … ∪ *supp*(**v**^*n*^), which is the support of any vector **v** in *cone*^+^(**v**^1^,…, **v**^*n*^) or *conv*^+^(**v**^1^,…, **v**^*n*^), thus their signs being less than or equal to the sign of **v**, and from the fact that a face of a polyhedron is the Minkowski sum of the convex hull of its vertices and the conical hull of its extreme vectors. Thus, if a vector **v** of a flux tope *FC*_≤**η**_ (resp., *FP*_≤**η**_) belongs to the solution space, the minimal face of *FC*_≤**η**_ (resp., *FP*_≤**η**_) containing **v** is completely included in it. As a sign-monotone constraint is sign-invariant, the results of Theorem 3.3 apply and provide formulas for EFVs and EFPs. Thus remains only the case of EFMs. Now, if an elementary flux mode **v** of the solution space were not support-minimal in *FC*, a nonzero vector **v**’ would exist in *FC* with *supp*(**v**’) ⊂ *supp*(**v**). And we could choose 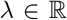 such that **v**″ = **v** – *λ***v**′ is nonzero, belongs to *FC* and verifies *supp*(**v**″) ⊂ *supp*(**v**) and *sign*(**v**″) ≤ *sign*(**v**). From the sign-monotonicity property of the constraint, **v**″ would also satisfy the constraint and would thus belong to the solution space, which would contradict the fact that **v** is an elementary flux mode in this space. So, **v** is an elementary flux mode in *FC* and we get the result for EFMs.

Of course, in the decomposition of *CFC*_**C**_(**x**) or *CFC*_**C**_ (resp., *CFP*_**C**_(**x**) or *CFP*_**C**_) as a union of certain faces of the *FC*_≤**η**_’s (resp., *FP*_≤**η**_’s), we can only keep those faces which are maximal. Theorem 3.9 applies in particular to all thermodynamic constraints, to those kinetic constraints and to those very few regulatory constraints described in Lemma 3.7. In particular, we directly obtain (except of course the reference to **ts_M_**) Proposition 2.5 for thermodynamic constraints **TC** and **TC**^*b*^, as well as the structure of the solution space as a union of flux cones (resp., flux polyhedra) for kinetic constraints **KC** and **KC**^*b*^ (in the absence of bounds on enzyme concentrations) and the characterization of elementary fluxes and elementary flux modes given by Proposition 2.8, which proves that these results depend only on the fact that these constraints are sign-monotone.

Note that the knowledge of EFVs, equal to EFMs, of C*FC*_**C**_ (resp., and of EFPs of *CFP*_**C**_) is not good enough to reconstruct the structure of the solution space *CFC*_**C**_ (resp., *CFP*_**C**_) as a union of polyhedral cones (resp., polyhedra). For this, it is necessary to know the (non-disjoint) decomposition {*E_i_*} of these EFVs (resp., and EFPs) as extreme vectors (resp., and extreme points) of the faces *F_i_* of the flux topes *FC*_≤**η**_ (resp., *FP*_≤**η**_) that constitute the solution space. The *E_i_*’s are exactly the maximal subsets of EFVs (resp., and EFPs) whose conformal conical hull (resp., conformal convex hull) is entirely contained in the solution space: *cone*_⊕_(*E_i_*) ⊆ *CFC*_**C**_ (resp., and *conv*_⊕_(*E_i_*) ⊆ *CFP*_**C**_). By analogy with LTCS, we will call each such *E_i_* a largest **C**-consistent set of EFVs or EFMs (resp., and of EFPs), noted LCS_**C**_.

In order to deal with enzymatic capacity constraints, we generalize the sign-monotonicity criterion exactly in the same way we generalized the sign-invariance criterion. A constraint **C_x_**(**v**) (resp., ∃**xC_x_**(**v**)), is said to be contracting-sign-monotone if, when satisfied by one vector, it is satisfied by any vector with a smaller or equal sign which is not greater (on each component) than the vector itself, i.e., by any vector that belongs to the rectangle parallelepiped defined by the null vector and the given vector:

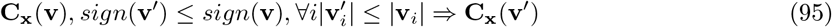

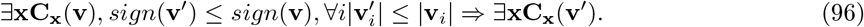

Obviously a sign-monotone constraint is contracting-sign-monotone and, if the **C_x_**(**v**)’s are contractingsign-monotone for all **x**, then ∃**xC_x_**(**v**) is contracting-sign-monotone. Also, any contracting-sign-monotone constraint is contracting-sign-invariant.

**Example 3.4.** The argument given in Example 3.2 to establish that the kinetic constraints 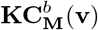 (80) and **KC**^*b*^(**v**) (76) are contracting-sign-invariant in the absence of positive lower bounds on enzyme concentrations (i.e., when **E**^−^ = **0**), proves in fact that they are contracting-sign-monotone.

#### Lemma 3.10.

*The kinetic constraints* 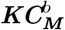 *and **KC**^b^ are contracting-sign-monotone in the absence of positive lower bounds on enzyme concentrations (i.e., when **E**^−^ = **0**)*.

#### Theorem 3.11.

*If **C** = **C_x_***(***v***) *(or **C*** = ∃**xC_x_**(***v***)*) is contracting-sign-monotone, then CFC_**C**_ is a **0**-star domain and there exists a neighborhood N* =]–*δ*, +*δ*[^*r*^ *of **0** for a certain δ* > 0 *such that CFC*_**C**_ ∩ *N is a finite union of N-truncated polyhedral cones, which are the intersection with N of faces of the flux topes FC*_≤**η**_. *In addition, results* (93) *regarding* EFVs *and* EFMs *hold in N, i.e., for sufficiently small fluxes. The same holds for CFP_**C**_ with flux topes FP*_≤**η**_ *if **0** is an interior point of FP, and in this case results* (94) *regarding* EFVs *and* EFPs *hold in N*.

*Proof*. The proof becomes identical to that of Theorem 3.6 by using the proof of Theorem 3.9.

Theorem 3.11 tells us that the result of Theorem 3.9 for sign-monotone constraints regarding the geometrical structure of the solution space and the determination of elementary fluxes and elementary flux modes applies identically for contracting-sign-monotone constraints locally in a neighborhood of **0**, i.e., when considering only pathways with sufficiently small amounts of fluxes. It applies in particular to those kinetic constraints described in Lemma 3.10, providing the result for the structure of 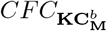 and *CFC*_**KC**^*b*^_ quoted in the last part of Proposition 2.8.

Finally, proposition 2.14 extends straightforwardly to the case where we have to deal with both one sign-monotone constraint and one sign-invariant constraint. As a sign-monotone constraint is sign-invariant the conjunction of the two constraints is sign-invariant and Theorem 3.3 applies.

#### Theorem 3.12.

*The space of flux vectors in FC (resp., FP) satisfying both a sign-monotone constraint and a sign-invariant constraint is a finite disjoint union of open polyhedral cones (resp., open polyhedra) which are certain open faces of the flux topes of FC (resp., FP). The elementary flux vectors (resp., elementary flux points and vectors) are those of FC (resp., FP) that satisfy both constraints*.

Note that, in the case of *FC*, Theorem 3.4 applies also to determine elementary flux modes and support-wise non-strictly-decomposable vectors.

### 3.3 Consequences on the computation of elementary fluxes

From the results above, we will see that the computation of elementary fluxes in the presence of a sign-monotone constraint can be efficiently performed with the Double Description (DD) method.

First let’s briefly remember the principle of the DD method [29]. This is an algorithm that takes as input the implicit description of a pointed convex polyhedral cone *C* as its representation matrix, i.e., a finite set of homogeneous linear inequalities defining *C* as the intersection of the corresponding vector half-spaces, and produces as output the explicit description of *C* as a (minimal) generating matrix, i.e., the set of extreme rays of *C*. More generally, it can deal in the same way with a pointed convex polyhedron *P* producing, from a finite set of linear inequalities defining *P* as the intersection of the corresponding affine half-spaces, the explicit description of *P* as two generating matrices providing respectively the vertices of *P* and the extreme rays of *C_P_*. As the first are obtained as the extreme rays of the pointed cone obtained from *P* by adding one dimension to the space and considering the conical hull of *P* from an origin of this extended vector space, it is enough to explain how the DD method works on pointed convex polyhedral cones. The DD method is an incremental algorithm that processes one by one each of the *n* homogeneous linear inequalities defining *C*. At each step *i*, 1 ≤ *i* ≤ *n*, it builds the intermediate extreme rays of the intermediate current cone *C_i_* defined by the *i* first linear inequalities from the knowledge of the extreme rays of *C*_*i*–1_ built at previous steps and of the *i*th linear inequality. At the end, for *i* = *n*, *C_n_* = *C* and the extreme rays are thus obtained. The *i*th linear inequality defines a half-space 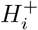 with a vector hyperplane *H_i_* as a frontier. All the extreme rays of *C*_*i*−1_ that are on the right side of *H_i_*, i.e., belong to 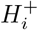, are extreme rays of *C_i_* and thus kept; all those that are on the wrong side do not belong to *C_i_* and will be discarded; the new extreme rays at step *i* appear from the intersection of *C*_*i*–1_ with *H_i_* and are obtained as the intersection with *H_i_* of the 2-faces of *C*_*i*–1_ defined by two adjacent extreme rays, one on each side of *H_i_*. We therefore just need to keep and update at each step this list of pairs of adjacent extreme rays and, when the ith linear inequality is processed, each such pair with elements on both sides of *H_i_* determines with *H_i_* an extreme ray for *C_i_*. The key fact is that each new extreme ray built at step *i* is the conical sum of two extreme rays existing at step *i* – 1 and, as we will proceed by flux tope, i.e., in a given closed *r*-orthant, this sum is conformal and the support of this new extreme ray is the union of the supports of two previously existing extreme rays. And, similarly, this new extreme ray, if involved in the next steps as a member of a relevant pair of adjacent rays, in order to build a new extreme ray, will have its support included in the support of the latter one. Finally, all we need to do is initialize *C*_1_. For a flux cone *FC* defined by (6), what has been shown as the most efficient initialization is to start from a basis of the nullspace of **S** ([45] proposed a method to compute EFMs directly as linear combinations of the vectors of a basis of this nullspace but it turned out to be less efficient than DD), and this basis (of size *r* — *m* by assuming **S** of full rank) can be chosen so that *r* – *m* of the *r* linear inequalities are satisfied (we consider here the worst case corresponding to the highest number of such inequalities, i.e., *r_I_* = *r*, which happens in particular if each reversible reaction is split into two irreversible ones). There thus remain *m* linear inequalities to satisfy, i.e., there are *n* = *m* steps in the DD algorithm.

Let us now consider a sign-monotone constraint **C**. From Theorem 3.9, the EFVs (identical to EFMs) of the constrained flux cone subset *CFC*_**C**_ are simply those of *FC*, i.e., of all its flux topes *FC*_≤**η**_ successively, that satisfy constraint **C** (and the same is true for EFPs of *CFP*_**C**_). So, we just have to build the extreme rays of each *FC*_≤**η**_ by using the DD algorithm as presented above and filter those that satisfy **C**. Obviously a filtering at the end, once all extreme rays built, will not at all improve the efficiency. However by exploiting the key fact that any newly built extreme ray at a certain step, if used in other next steps to build other new extreme rays will have necessarily its support included in each support of the latter ones, and exploiting the sign-monotonicity of **C**, we can conclude that, each time a newly built extreme ray does not satisfy the constraint **C**, it can be immediately discarded because not only it has no use at the step of its discovery but also no use at the future steps as all extreme rays, in the building of which it could be involved, would have a larger support and thus would thus not satisfy **C** either. Thus the computation of elementary fluxes or elementary flux modes in the presence of a sign-monotone constraint can be fully integrated into the incremental DD algorithm, the filtering of the extreme rays satisfying the constraint being achieved at each step when they are newly created. The extra-cost is having to check all intermediate extreme rays built for satisfiability by **C** (for most constraints, this can be practically instantaneous) and the gain is that all intermediate extreme rays for which this checking gives a negative answer are discarded.

#### Proposition 3.13.

*Given a sign-monotone constraint **C**, the* EFVs *(or* EFMs*) of the constrained flux cone subset CFC_**C**_ are obtained by the DD algorithm as the extreme rays of each flux tope FC*_≤**η**_ *by filtering out at each step all those newly built extreme rays that do not satisfy **C***.

This result applies in particular to thermodynamic constraints **TC** and **TC**^*b*^ [11, 13] and to kinetic constraints **KC** and **KC**^*b*^ (in the absence of bounds on enzyme concentrations). Actually, for **TC** (or **KC** which is identical from Proposition 2.8), Proposition 2.6 demonstrated that the criterion for an extreme ray to be thermodynamically satisfiable is to belong to a given fixed open half-space delimited by a vector hyperplane. Checking this satisfiability thus boils down to computing the scalar product of the extreme ray with a fixed vector (normal to the hyperplane and oriented towards the half-space) and verifying it is positive. In this case, as it has been shown, it is much simpler to integrate this thermodynamic constraint as one supplementary (*n* + 1)th homogeneous linear inequality and we can choose for example that it is processed first by the DD algorithm. However, as it has been already pointed out, providing the list of the extreme flux vectors of *CFC*_**C**_ does not provide its structure in terms of a union of polyhedral cones. For this, we have to identify all the LCS_**C**_’s, i.e., the maximal subsets of those extreme flux vectors whose conformal conical hull is entirely contained in *CFC*_**C**_. For each flux tope *FC*_≤**η**_, this amounts to computing all maximal upper bounds for sets of signs of those extreme rays built from this flux tope such that any vector of this tope with a sign equal to such an upper bound satisfies constraint **C**, as the vectors of the strict conical hull of a subset of extreme rays have as a common sign the upper bound for the signs of these extreme rays. To each such upper bound identified is associated the LCS_**C**_ made up of all the extreme rays whose signs are smaller than, or equal to, this upper bound.

For sign-invariant constraints, thus for regulatory constraints, no such incremental filtering is possible as a newly built intermediate extreme ray may have a sign that does not satisfy the constraint but may participate at some later stage in the construction of other new extreme rays (with necessarily larger supports) whose sign will satisfy the constraint. Thus such an extreme ray cannot be discarded. In addition we saw in any case there is no longer identity between those EFMs of *FC* that satisfy the constraint, the EFMs of *CFC*_**C**_ and the support-wise non-strictly-decomposable vectors in *CFC*_**C**_ and, as only the last two are biologically relevant, computing only the first ones would be of limited interest.

## 4 Conclusion

The analysis of metabolic networks in a steady state takes classically into account only stoichiometric and reactions irreversibility constraints. In this context, well-known pathways have been introduced, such as elementary flux modes or elementary flux vectors, which are biologically relevant in their own and from which all pathways, whose structure is a convex polyhedral flux cone *FC*, can be reconstructed. However first the computation of these EFMs or EFVs is hampered by the combinatorial explosion of their number in genome-scale metabolic models, second most of them are biologically irrelevant, because other important biological constraints are not taken into account. With the objective of both enumerating only biologically feasible EFMs or EFVs and, as there are considerably fewer of them, improving the scalability of this computation, we took in consideration in this paper one one side thermodynamic and, more generally, kinetic constraints and on the other side regulatory constraints, and we tackled the problem of revisiting in this new extended framework the concept of EFMs and EFVs and, more largely, of characterizing the geometry of the solution space. Actually, we considered a more general conceptual framework for constraints, namely their property to be compatible with vector signs (i.e., with vector supports separately in each closed *r*-orthant), because most properties of the geometrical structure of the solution space happen to depend only on this very general sign-compatibility of constraints. This is how we demonstrated, for constraints which are sign-monotone (i.e., once satisfied by a certain vector are satisfied by any vector with smaller or equal sign), which is the case of thermodynamic constraints and of kinetic constraints in the absence of bounds on enzyme concentrations, that the solution space is a union of convex polyhedral cones (which are faces of the flux topes of *FC*) and that the EFMs, which coincide with the EFVs, are simply those of *FC* that satisfy the constraint. In addition, we showed that their computation can be efficiently integrated into the classical double description algorithm, as each newly built intermediate extreme ray that does not satisfy the constraint can be filtered out at each incremental step. For the specific case of thermodynamic constraints or of kinetic constraints in the absence of bounds on enzyme concentrations, we demonstrated that their solution spaces are identical and, when there are also no bounds on metabolite concentrations, made up of those maximal faces of the flux topes of *FC* which are entirely contained in a fixed open vector half-space and thus that the EFMs are simply those of *FC* belonging to this half-space and computable simply by adding one homogeneous linear inequality to the initial (stoichiometric and reactions irreversibility) ones. The situation is more complex for constraints which are only sign-invariant (i.e., once satisfied by a certain vector are satisfied by any vector with the same sign). We demonstrated that in fact support-invariant constraints coincide with regulatory (Boolean) constraints, so, in this sense, this is not a generalization. For sign-invariant constraints (i.e., support-invariant in each closed r-orthant separately), we demonstrated that the solution space is a finite disjoint union of open (i.e., without their facets) convex polyhedral cones and that the EFVs are simply those of *FC* (equal to the EFMs of *FC*) that satisfy the constraint. However these cannot be efficiently computed by the double description algorithm because they cannot be filtered out during the incremental process. In addition it is no longer true that EFVs identify to EFMs and that EFMs identify to support-wise non-strictly-decomposable vectors: there are in general strict inclusions between these three sets and we provided again a complete characterization of the two latter ones. Finally, we extended all these results to the case where inhomogeneous linear constraints (expressing for example capacity constraints or bounds on fluxes) exist, dealing thus with a convex flux polyhedron *FP* instead of a flux cone *FC*. Basically, most of the results regarding the geometrical structure of the solution space in the presence of the above biological constraints remain the same with cones replaced by polyhedra.

Future work will be carried out along two paths. Firstly the present theoretical work will be extended to minimal cut sets (MCSs). Such MCSs are defined as minimal (for set inclusion) sets of reactions whose deletion will block the operation of given objectives or target reactions (as, e.g., those producing some toxic or undesirable product), i.e., removal of an MCS (that is, the knockout of its reactions) implies a zero flux for the target reactions in a steady state. MCSs are important for computing intervention strategies, e.g., for metabolic engineering. It is obvious from this definition that, for a metabolic network modeled in a steady state by a flux cone *FC*, the MCSs are the minimal hitting sets of the set of target EFMs (identical to EFVs) of the given metabolic network (i.e., the set of EFMs that comprise at least one of the target reactions), where a hitting set of the target EFMs is a set (of reactions) that has a nonempty intersection with each one of these EFMs. And this generalizes to a metabolic network modeled by a flux polyhedron *FP* by using EFPs. This gives an indirect method for computing MCSs [15, 18] from the preliminary computation of EFVs (or EFPs) that can be applied in the presence of biological constraints by using the results obtained in this paper. However it is known that there is also a method for computing MCSs directly as the EFVs of a dual network [4, 44], obtained basically by transposing the stoichiometric matrix of the original matrix (thus in some sense, exchanging reactions and metabolites). We will therefore study how to define this dual operation properly for metabolic networks in the presence of biological constraints in order to preserve this result (or most of it). Secondly, algorithms described in this paper will be implemented and testing conducted on metabolic networks described in the literature for which biological (thermodynamic, kinetic or regulatory) data are known. As it has been shown, the computation of elementary flux vectors (or elementary flux points) boils down to filtering those of extreme rays (or vertices) in each flux tope which satisfy the biological constraints. And for sign-monotone constraints as thermodynamic constraints or kinetic constraints in the absence of bounds on enzyme concentrations, this filtering can be easily integrated at each step of the incremental double description method, so it is natural to rely first on this method which benefits from very efficient implementations. In the absence of bounds on metabolite concentrations, the advantage is obvious as it is just necessary to add one step (i.e., one homogeneous linear constraint to deal with) in the DD algorithm. It has nevertheless been shown that at most 50% of the EFVs can be ruled out that way, which is clearly insufficient to scale up to GSMMs. Adding bounds on metabolite concentrations has already proved capable of ruling out a higher percentage of EFVs. This is achieved by checking the extreme rays at each step of the DD algorithm thanks to a call to a linear programming solver and it will be interesting to compare its efficiency with the method given by proposition 2.7 which does not need using an LP solver but has the disadvantage of requiring reasoning in a higher dimension. As the structure of the solution space cannot be deduced from merely the knowledge of the EFVs but requires the identification of the largest consistent (w.r.t. the constraints) sets of EFVs, efficient computation of these LCS_**C**_’s will be studied. We can reasonably think that scaling up will be obtained only by dealing with all biological constraints together. However, as it has been shown, handling regulatory constraints poses serious problems as these constraints are not sign-monotone and thus filtering of the EFVs that satisfy them cannot be integrated incrementally into the DD algorithm. In addition the structure of the solution space as union of open polyhedral cones (or open polyhedra) is more complex and other pathways of interest for biologists, such as elementary flux modes or support-wise non-strictly-decomposable vectors, no longer coincide with EFVs and require more complex computations to be identified. Algorithms will be carefully studied for maximal efficiency and novel ways of using the DD method or the use of other methods recently proposed such as local reverse search or satisfiability based methods [28, 26] will be investigated.

## Acknowledgments

I would like to express my sincere thanks to my colleagues Sabine Peres, who introduced me a few years ago to bioinformatics and more particularly to metabolic pathway analysis, that was a field completely unknown to me, and Antoine Deza, who explained to me all I had to know about convex geometry and the double description algorithm. Without their help and the huge number of hours we spent working and discussing together, this work could never have been accomplished. I also warmly thank Rosemary Patricot for having proofread the English.

## Notes

### Competing Interest Statement

The authors have declared no competing interest.

### Summary of Updates

Proofreading of the English and update of the refrences.

## References

[1] V. Acuña, F. Chierichetti, V. Lacroix, A. Marchetti-Spaccamela, M.-F. Sagot, and L. Stougie. Modes and cuts in metabolic networks: complexity and algorithms. BioSystems, 95:51–60, 2009.

[2] V. Acuña, A. Marchetti-Spaccamela, M.-F. Sagot, and L. Stougie. A note on the complexity of finding and enumerating elementary modes. BioSystems, 99(3):210–214, 2010.

[3] P. Atkins and J. de Paula. Physical Chemistry. Freeman, tenth edition, 2014.

[4] K. Ballerstein, A. von Kamp, S. Klamt, and U. Haus. Minimal cut sets in a metabolic network are elementary modes in a dual network. Bioinformatics, 28(3):381–387, 2012.

[5] A. P. Burgard, E. V. Nikolaev, C. H. Schilling, and C. D. Maranas. Flux coupling analysis of genome-scale metabolic network reconstructions. Genome research, 14:301–312, 2004.

[6] B. L. Clarke. Stoichiometry network analysis. Cell Biophys., 12:237–253, 1988.

[7] L. F. de Figueiredo, A. Podhorski, A. Rubio, C. Kaleta, J. E. Beasley, S. Schuster, and F. J. Planes. Computing the shortest elementary flux modes in genome-scale metabolic networks. Bioinformatics, 25(23):3158–3165, 2009.

[8] K. Fukuda and A. Prodon. Double description method revisited. In M. Deza, R. Euler, and I. Manoussakis, editors, Combinatorics and Computer Science, volume 1120 of Lecture Notes in Computer Science, pages 91–111. Springer, 1996.

[9] J. Gagneur and S. Klamt. Computation of elementary modes: a unifying framework and the new binary approach. BMC Bioinformatics, 5(175), 2004.

[10] M. P. Gerstl, C. Jungreuthmayer, S. Muller, and J. Zanghellini. Which sets of elementary flux modes form thermodynamically feasible flux distributions? FEBS Journal, 283:1782–1794, 2016.

[11] M. P. Gerstl, C. Jungreuthmayer, and J. Zanghellini. tEFMA: computing thermodynamically feasible elementary flux modes in metabolic networks. Bioinformatics, 31(13):2232–2234, 2015.

[12] M. P. Gerstl, S. Müller, G. Regensburger, and J. Zanghellini. Flux tope analysis: studying the coordination of reaction directions in metabolic networks. Bioinformatics, 35(2):266–273, 2019.

[13] M. P. Gerstl, D. E. Ruckerbauer, D. Mattanovich, C. Jungreuthmayer, and J. Zanghellini. Metabolomics integrated elementary flux mode analysis in large metabolic networks. Scientific Reports, 8930(5), 2015.

[14] S. Gudmundsson and I. Thiele. Computationally efficient flux variability analysis. BMC Bioinformatics, 11(1):489, 2010.

[15] U.-U. Haus, S. Klamt, and T. Stephen. Computing knock-out strategies in metabolic networks. Journal of Computational Biology, 15(3):259–268, 2008.

[16] C. S. Henry, L. J. Broadbelt, and V. Hatzimanikatis. Thermodynamics-based metabolic flux analysis. Biophysical journal, 92(5):1792–1805, 2007.

[17] D. Jevremovic, C. T. Trinh, F. Srienc, and D. Boley. On algebraic properties of extreme pathways in metabolic networks. Journal of Computational Biology, 17(2):107–119, 2010.

[18] C. Jungreuthmayer, G. Nair, S. Klamt, and J. Zanghellini. Comparison and improvement of algorithms for computing minimal cut sets. BMC Bioinformatics, 14:318, 2013.

[19] C. Jungreuthmayer, D. E. Ruckerbauer, M. P. Gerstl, M. Hanscho, and J. Zanghellini. Avoiding the enumeration of infeasible elementary flux modes by including transcriptional regulatory rules in the enumeration process saves computational costs. PLoS ONE, 10(6):e0129840, 2015.

[20] C. Jungreuthmayer, D. E. Ruckerbauer, and J. Zanghellini. Utilizing gene regulatory information to speed up the calculation of elementary flux modes. arXiv:1208.1853 [q-bio.MN], Aug. 2012.

[21] C. Jungreuthmayer, D. E. Ruckerbauer, and J. Zanghellini. *regEfmtool*: speeding up elementary flux mode calculation using transcriptional regulatory rules in the form of three-state logic. BioSystems, 113(1):37–39, 2013.

[22] S. Klamt, G. Regensburger, M. P. Gerstl, C. Jungreuthmayer, S. Schuster, R. Mahadevan, J. Zanghellini, and S. Müller. From elementary flux modes to elementary flux vectors: metabolic pathway analysis with arbitrary linear flux constraints. PLoS Computational Biology, 13:e1005409, Apr. 2017.

[23] A. Larhlimi, L. David, J. Selbig, and A. Bockmayr. F2c2: a fast tool for the computation of flux coupling in genome-scale metabolic networks. BMC bioinformatics, 13(57), 2012.

[24] W. Liebermeister, J. Uhlendorf, and E. Klipp. Modular rate laws for enzymatic reactions: thermodynamics, elasticities and implementation. Bioinformatics, 26(12):1528–1534, 2010.

[25] F. Llaneras and J. Picó. Which metabolic pathways generate and characterize the flux space? A comparison among elementary modes, extreme pathways and minimal generators. Journal of Biomedicine and Biotechnology, 2010:1–13, 2010.

[26] M. Mahout, R. P. Carlson, and S. Peres. Answer set programming for computing constraints-based elementary flux modes: Application to escherichia coli core metabolism. Processes, 8(12), 2020.

[27] P. McMullen. The maximum numbers of faces of a convex polytope. Mathematika, 17(2):179–184, 1970.

[28] M. Morterol, P. Dague, S. Peres, and L. Simon. Minimality of metabolic flux modes under Boolean regulation constraints. In 12th International Workshop on Constraint-Based Methods for Bioinformatics (WCB’16), Toulouse, September 2016.

[29] T. S. Motzkin, H. Raiffa, G. L. Thompson, and R. M. Thrall. The double description method. In H. W. Kuhn and A. W. Tucker, editors, Contributions to the theory of games II, Annals of Math. Studies, volume 28. Princeton University Press, 1953.

[30] S. Müller and G. Regensburger. Elementary vectors and conformal sums in polyhedral geometry and their relevance for metabolic pathway analysis. Frontiers in Genetics, 7(90), 2016.

[31] S. Müller, G. Regensburger, and R. Steuer. Enzyme allocation problems in kinetic metabolic networks: optimal solutions are elementary flux modes. Journal of Theoretical Biology, 347:182–190, 2014.

[32] E. Noor, A. Flamholz, W. Liebermeister, A. Bar-Even, and R. Milo. A note on the kinetics of enzyme action: a decomposition that highlights thermodynamic effects. FEBS Letters, 587:2772–2777, 2013.

[33] S. Peres, M. Jolicœur, C. Moulin, P. Dague, and S. Schuster. How important is thermodynamics for identifying elementary flux modes? PLoS ONE, 12(2):e0171440, 2017.

[34] S. Peres, S. Schuster, and P. Dague. Thermodynamic constraints for identifying the elementary flux modes. Biochemical Society Transactions, 46(3):641–647, 2018.

[35] R. T. Rockafellar. The elementary vectors of a subspace of *R^N^*. In Combinatorial Mathematics and its Applications (Proc. Conf., Univ. North Carolina, Chapel Hill, N.C., 1967), pages 104–127. Univ. North Carolina Press, Chapel Hill, N.C., 1969.

[36] A. Röhl, Y. Goldstein, and A. Bockmayr. EFM–Recorder – Faster elementary mode enumeration via reaction coupling order. In Advances in Systems and Synthetic Biology, pages 91–99, Strasbourg, March 2015.

[37] C. H. Schilling, D. Letscher, and B. O. Palsson. Theory for the systemic definition of metabolic pathways and their use in interpreting metabolic function from a pathway-oriented perspective. Journal of Theoretical Biology, 203:229–248, 2000.

[38] C. H. Schilling and B. O. Palsson. The underlying pathway structure of biochemical reaction networks. Proceedings of the National Academy of Sciences USA, 95:4193–4198, 1998.

[39] S. Schuster and C. Hilgetag. On elementary flux modes in biochemical reaction systems at steady state. Journal of Biological Systems, 2(2):165–182, 1994.

[40] M. Terzer and J. Stelling. Large-scale computation of elementary flux modes with bit pattern trees. Bioinformatics, 24(19):2229–2235, 2008.

[41] C. T. Trinh, A. Wlaschin, and F. Srienc. Elementary mode analysis: a useful metabolic pathway analysis tool for characterizing cellular metabolism. Applied Microbiology and Biotechnology, 81(5):813–826, 2009.

[42] R. Urbanczik and C. Wagner. An improved algorithm for stoichiometric network analysis: theory and applications. Bioinformatics, 21(7):1203–1210, 2005.

[43] J. B. van Klinken and K. Willems van Dijk. FluxModeCalculator: an efficient tool for large-scale flux mode computation. Bioinformatics, 32(8):1265–1266, 2016.

[44] A. von Kamp and S. Klamt. Enumeration of smallest intervention strategies in genome-scale metabolic networks. PLoS Computational Biology, 10(1):e1003378, 2014.

[45] C. Wagner. Nullspace approach to determine the elementary modes of chemical reaction systems. J. Phys. Chem. B, 108(7):2425–2431, 2004.

[46] C. Wagner and R. Urbanczik. The geometry of the flux cone of a metabolic network. Biophysical Journal, 89(6):3837–3845, 2005.

[47] M. T. Wortel, H. Peters, J. Hulshof, B. Teusink, and F. J. Bruggeman. Metabolic states with maximal specific rate carry flux through an elementary flux mode. FEBS Journal, 281:1547–1555, 2014.

[48] J. Zanghellini, D. E. Ruckerbauer, M. Hanscho, and C. Jungreuthmayer. Elementary flux modes in a nutshell: properties, calculation and applications. Biotechnology Journal, 8(9):1009–1016, 2013.

